# IL-6 SIGNALING EXACERBATES HALLMARKS OF CHRONIC TENDON DISEASE BY STIMULATING REPARATIVE FIBROBLASTS

**DOI:** 10.1101/2023.02.13.528273

**Authors:** Tino Stauber, Greta Moschini, Amro A. Hussien, Patrick K. Jaeger, Katrien De Bock, Jess G. Snedeker

**Affiliations:** University of Zurich, Department of Orthopedics ETH Zurich, Institute for Biomechanics Lengghalde 5, 8008 Zurich, Switzerland; Laboratory of Exercise and Health, Department of Health Sciences and Technology (D-HEST) ETH Zurich, Swiss Federal Institute of Technology

**Keywords:** tendon, interleukin-6, fibroblasts, assembloid

## Abstract

Tendinopathies are debilitating diseases currently increasing in prevalence and associated costs. There is a need to deepen our understanding of the underlying cell signaling pathways to unlock effective treatments. In this work, we screen cell signaling pathways in human tendinopathies and find positively enriched IL-6/JAK/STAT signaling alongside signatures of cell populations typically activated by IL-6 in other tissues. In human tendinopathic tendons, we also confirm the strong presence and co-localization of IL-6, IL6R, and CD90, an established marker of reparative fibroblasts. To dissect the underlying causalities, we combine IL-6 knock-out mice with an explant-based assembloid model of tendon damage to successfully connect IL-6 signaling to reparative fibroblast activation and recruitment. Vice versa, we show that these reparative fibroblasts promote the development of tendinopathy hallmarks in the damaged explant upon IL-6 activation. We conclude that IL-6 activates tendon fibroblast populations which then initiate and deteriorate tendinopathy hallmarks.

## INTRODUCTION

Tendons are essential to every human movement.^[1]^ Tendinopathies represent the largest group of common tendon diseases and approximately 22% of all sport-related injuries.^[2]^ They can strike tendons at many different anatomical locations, can dramatically diminish quality of life by limiting the associated movements, and often share a history of repetitive overuse-induced damage and repair cycles.^[2–5]^ Once adult tendon regions fail to keep up with functional demands, they fall into a state of non-resolving, uncontrolled lesion repair.^[6–12]^

These chronic tendon lesions underlying tendinopathies feature characteristics of normal wound healing including accelerated extracellular matrix turnover and proliferation of reparative fibroblast populations as well as their migration to replace and repopulate damaged tissues.^[12–17]^ In tendon, reparative (e.g. Scx^+^) fibroblast populations are assumed to reside primarily in the extrinsic compartment comprising epitenon and paratenon from where they are recruited to the damaged intrinsic compartment embodied by the load-bearing tendon core.^[15,18–21]^ While the mechanisms governing activation, proliferation, and recruitment of these populations are unclear, some insight can be gleaned from studies on other musculoskeletal tissues.^[4,22]^

In acute muscle lesions, mechanisms for repair, hypertrophy, and hyperplasia are dominated by the satellite cells residing in the muscle basal lamina.^[23–26]^ Satellite cells in muscle and dermal fibroblasts in skin are activated and recruited by interleukin-6 (IL-6), a key player in the acute phase response to stress.^[23,26,27]^ Stress-related mechanisms activating *IL-6* mRNA transcription in the absence of exogenous pathogens include damage-associated molecular patterns (DAMPs), calcium signaling after membrane-depolarization, but also energetic stressors like glycogen depletion and redox signaling following exercise.^[28,29]^ In humans, IL-6 transmits its signal via classical or trans-signaling.^[30]^ Classical signaling involves the membrane-bound receptor IL6R, which forms a homodimer with gp130 upon IL-6 binding. Trans-signaling works similarly, except that the IL6R has been solubilized (sIL6R) by metalloproteases (mostly ADAM10 & 17)^[31]^, which cleave it from the cell membrane.^[30,31]^ Since not all cell populations express IL6R, trans-signaling via sIL6R enables IL-6 signaling for a wider range of cell populations. Regardless of classical or trans-signaling initiation, further transduction of the IL-6 signal runs via two major pathways (JAK/STAT/ERK^[32,33]^ & SHP2/GAB2/MAPK^[34,35]^) to turn on cellular processes inducing proliferation, migration, metabolic adaptations, and tissue turnover.^[30,36]^ In chronic muscle lesions like Duchenne muscular dystrophy, IL-6 is persistently upregulated, and anti-IL-6 receptor antibodies have been proposed as treatment options.^[37]^ Anti-IL-6 receptor antibodies (IL-6 inhibitors) like Tocilizumab inhibit both classical and trans-signaling and are routinely used in other chronic inflammatory diseases like systemic sclerosis, psoriasis, and rheumatoid arthritis.^[28,38–42]^ In this context, IL-6 inhibitors have been shown to reduce disease-hallmarks including arthritis-concomitant tendon inflammation.^[43–45]^ Analogous to these chronic inflammatory musculoskeletal diseases, tendinopathies present with localized pain, swelling, and functional decline in the affected organ.^[4,7,46,47]^ Histological and molecular characteristics of tendinopathy include hypercellularity, disorganized collagen fibers including mechanically inferior collagen-3, and dysregulated extracellular matrix homeostasis.^[14,48–50]^ Based on this research performed in other tissues, repair-competent fibroblasts appear as prime targets and effectors for IL-6 signaling in a tendon wound healing context.^[27,51–53]^

Here, we investigated the hypothesis that IL-6 plays a vital role in activating reparative fibroblasts within the extrinsic tissue compartment (i.e. epitenon or paratenon) and recruitment of these cells to the damaged tendon core tissue in non-sheathed tendons.^[12,54–56]^ While this likely represents a critical step in normal tendon healing^[57–59]^, we propose that extended and excessive IL-6 signaling may causally exacerbate tendinopathy in non-sheathed tendons.^[60]^

We experimentally tested and confirmed these hypotheses in four steps:

1. We showed that the IL-6/JAK/STAT signaling cascade is positively enriched in (non-sheathed) human tendinopathic tendons alongside gene signatures typical for fibroblasts as well as downstream gene sets suggesting excessive cell proliferation (hypercellularity), imbalanced extracellular matrix turnover, and neo-vascularization.
2. We verified increased IL-6 and IL6R levels in tendinopathic human tissue samples with fluorescence microscopy. We also co-stained these sections with a marker for reparative fibroblasts (CD90^+^) to confirm their increased presence and gauge their contribution to the elevated IL-6 and IL6R levels in tendinopathic human tissues.
3. We exploited an explant-based assembloid model system to confirm the causal effect of IL-6 signaling on extrinsic fibroblast activation, recruitment, and proliferation in tendinopathic niche conditions.
4. We followed the downstream effects of enhanced extrinsic fibroblast activation and accumulation on the tendon core embedded in our assembloids. Here, we document the emergence of central tendinopathic hallmarks including aberrant (catabolic) matrix turnover, hypercellularity, and hypoxic responses with a stronger fibroblast presence, which is reduced when IL-6 core signaling is inhibited.

## RESULTS

### The IL-6/JAK/STAT signaling signature is positively enriched in human tendinopathic tendons alongside signatures of extrinsic cell population activation and hallmarks of clinical tendinopathy

To better illuminate the ongoing wound healing processes that are a central feature of chronic tendon disease, we first searched for enriched signaling pathways underlying tendinopathic tendons in a publicly available dataset (GEO: GSE26051).^[61]^ To deduce common disease patterns affecting tendons from diverse anatomical locations, the microarray dataset contained analyzed samples from 23 normal and 23 tendinopathic human tendons of mixed anatomical origin. We excluded 5 normal and 5 tendinopathic samples from sheathed tendons originally included in the dataset, as those tendons are organized into different sub-compartments than the non-sheathed tendons remaining in the dataset (**Supplementary Table 1 & Supplementary Figure 2**). Overall, we found 240 significantly upregulated genes, 474 significantly downregulated genes, and 20,359 not-differentially regulated genes in tendinopathic compared to normal control tendons.

Further analysis revealed significantly upregulated *IL-6* and *IL-6* signaling transducer (*IL6ST,* also known as *GP130*) transcripts in tendinopathic tendon tissue (**Figure 1A & B, Table 1**). Conversely, *IL-6* receptor (*IL6Rα*) expression trended toward downregulation. Focusing on proteases that generate soluble *IL6Rα*, we found significant upregulation of both *ADAM10* and *ADAM17* in human tendinopathic tendons. Downstream of *IL-6* receptor binding, *JAK1* trended towards upregulation and *STAT3* was upregulated significantly. Other *IL-6* regulated signaling checkpoints such as *MAPK3* and *GAB2* trended towards downregulation. Aside from *IL-6*, another significantly upregulated member of the IL-6 family was interleukin-11 (*IL-11*).

**Figure 1:**
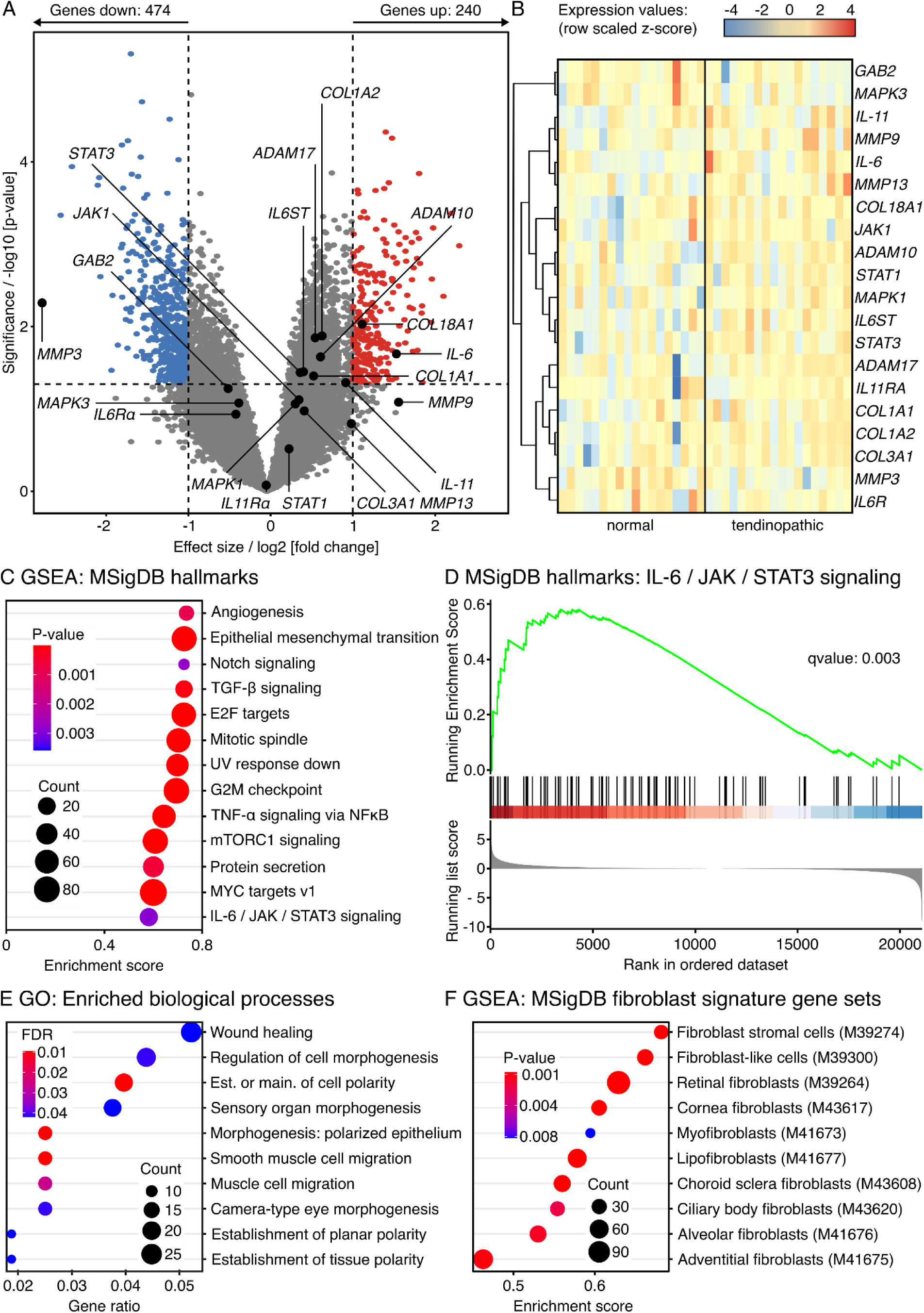
Transcriptome analysis of up- and downregulated genes and pathways in tendinopathic compared to normal control human tendons (non-sheathed) (A) Volcano plot of differentially expressed genes (DEGs) comparing tendinopathic to normal human tendons. Genes colored in red have a log2 (fold change) > 1, a p-value < 0.05, and are therefore considered to be significantly increased in tendinopathic tendons. Genes colored in blue have a log2 (fold change) < −1, a p-value < 0.05, and are therefore considered to be significantly decreased in tendinopathic tendons. The log2 and p-value thresholds are represented by the dashed lines. Annotated genes are part of the IL-6 cytokine superfamily, the IL-6 signaling cascade, or involved in matrix turnover. (B) Unsupervised hierarchical clustering of expression values from members of the IL-6 cytokine superfamily, their receptors, and parts of the IL-6 signaling cascade (N=18 normal, N=18 tendinopathic). Genes are clustered by color with positive (red) or negative (blue) row-scaled z-scores. (C) Dotplot showing significantly enriched gene sets (p-value < 0.05) as determined by GSEA based on the MSigDB human hallmark gene sets. The color of the circles represents their p-value, the size the number of enriched genes (count), and the position on the x-axis the enrichment score as well as its direction. (D) GSEA plot for the IL-6/JAK/STAT3 signaling hallmark contained in the MSigDB human hallmark gene sets. The green line traces the running enrichment score on the y-axis while going down the rank of genes listed on the x-axis, the black lines standing in blue and red bars indicate the locations of the genes related to the pathway in the ranked list, and the grey histogram shows the running list score across the ranks. (E) Dotplot showing the top 10 GO gene sets for biological processes most significantly enriched by overlapping DEG sets. The color of the circles represents their adjusted p-value (FDR), the size the number of enriched genes (count), and the position on the x-axis the number of enriched genes in ratio to the total number of genes annotated to the gene set (gene ratio). (F) Dotplot showing significantly enriched fibroblast signature gene sets (p-value < 0.05) as determined by GSEA based on the MSigDB human cell type signature gene sets. The color of the circles represents their p-value, the size the number of enriched genes (count), and the position on the x-axis the enrichment score as well as its direction.

**Table 1:** Effect sizes and p-values for selected transcripts. The data describes the differences between tendinopathic and normal control human tendons (non-sheathed).

| Transcript | Effect size | P-value |
| --- | --- | --- |
| *IL-6* | 1.529 | 0.021 |
| *IL6Ra* | -0.423 | 0.116 |
| *IL6ST / GP130* | 0.399 | 0.035 |
| *ADAM10* | 0.606 | 0.023 |
| *ADAM17* | 0.541 | 0.014 |
| *JAK1* | 0.344 | 0.078 |
| *STAT1* | 0.224 | 0.305 |
| *STAT3* | 0.36 | 0.036 |
| *MAPK1* | 0.3 | 0.086 |
| *MAPK3* | -0.386 | 0.085 |
| *GAB2* | -0.517 | 0.057 |
| *IL-11* | 0.913 | 0.048 |
| *IL11Ra* | -0.053 | 0.844 |
| *COL1A1* | 0.523 | 0.04 |
| *COL1A2* | 0.624 | 0.013 |
| *COL3A1* | 0.408 | 0.106 |
| *COL18A1* | 1.112 | 0.009 |
| *MMP9* | 1.556 | 0.083 |
| *MMP13* | 0.981 | 0.151 |
| *MMP3* | -2.779 | 0.005 |

In line with aberrant matrix turnover generally featured in tendinopathy, the transcripts of the following genes were significantly increased in the tendinopathic samples: *COL1A1*, *COL1A2, COL18A1*.

Since changes in single transcripts alone have a limited predictive value for pathway-level changes, we next performed unbiased gene set enrichment analysis (GSEA) using the human hallmark dataset from MSigDB.^[62,63]^ Confirming the trends from the single transcript analysis, GSEA revealed a positive enrichment of the IL-6/JAK/STAT pathway (qvalue: 0.003) alongside gene sets matching well known tendinopathy hallmarks such as neo-vascularization (i.e. angiogenesis, mTORC1 signaling) and hypercellularity (i.e. G2M checkpoint, mitotic spindle, MYC targets v1, epithelial mesenchymal transition, and E2F targets) in the tendinopathic samples compared to the normal controls (**Figure 1C & D**). We then looked further into aberrant biological processes by mapping the significantly changed single transcripts (p-value < 0.01) to the respective gene ontology (GO) database in an overrepresentation analysis (**Figure 1E & Supplementary Figures 3-4**). The emerging processes pointed towards ongoing morphogenesis and wound healing favoring hypercellularity (i.e. proliferation, migration) and extracellular matrix (ECM) turnover, which are both established hallmarks of tendinopathy. Lastly, we matched the detected transcript changes to the human cell type signature gene sets from MSigDB in a GSEA to estimate the contribution of fibroblasts to the aberrant processes.^[62,63]^ While this database does not yet include tendon-specific fibroblast populations, the signature gene sets of several fibroblast populations were significantly enriched by the transcript changes detected in tendinopathic tendons (**Figure 1F**). To confirm the increase of transcripts related to IL-6 signaling and fibroblast presence on the protein level, we next assessed human patient samples using fluorescence microscopy.

### IL-6, IL6R, and CD90 are elevated on the protein level in tendinopathic human tendons compared to normal control tendons

In a second step, we sought to validate the gene array analysis highlighting elevated IL-6-IL6R signaling as well as the elevated presence of fibroblasts in non-sheathed tendinopathic tendons compared to non-sheathed normal control tendons on the protein level. We thus extracted tissue sections from tendinopathic biceps tendons and from normal control tendons leftover after anterior cruciate ligament reconstruction (ACL) surgery (**Figure 2A and Supplementary Figure 5**) and stained them with fluorescently labelled IL-6, IL6R, and CD90 antibodies.

**Figure 2:**
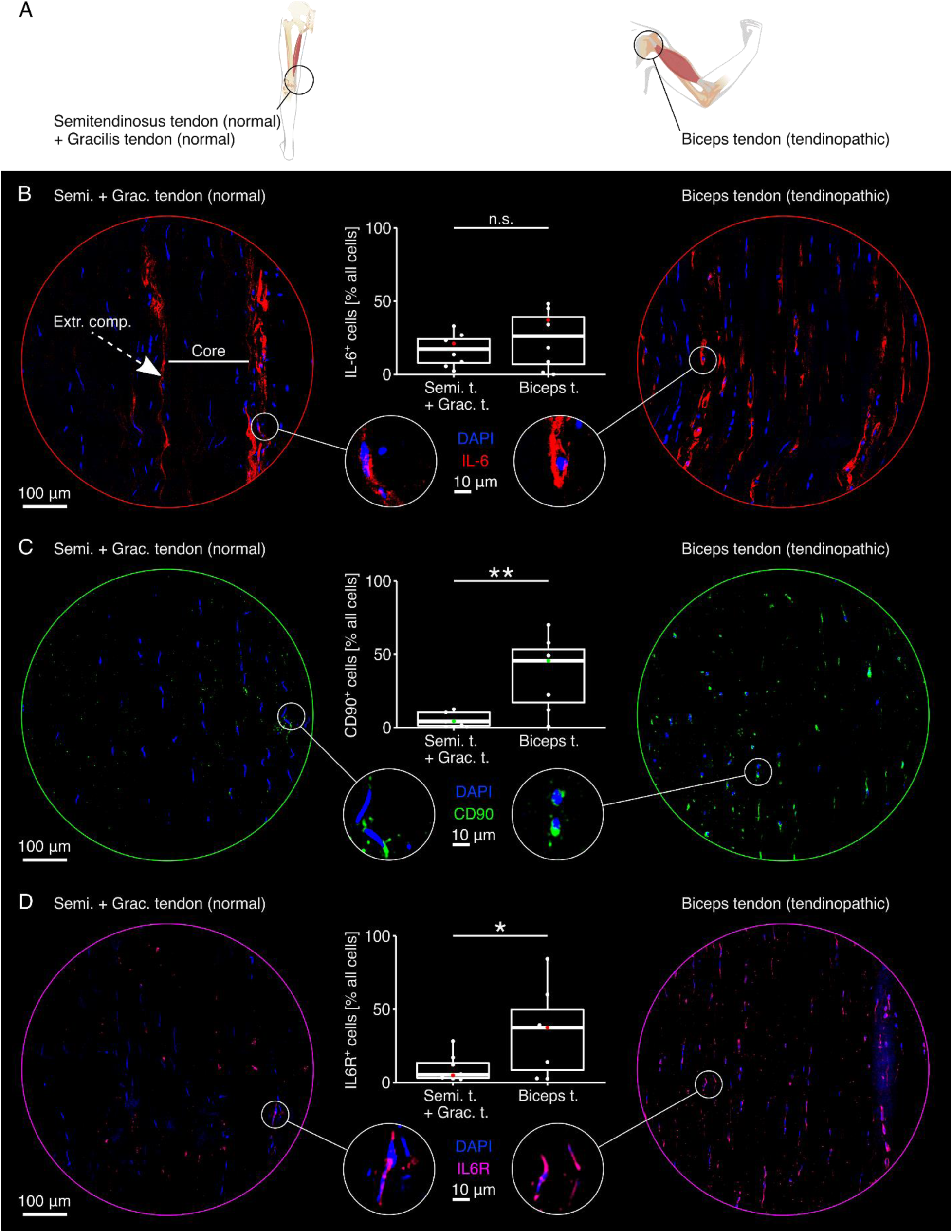
Distribution of IL-6, CD90, and IL6R in normal control and tendinopathic human tendons (non-sheathed) (A) Illustrative depiction of the origins of the tendons used in this experiment. Normal control tendon tissues were taken from semitendinosus and gracilis tendons leftover from ACL reconstruction surgery. Tendinopathic tissues were taken from painful shoulders during surgery (**Supplementary Figure 4**). (B) Representative fluorescence microscopy images of normal control (left) and tendinopathic tendons (right) stained with DAPI (blue) and an IL-6 antibody (red). Boxplots depict the quantified co-localization of DAPI and IL-6 (IL-6^+^ cells) calculated as percentage of the total number of cells. N = 8. (C) Representative fluorescence microscopy images of normal control (left) and tendinopathic tendons (right) stained with DAPI (blue) and a CD90 antibody (green). Boxplots depict the quantified co-localization of DAPI and CD90 (CD90^+^ cells) calculated as percentage of the total number of cells. N = 7. (D) Representative fluorescence microscopy images of normal control (left) and tendinopathic tendons (right) stained with DAPI (blue) and an IL6R antibody (magenta). Boxplots depict the quantified co-localization of DAPI and IL6R (IL6R^+^ cells) calculated as percentage of the total number of cells. N = 7. In all boxplots, each datapoint was calculated from 8 representative fluorescence microscopy images taken from the same sample. The colored datapoint matches the presented fluorescence microscopy image. The upper and lower hinges correspond to the first and third quartile (25th and 75th percentile), the middle one to the median, the whiskers extend from the hinges no further than 1.5 times the interquartile range, and the points beyond the whiskers are treated as outliers. Results of the statistical analysis are indicated as follows: ^n.s.^p >= 0.05, *p < 0.05, **p < 0.01. The applied statistical test was the Student’s t-test.

In normal control tendons, the fluorescent signal stemming from the IL-6 antibody (**Figure 2B**, left side, red) appeared to be confined to the extrinsic compartment, which we identified based on the clustering of cells with a roundish nucleus (blue). In tendinopathic tendons (**Figure 2B**, right side), it was challenging to identify the extrinsic compartment due to the characteristic change in cell shape from elongated to more roundish in both the extrinsic compartment and the load-bearing core tissue. While the signal of the IL-6 antibody was more evenly distributed over the tendinopathic tissue section and tendon compartments compared to the control, it was still more prominent around roundish than elongated cells. Using the nuclear staining as a mask, we attributed IL-6 secretion to cells based on spatial proximity. The percentage of IL-6 secreting cells was only slightly increased in tendinopathic compared to healthy control tendons (**Table 2**).

**Table 2:** Percentages of IL-6^+^, CD90^+^, and IL6R^+^ cells of all cells in tendinopathic and normal control tissues derived from human patients. The values are given as median (IQR)

| Condition | IL-6<sup>+</sup> % all cells | CD90<sup>+</sup> % all cells | IL6R<sup>+</sup> % all cells |
| --- | --- | --- | --- |
| Tendinopathic | 33.3(38.4) | 45.6(36.3) | 37.5(41.2) |
| Normal control | 22.5(17.2) | 4.3(9.0) | 5.3(10.2) |

The cell-surface protein CD90 is a common marker of reparative fibroblasts.^[64,65]^ Here, we visually detected its signal on only a few cells in the normal control tendons (**Figure 2C**, left side, green) but on a large number of cells in the tendinopathic tendons (**Figure 2C**, right side). The subsequent spatial proximity-based quantification confirmed this initial visual impression by detecting a statistically significant difference in the percentage CD90^+^ in the normal control compared to the tendinopathic tendon (**Table 2**).

The IL-6 receptor (IL6R) is another central part of the IL-6 signaling cascade. While some cells in the normal control tendons stained positively for IL6R, many more seemed to be present in the tendinopathic tendons. We could confirm this again with spatial proximity-based quantification detecting a statistically significant difference in the percentage of IL6R^+^ cells in normal control compared to tendinopathic tendons (**Table 2**).

### Both IL-6 and IL6R appear in spatial proximity to CD90^+^ and CD68^+^ cells in non-sheathed human tendinopathic tendons

In another study conducted in mouse tendons, immune cells such as macrophages were reported as major sources of IL6R during tendon growth.^[66]^ We therefore co-stained cells with IL-6, IL6R, and the established human macrophage surface marker CD68 to see whether this was also true in the context of human tendinopathic tendons. To further check whether (reparative) fibroblasts could indeed be involved in IL-6 – IL6R signaling as initially hypothesized, we also co-stained human tendinopathic tendons with CD90, IL-6, and IL6R. We again used tendinopathic tissues extracted from diseased biceps tendons (**Figure 3A** and **Supplementary Figure 5**).

**Figure 3:**
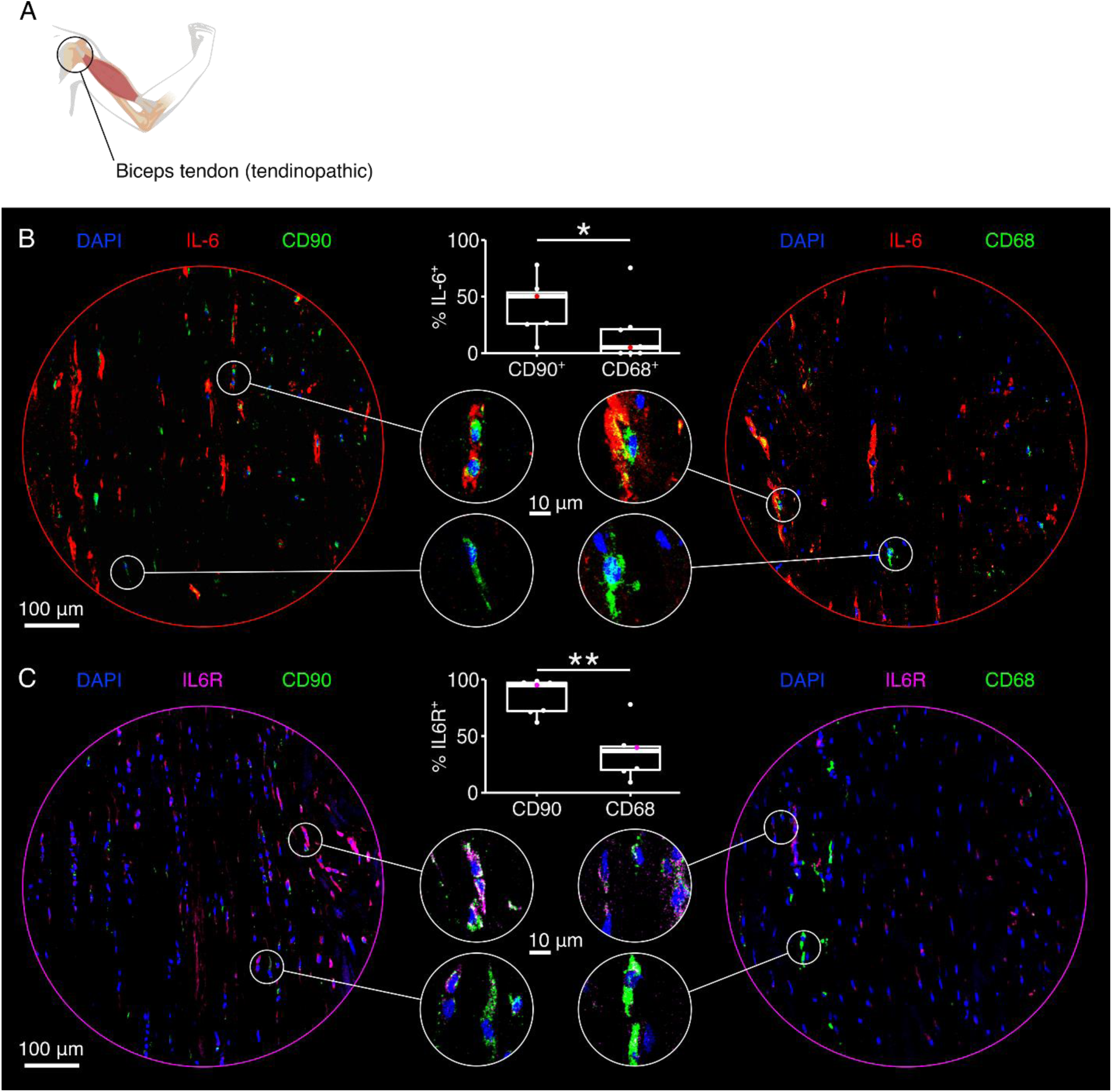
Co-localization of IL-6, IL6R, CD90, and CD68 in tendinopathic human tendons (non-sheathed) (A) Illustrative depiction of the origin of the tendon used in this experiment (painful biceps tendons, **Supplementary Figure 4**). (B) Representative fluorescence microscopy images of tendinopathic tendons stained with DAPI (blue), an IL-6 antibody (red), and either a CD90 antibody (left image, green) or a CD68 antibody (right image, green). Boxplots depict the quantified co-localization of DAPI, IL-6, and either CD90 (N = 7) or CD68 (N = 8) calculated as percentage of the number of DAPI^+^IL-6^+^ cells. (C) Representative fluorescence microscopy images of tendinopathic tendons stained with DAPI (blue), an IL6R antibody (magenta), and either a CD90 antibody (left image, green) or a CD68 antibody (right image, green). Boxplots depict the quantified co-localization of DAPI, IL-6, and either CD90 or CD68 calculated as percentage of the number of DAPI^+^IL6R^+^ cells. N = 7. In all boxplots, each datapoint was calculated from 8 representative fluorescence microscopy images taken from the same sample. The colored datapoint matches the presented fluorescence microscopy image. The upper and lower hinges correspond to the first and third quartile (25th and 75th percentile), the middle one to the median, the whiskers extend from the hinges no further than 1.5 times the interquartile range, and the points beyond the whiskers are treated as outliers. Results of the statistical analysis are indicated as follows: *p < 0.05, **p < 0.01. The applied statistical test was the Mann-Whitney-Wilcoxon Test.

We found a high percentage of cells in close spatial proximity to fluorescent signals generated by the IL-6 antibodies (**Figure 3B**, left side, red) to be CD90^+^ (green and blue), identifying them as a likely source of IL-6. Again, IL-6 secreting, CD90^+^ cells assumed a more roundish phenotype in contrast to the CD90^+^ cells not secreting IL-6. The overlap between signals from CD68^+^ cells (**Figure 3B**, right side, green and blue) and the IL-6 antibodies (red) seemed less pronounced. Indeed, quantification of the spatial signal overlay showed a significantly higher percentage of cells in spatial proximity to IL-6 to be CD90^+^ rather than CD68^+^ cells(**Table 3**).

The presence of IL6R on different cell populations could provide cues on the targets of IL-6 signaling in tendinopathic tendons. In the tendinopathic sections probed here, almost all cells that stained positively for IL6R (**Figure 3C**, left side, magenta) were also CD90^+^ (green and blue) but less than half of them were CD68^+^ cells (**Figure 3C**, right side, green and blue). Quantifying the difference based on spatial proximity confirmed this impression (**Table 3**) and the statistical analysis judged it to be statistically significant. The staining antibody and quantification method deployed here likely cannot discriminate between IL6R produced by the cell carrying it and IL6R that was solubilized participates in trans-signaling.

We conclude from the above analysis that IL-6 signaling in non-sheathed tendinopathic tendons plausibly contributes to chronic tendinopathic hallmarks like hypercellularity and aberrant matrix turnover, potentially by activating reparative (CD90^+^) fibroblast populations through IL6R. On this basis, we sought to directly test whether a causal relationship exists between the observed changes in IL-6 signaling and the associated disease processes. To this end, we harnessed an *in vitro* assembloid model of inter-compartmental crosstalk to better dissect the role of IL-6 in tendinopathy.

**Table 3:** CD90^+^ and CD68^+^ cells as percentages of IL-6^+^ and IL6R^+^ cells in tendinopathic tissues derived from human patients. The values are given as median(IQR)

|  | CD90<sup>+</sup> (median(IQR)) | CD68<sup>+</sup> (median(IQR)) |
| --- | --- | --- |
| % of IL-6<sup>+</sup> | 50.2(27.6) | 5.02(21.1) |
| % of IL6R<sup>+</sup> | 95.0(25.1) | 37.0(20.6) |

### IL-6 signaling by tendon core explants activates extrinsic fibroblasts

We have previously validated a hybrid explant // hydrogel assembloid model that reproduces the *in vivo* tissue compartment interface between the load-bearing tendon core and the extrinsic compartment (i.e. epitenon and paratenon) of non-sheathed tendons.^[54,67]^ We exploited this model to test whether IL-6 signaling across tissue compartments could activate fibroblast populations in the peritendinous space in a manner that mimics the IL-6 signaling signatures we uncovered in the human data analysis (**Figure 4A**). Briefly, we isolated and clamped mouse tail tendon fascicles to represent the tendon core while selecting (mainly Scx^+^ and CD146^+^) fibroblasts from digested Achilles tendons based on plastic adherence growth and surface marker expression as established previously and repeated here (**Figure 4B, Supplementary Figure 6**).^[54,68]^ To form the artificial extrinsic compartment, we encapsulated these fibroblast populations into a collagen hydrogel which we then let polymerize around the clamped core explants (**Figure 4C**).

**Figure 4:**
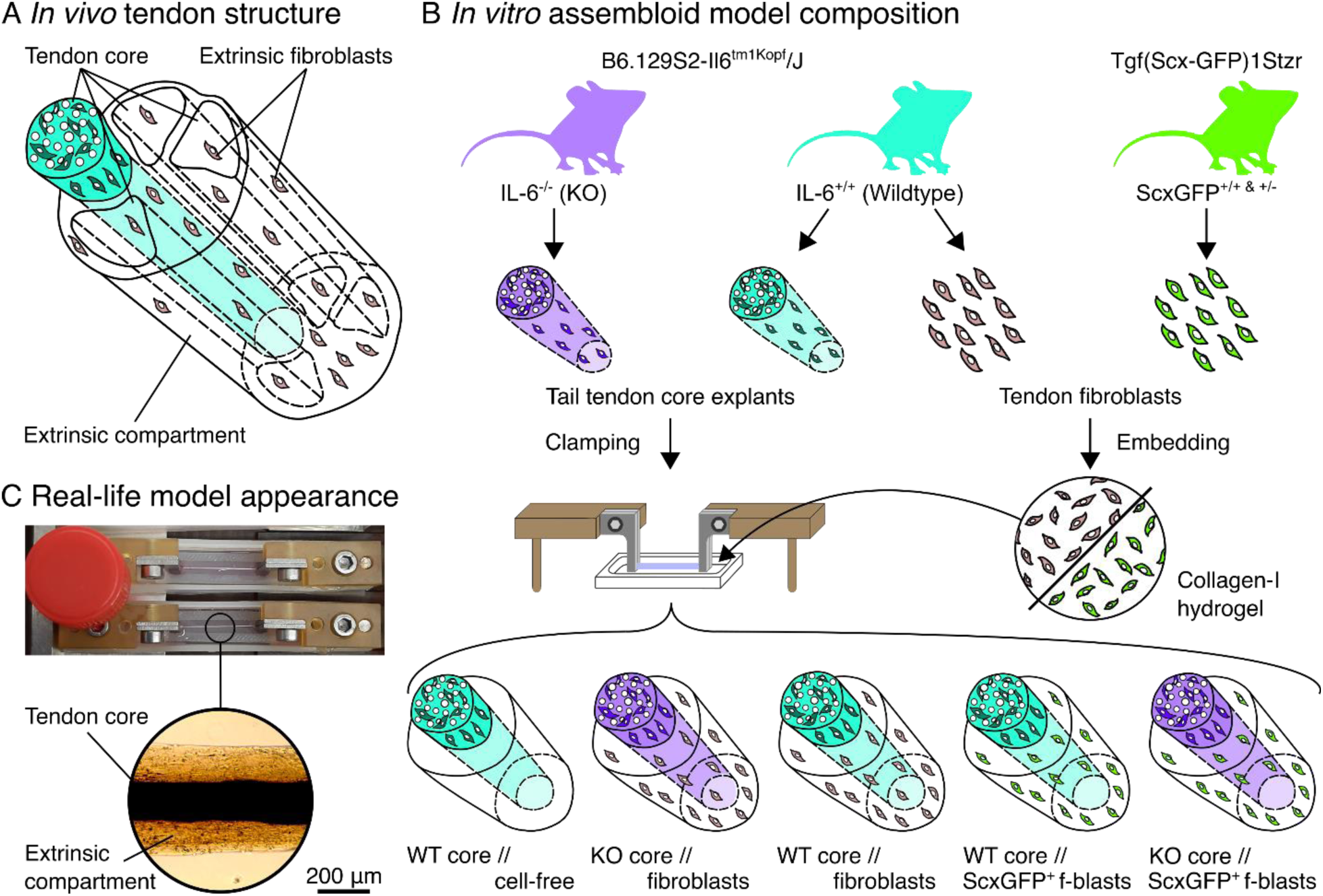
Concept behind the *in vitro* hybrid explant // hydrogel assembloid system. (A) Abstract representation of the *in vivo* load-bearing tendon core subunits (light blue / white) surrounded by the extrinsic compartment (white) containing i.e. extrinsic fibroblasts (light brown). (B) Sources of the *in vitro* model system components with the IL-6 knock-out core (KO core) in violet, the IL-6 wildtype core (WT core) in light blue, the IL-6 wildtype fibroblasts in light brown, and the ScxGFP^+^ fibroblasts in green. Core explants were clamped, and the fibroblasts embedded in a (liquid) collagen solution before crosslinking the mixture into a hydrogel around the clamped core explants in various combinations. (C) Photographic and light microscopic images of the *in vitro* assembloid model system. Lid of a 15 ml Falcon® tube (Ø: 17 mm) used for scale.

In a separate experiment, we verified the presence and located the cellular sources of IL-6 and IL6R using supernatant and flow cytometric analysis respectively (**Supplementary Figure 7**). Both were present in the supernatant (IL-6) and on CD45^+^ cell populations (IL6R) from core explants, but not in the supernatant or on the surface of extrinsic fibroblasts cultured in a collagen hydrogel. This indicates that the followingly described effects of IL-6 on extrinsic fibroblasts could be dominated by trans-signaling.

To dissect the effect of IL-6 signaling on the extrinsic target populations, we integrated either wildtype-derived (WT) or IL-6 knock-out-derived (KO) explants from the B6.129S2-Il6^tm1Kopf^/J^[69]^ mouse line (**Figure 5A**). We then performed bulk RNA-sequencing on the extrinsic populations after one week of co-culture in tendinopathic niche conditions.^[54,70,71]^

**Figure 5:**
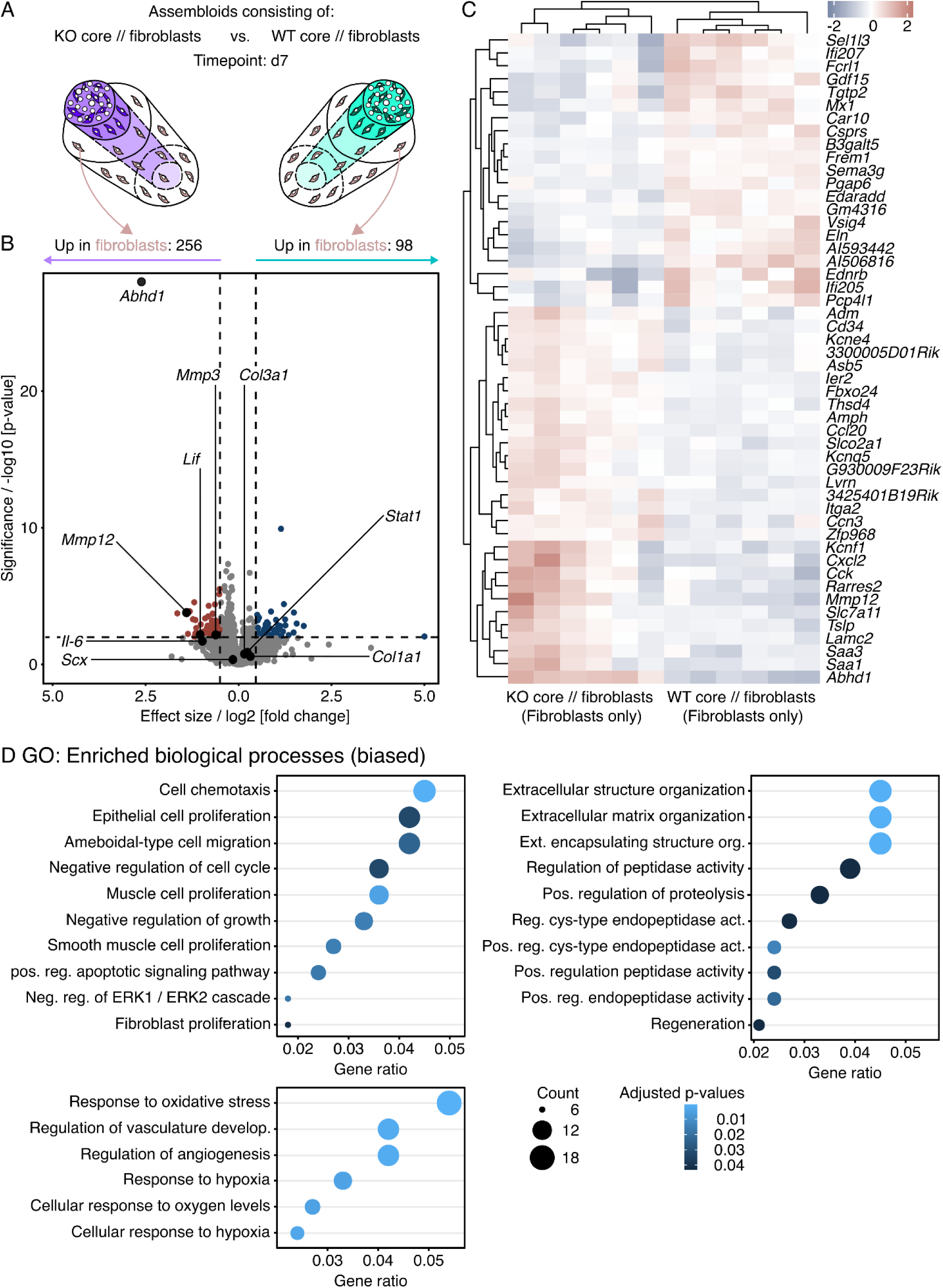
Transcript changes in hydrogel-embedded fibroblasts seeded around an IL-6 knock-out (KO) core explant compared those seeded around a wildtype (WT) core. (A) Illustration of the assembloid combinations compared here (KO core // fibroblasts vs. WT core // fibroblasts), the assessed timepoint (d7), and the analyzed compartment (extrinsic fibroblasts only). (B) RNA-seq volcano plot of differentially expressed genes (DEGs). Genes colored in red have a log2 (fold change) > 0.5, a p-value < 0.05, and are considered to be significantly increased in the extrinsic compartment of KO core // fibroblast assembloids. Genes colored in blue have a log2 (fold change) < - 0.5, a p-value < 0.05, and are considered to be significantly increased in the extrinsic compartment of WT core // fibroblast assembloids. The log2 and p-value thresholds are represented by the dashed lines. (C) Unsupervised hierarchical clustering of the top 50 differentially expressed genes. Genes are clustered by color with positive (red) or negative (blue) row-scaled z-scores. Columns represent individual samples. N=6. (D) Dotplots depicting a selection of GO annotations significantly enriched (adjusted p-value < 0.05) by the DEGs. The selection was biased by GO biological process annotations enriched in the human dataset (**Figure 1E**). The color of the circles represents their adjusted p-value, the size the number of enriched genes (count), and the x-axis the number of enriched genes in ratio to the total number of genes annotated to the gene set (gene ratio).

Overall, integration of an IL-6 KO core increased transcripts of 256 genes in the surrounding extrinsic compartment, decreased transcripts of 98 genes, and left 15,295 unchanged (**Figure 5B** & **C)**. After mapping the significant transcript changes to biological processes in the GO database, we conducted a biased search for processes matching those dysregulated in human tendinopathic tendons. Out of the 195 significantly enriched biological processes (adjusted p-value < 0.05) in the extrinsic compartment of KO core // fibroblast assembloids, 10 (5.1%) could be linked to changes in local cellularity and another 10 to ECM turnover (**Figure 5D**).

When stratifying the respective contribution of transcripts increased and decreased around a KO core compared to a WT core to the top 20 enriched GO biological processes and significantly enriched MSigDB mouse hallmarks, we found that IL-6 signaling by the core correlated positively with processes aimed at increasing cellularity and ECM turnover (**Supplementary Figure 8B-D**), which are both hallmarks of tendinopathic tissues and indicators of an activated tissue state.

To verify the transcript-level differences connected to hypercellularity also on the tissue-level, we next performed a proliferation and migration analysis in our assembloid model system.

### IL-6 signaling by tendon core explants stimulates cell proliferation and Scx^+^ cell recruitment to the signaling tendon core

Using the assembloid model, we investigated whether IL-6 signaling could play a causative role in the hypercellularity that is a major hallmark of tendinopathy. Closely mimicking human tendinopathic tendons, we indeed found cellularity-increasing biological processes to be positively enriched in cell populations around WT compared to those around IL-6 KO tendon core explants. We then assessed whether these IL-6-dependent transcript-level changes would translate to an increased cell density. To do this, we seeded tendon fibroblast populations isolated from ScxGFP^+^ mice (co-expressing the tendon marker Scleraxis (Scx) with a green fluorescent protein) into the hydrogel extrinsic compartment of our assembloids and incorporated either a WT or an IL-6 KO core into the center (**Figure 6A, top panels)**.

**Figure 6:**
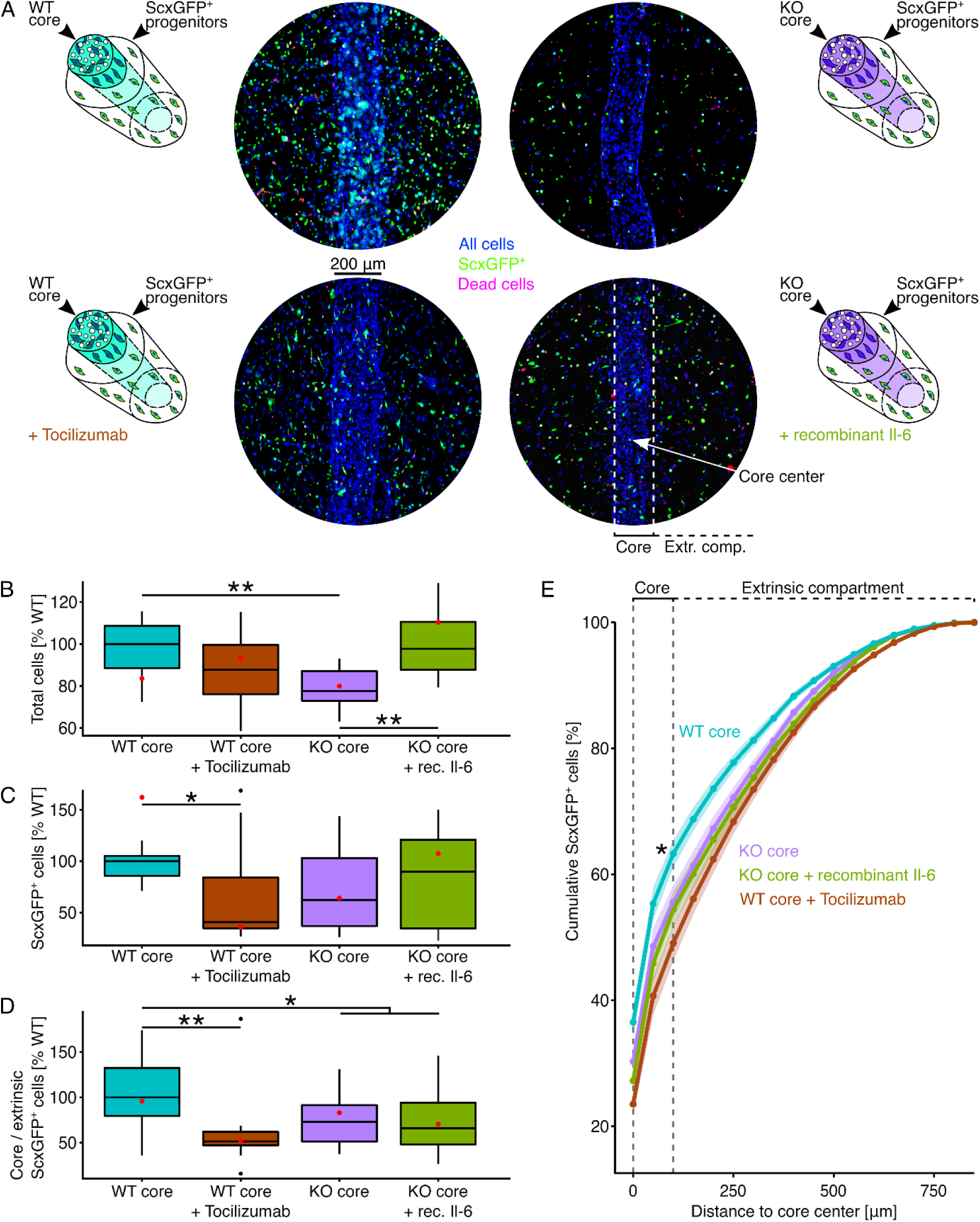
Cell proliferation around and ScxGFP^+^ fibroblast recruitment to core explants. (A) Illustrative depictions and representative fluorescence microscopy images of wildtype (WT) core explants surrounded by fibroblast populations from ScxGFP^+^ mice cultured with or without Tocilizumab (10 µg/ml), and IL-6 knock-out (KO) explants cultured with or without recombinant IL-6 (25 ng/ml) for 7 days. All cells are colored in blue (NucBlue), ScxGFP^+^ fibroblasts in green (GFP), and dead cells in red (EthD). (B, C, D) Boxplots depicting the total number of cells, the number of ScxGFP^+^ cells, and the ratio between core-resident and extrinsic ScxGFP^+^ cells normalized to the WT median. Each datapoint was calculated from 3 representative fluorescence microscopy images taken from the same sample. The red datapoint matches the presented fluorescence microscopy image. The upper and lower hinges correspond to the first and third quartile (25th and 75th percentile), the middle one to the median, the whiskers extend from the hinges no further than 1.5 times the interquartile range, and the points beyond the whiskers are treated as outliers. (E) Lineplot depicting the cumulative percentage of ScxGFP^+^ cells depending on their distance from the center line of the core explant. The points and the line represent the mean cumulative percentages and the error bands the standard error of the mean (sem). The dashed line indicates locations inside the core area. N=12. Results of the statistical analysis are indicated as follows: *p < 0.05, **p < 0.01. The applied statistical test was the Mann-Whitney-Wilcoxon Test.

Representative fluorescence microscopy images taken after seven days in co-culture confirmed a higher total cell number in WT core // ScxGFP^+^ fibroblast assembloids compared to KO core // ScxGFP^+^ fibroblast assembloids. These cell number differences were not confined to ScxGFP^+^ fibroblasts (green) but extended to other populations (blue). In addition, ScxGFP^+^ fibroblasts only accumulated around the WT core in WT core // ScxGFP^+^ fibroblast assembloids, presumably either through increased core-directional migration or faster proliferation closer to the core. These observations are consistent with IL-6 being essential to increased cellularity and core (damage)-directed migration in this model.^[54]^

We went on to confirm these visual impressions using quantitative methods, finding a significantly increased total cell number in assembloids with a WT core compared to those with a KO core (**Table 4 & Figure 6B**). The effect of IL-6 signaling on the proliferation of ScxGFP^+^ fibroblasts (**Table 4 & Figure 6C**) was less pronounced compared to that on all populations, but the trend remained the same. To quantify migration, we analyzed the spatial distribution of ScxGFP^+^ fibroblasts by calculating the ratio between core-resident and extrinsic ScxGFP^+^ fibroblasts (**Table 4 & Figure 6D**). The WT core // ScxGFP^+^ fibroblast assembloids exhibited the highest core-resident to extrinsic ScxGFP^+^ fibroblast ratio and KO core // ScxGFP^+^ fibroblast assembloids had a significantly lower core-resident to extrinsic ScxGFP^+^ fibroblast ratio. The cumulative spatial distribution of ScxGFP^+^ fibroblasts (**Table 4 & Figure 6E**) supported these insights.

**Table 4:** Total cell numbers, ScxGFP^+^ cell numbers, and the ratios between core-resident and extrinsic ScxGFP^+^ cells in assembloids. The values were normalized to the WT median and are given as median(IQR)

| **Condition** | **Total cell number**<br>**norm. to WT**<br>**(median(IQR))** | **ScxGFP<sup>+</sup> cell number**<br>**norm. to WT**<br>**(median(IQR))** | **Core / extrinsic**<br>**ScxGFP<sup>+</sup> ratio,**<br>**norm. to WT**<br>**(median(IQR))** |
| --- | --- | --- | --- |
| WT core // ScxGFP<sup>+</sup><br>fibroblasts | 100(20.1)% | 100(19.4)% | 100(52.3)% |
| WT core // ScxGFP<sup>+</sup><br>fibroblasts +<br>Tocilizumab | 87.7(23.5)% | 40.7(49.3)% | 51.5(14.8)% |
| KO core // ScxGFP<sup>+</sup><br>fibroblasts | 77.6(14.1)% | 62.2(66.1)% | 73(40)% |
| KO core // ScxGFP<sup>+</sup><br>fibroblasts + IL-6 | 97.7(22.8)% | 89.7(86.1)% | 66.1(45.9)% |

To further confirm the specific impact of IL-6 signaling on overall cell proliferation and core-directed migration of Scx^+^ fibroblasts, we desensitized the WT core // fibroblast assembloids to IL-6 by neutralizing IL6R with Tocilizumab and attempted to rescue the KO core // fibroblast assembloids by adding recombinant IL-6 to compensate for their reduced IL-6 levels (**Figure 6A, bottom panels**). In alignment with the previous results and the hypothesis, IL-6 desensitization decreased the total cell number in trend (**Table 4 & Figure 6B**), the ScxGFP^+^ cell number significantly (**Table 4 & Figure 6C**,), and the ratio between core-resident and extrinsic ScxGFP^+^ fibroblasts significantly as well (**Table 4 & Figure 6D**). The addition of recombinant IL-6 to KO core // fibroblast assembloids significantly increased the total cell number and the number of ScxGFP^+^ cells, rescuing the wildtype phenotype of IL-6 enhanced cell proliferation in the extrinsic compartment. However, core-directed migration was not rescued by recombinant IL-6.

Fully in line with transcript signature changes detected in the extrinsic compartment, these data suggest that IL-6 signaling increased local cellularity in at least one of two ways. First, IL-6 stimulated both overall and specific ScxGFP^+^ cell proliferation. Second, IL-6 gradient effects (i.e. IL-6 induced secondary gradients) caused core-directed ScxGFP^+^ cell migration.

### Disrupting IL-6 signaling does not detectably alter Scx^+^ cell proliferation or recruitment into an acutely damaged Achilles tendon in vivo

After clarifying the role of IL-6 in activating fibroblasts in the assembloid model of chronic tendon disease, we sought to assess whether IL-6 signaling also enhances overall cell proliferation and migration of Scx^+^ fibroblasts to acute tendon damage. To test this, we first bred IL-6 wildtype and IL-6 KO mice with ScxGFP^+^ mice. Then, we assessed the presence of ScxGFP^+^ cells in the Achilles tendons (AT) of four IL-6 WT mice compared to those of four IL-6 KO mice 14 days after Achilles tenotomy. In addition, we used an EdU staining to assess the proliferation of cells within the healing tendon (**Figure 7A**).

**Figure 7:**
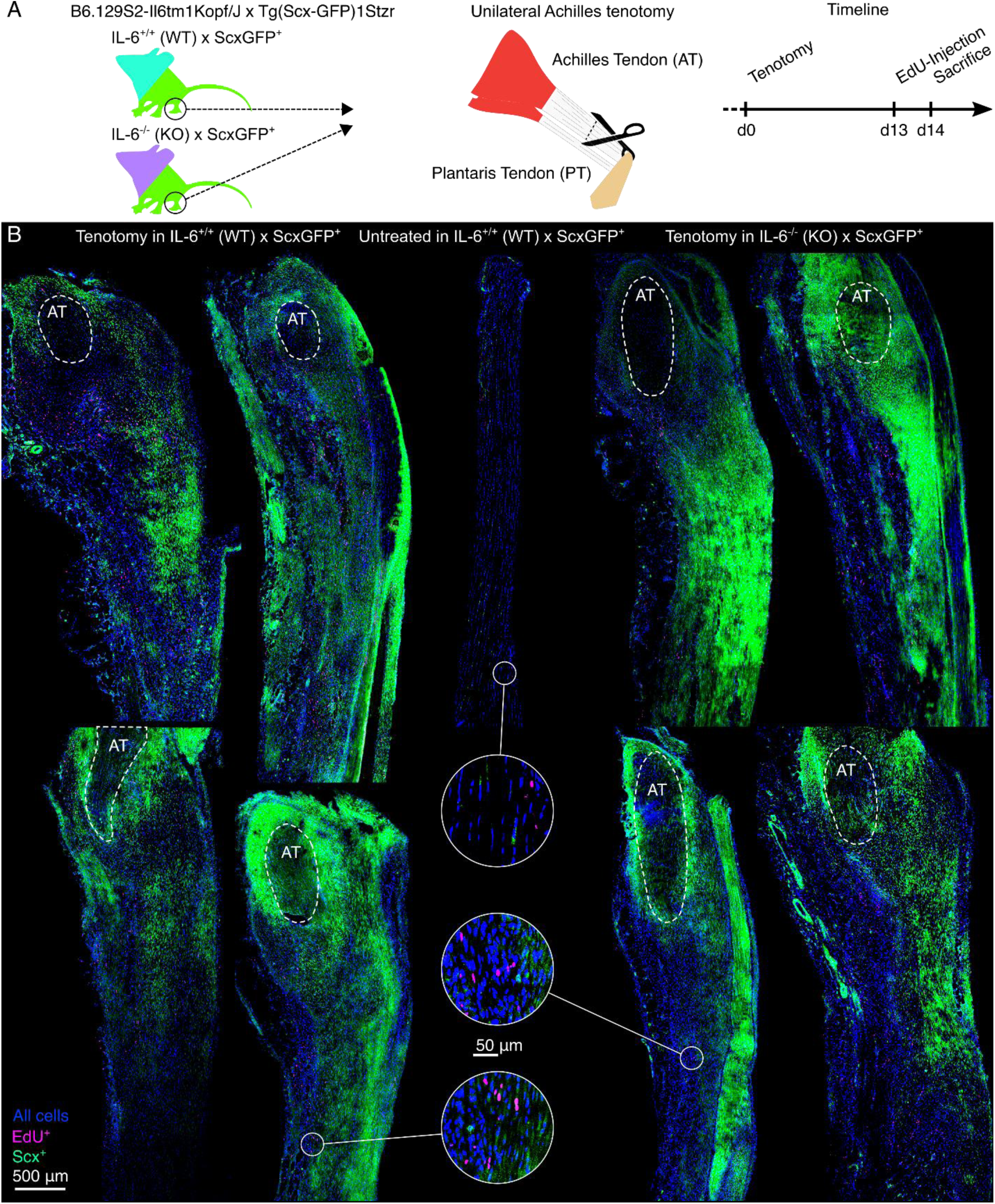
Cell proliferation around and Scx^+^ fibroblasts recruitment to acutely damaged mouse Achilles tendons 14 days after injury. (A) Illustrative depiction of the experimental setup and the time schedule. (B) Representative fluorescence microscopy images from all four mice assessed showing longitudinal mouse hindleg sections from IL-6 wildtype, ScxGFP^+^ (IL-6^+/+^ (WT) x ScxGFP^+^) Achilles tendons that underwent tenotomy (left), the contralateral untreated control (middle), as well as sections from IL-6 knock-out (IL-6^-/-^ (KO) x ScxGFP^+^) Achilles tendons that underwent tenotomy (right). In addition to the signal provided by the ScxGFP^+^ cells (green), NucBlue was used to identify all cell nuclei (blue), and EdU was used to identify proliferating cells (magenta). The dashed circles indicate the remaining Achilles tendon (AT) stump close to the calcaneus. The healing neo-tendon tissue surrounding the calcaneal AT stump bridges the gap to the AT stump connected to the calf muscles further down (not shown). In both compartments, ScxGFP^+^ cell distribution was highly variable across acutely damaged samples, with no observable trends or statistically detectable differences between the conditions.

Overall, the fluorescence microscopy images revealed a strong presence of ScxGFP^+^ cells (green) and EdU^+^ cells (magenta) in the neo-tendon (**Figure 7B**, tissue around the dashed circles) formed around the calcaneal Achilles tendon stumps (**Figure 7B**, AT within the dashed circles) after tenotomy (**Figure 7B**, left), but not in undamaged hindleg tendons (**Figure 7B**, middle). Similar levels of overall cellularity and presence of ScxGFP^+^ and EdU^+^ cells were observed in the calcaneal Achilles tendon stump and the surrounding neo-tendon of IL-6 WT and IL-6 KO mice. No observable trends or statistically detectable differences in the highly variable and complex ScxGFP^+^ cell migration patterns were detected between IL-6 WT and IL-6 KO mice (**Figure 7B**, left & right).

### Activated, recruited, and proliferating extrinsic fibroblasts promote tendinopathy hallmarks in tendon core explants

Building upon the evidence that IL-6 potentiates tendon fibroblast activation and migration to damage *in vitro*, we then sought to clarify the nature of interactions between these recruited repair cells and the damaged tissue. We asked whether these activated fibroblasts might be capable of driving disease-relevant tissue processes. To assess this, we first looked at transcriptional changes induced in core explants when fibroblasts were present in the artificial extrinsic compartment by comparing them to explants cultured in an initially cell-free hydrogel (**Figure 8A**).

**Figure 8:**
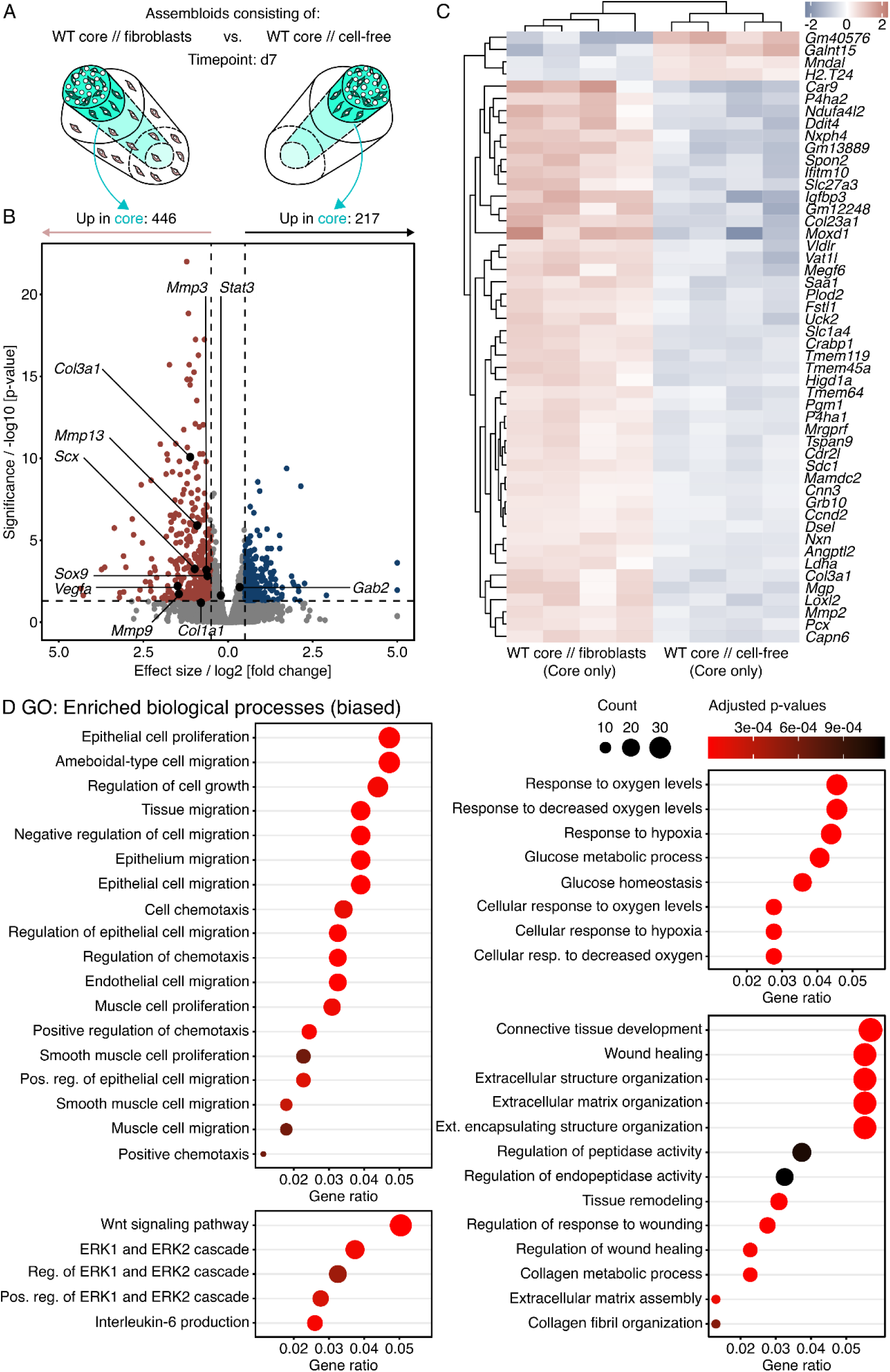
Transcript analysis of differentially regulated genes and pathways in wildtype (WT) core explants surrounded by a hydrogel seeded with fibroblasts compared to a WT core surrounded by a cell-free hydrogel. (A) Illustration depicting the assembloid combinations compared here (WT core // fibroblasts vs. WT core // cell-free), the assessed timepoint (d7), and the analyzed compartment (core only). (B) RNA-seq volcano plot of differentially expressed genes (DEGs). Genes colored in red have a log2 (fold change) > 0.5, an adjusted p-value < 0.05, and are considered to be significantly increased in the core of WT core // fibroblast assembloids. Genes colored in blue have a log2 (fold change) < - 0.5, an adjusted p-value < 0.05, and are considered to be significantly increased in in the core of WT core // cell-free assembloids. The log2 and p-value thresholds are represented by the dashed lines. (C) Unsupervised hierarchical clustering of the top 50 differentially expressed genes. Genes are clustered by color with positive (red) or negative (blue) row-scaled z-scores. Columns represent individual samples. N=4. (D) Dotplots depicting a selection of GO annotations significantly enriched (adjusted p-value < 0.05) by the DEGs. The selection was biased by GO biological processes and GSEA hallmark annotations enriched in the human dataset (**Figure 1C & E**). The color of the circles represents their adjusted p-value, the size the number of enriched genes (count), and the x-axis the number of enriched genes in ratio to the total number of genes annotated to the gene set (gene ratio).

Exposing WT core explants to fibroblasts (WT core // fibroblasts) for seven days increased 446 transcripts, decreased 217 transcripts, and left 19,694 transcripts unchanged (**Figure 8B** & **C**). In line with the previous paragraphs reporting fibroblast migration *in vitro*, some of the increased transcripts (i.e. Scx and Sox9) indicated an enrichment of Scx^+^ and / or Sox9^+^ fibroblasts in the WT core explants of WT core // fibroblast assembloids compared to those of WT core // cell-free assembloids. Similarly, GSEA on the full MSigDB cell type signature gene sets proposed an amplified contribution of fibroblasts, fibroblast-like, and progenitor cells to the emerging assembloid phenotype (**Supplementary Figure 8B**). *In vivo*, extrinsic (i.e. paratenon-derived) fibroblasts differentially express selected genes compared to tendon core (i.e. tendon proper-derived) fibroblasts.^[72]^ The GO gene sets annotated with these differentially expressed genes (DEGs) overlap with those enriched by DEGs between WT core // fibroblast and WT core // cell-free assembloids (**Supplementary Figure 9A-D**). This could mean that the core explants exposed to extrinsic fibroblasts change into more paratenon-like tissue and again highlights the contribution of extrinsic fibroblast migration and accumulation to assembloid behavior.

To compare this phenotype to human tendinopathic tendons, we looked for ECM turnover-related transcripts that were enriched in human tendinopathic tendons compared to normal controls. Indeed, transcripts for *Col3a1*, *Col1a1*, *Mmp13*, *Mmp3*, and *Mmp9* were increased in core explants co-cultured with fibroblasts (**Figure 8A** & **Table 5**). When combined through ORA, many of the top 30 biological processes enriched by significantly changed transcripts (adjusted p-value < 0.01) were also enriched in human tendinopathic tendons (**Supplementary Figure 11**). The curated list presented here (**Figure 8D**) pinpoints significantly enriched processes likely to be involved in tendinopathic hallmarks such as ECM turnover & tissue development, hypoxia & glucose metabolism, and hypercellularity.

**Table 5:** Effect sizes and p-values for selected transcripts. The data describes differences in transcripts between the core explants from WT core // fibroblasts and those from WT core // cell-free assembloids.

| Transcript | Effect size | P-value |
| --- | --- | --- |
| *Col3a1* | 1.1 | 8.4e-11 |
| *Col1a1* | 0.8 | 0.06 |
| *Mmp13* | 0.91 | 1.2e-06 |
| *Mmp3* | 0.64 | 0.0006 |
| *Mmp9* | 1.45 | 0.019 |
| *Scx* | 0.98 | 0.00006 |
| *Sox9* | 0.6 | 0.001 |

Overall, it appears that extrinsic fibroblasts are sufficient to invoke several tendinopathic hallmarks in tendon core explants and accelerated catabolic matrix turnover in particular. We have previously reported an increase of IL-6 in the supernatant of WT core // fibroblast assembloids that correlated with an increased catabolic breakdown of the core.^[54]^ The insights gained here connect this catabolic breakdown to gene sets involved in ECM remodeling. Another set of previously published experiments suggests that the ERK1/2 signaling cascade enriched in the core of WT core // fibroblast favors tissue breakdown as well.^[70,71]^

### Disrupting IL-6 signaling in core explants diminishes emergence of tendinopathic hallmarks

So far, our results have shown that IL-6 signaling enhances proliferation and migration of fibroblasts towards the tendon core and that the presence of fibroblasts invokes tendinopathy-like changes in the tendon core *in vitro*. Consequently, the last step was to see whether an IL-6 knock-out not only prevents the fibroblast migration and proliferation, but also reduces fibroblast-invoked tendinopathic hallmarks in the core. To assess this, we again studied assembloids containing an IL-6 KO core, but this time focused on biological processes emerging in the core by leveraging bulk RNA-sequencing (**Figure 9A**).

**Figure 9:**
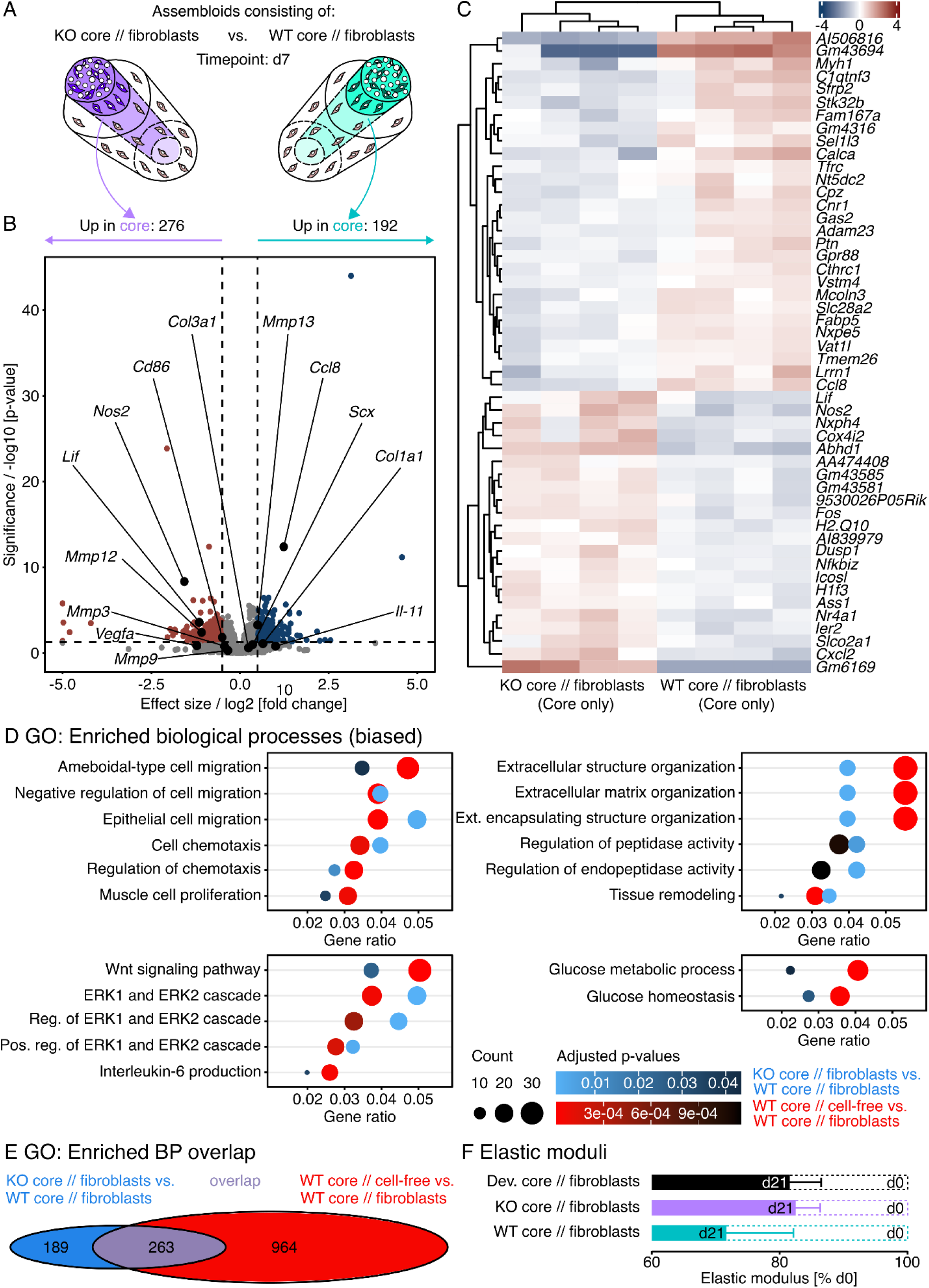
Transcript analysis of differentially regulated genes and pathways in IL-6 knock-out (KO) core explants surrounded by a hydrogel seeded with fibroblasts compared to a wildtype (WT) core surrounded by fibroblasts. (A) Illustration depicting the assembloid combinations compared here (KO core // fibroblasts vs. WT core // fibroblasts), the assessed timepoint (d7), and the analyzed compartment (core only). (B) RNA-seq volcano plot of differentially expressed genes (DEGs). Genes colored in red have a log2 (fold change) > 0.5, a p-value < 0.05, and are considered to be significantly increased in the core of KO core // fibroblast assembloids. Genes colored in blue have a log2 (fold change) < - 0.5, a p-value < 0.05, and are considered to be significantly increased in the core of WT core // fibroblast assembloids. The log2 and p-value thresholds are represented by the dashed lines. (C) Unsupervised hierarchical clustering of the top 50 differentially expressed genes. Genes are clustered by color with positive (red) or negative (blue) row-scaled z-scores. Columns represent individual samples. N=4. (D) Dotplots depicting a selection of GO annotations significantly enriched (adjusted p-value < 0.05) by the DEGs in both the WT core // cell-free vs. WT core // fibroblast assembloid comparison (red to black gradient) and the KO core // fibroblast vs. WT core // fibroblast assembloid comparison (light blue to black gradient). The selection was biased by enriched GO biological process and GSEA hallmark annotations in the human dataset (**Figure 1C & E**). The color gradient of the circles represents their adjusted p-value, the size the number of enriched genes (count), and the x-axis the number of enriched genes in ratio to the total number of genes annotated to the gene set (gene ratio). (E) Venn-Diagramm depicting the number and the overlap (violet) of significantly enriched GO annotations for biological processes between the WT core // cell-free vs. WT core // fibroblast assembloid comparison (red) and the KO core // fibroblast vs. WT core // fibroblast assembloid comparison (blue). (F) Linear elastic moduli of devitalized (Dev.), IL-6 knock-out (KO), and wildtype (WT) core explants surrounded by hydrogel-embedded fibroblast populations at day 21 normalized to day 0. N=8. The data are displayed as barplots with mean ± standard error of the mean (sem). The applied statistical test was the Mann-Whitney-Wilcoxon Test and yielded no significant differences.

On the transcript level, we found 276 upregulated, 192 downregulated, and 20,204 unchanged genes in the core of KO core // fibroblast assembloids compared to that of WT core // fibroblast assembloids (**Figure 9B & C**). To see whether an IL-6 knock-out would partially reverse fibroblast-invoked hallmarks, we matched the list of differentially expressed genes (p-value < 0.01) to the signatures in the GO database and then compared the surfacing enriched biological processes with those enriched by differentially expressed genes between the core of a WT core // fibroblast assembloid and a WT core // cell-free assembloid. The largest overlap lay in the signaling pathways (Wnt, ERK1/2, and IL-6), where 5/5 signatures for biological processes remained similarly enriched (**Figure 9E**). We found slightly fewer overlapping signatures connected to ECM turnover (6/14) and cellularity (6 /18) while there seemed to be a disconnect in hypoxia & glucose metabolism (2/8). Overall, about a third of all signatures enriched by the presence of fibroblasts were also enriched by the IL-6 KO (**Figure 9E**). In contrast to the signatures emerging from the presence of fibroblasts however, the respective contribution of transcripts increased and decreased by the IL-6 KO to the enriched GO biological processes and molecular functions indicates a decreasing cellularity and ECM turnover in the core (**Supplementary Figure 12**), which could mean that IL-6 signaling contributes to the gene expression behind these tendinopathy hallmarks.

As we have already demonstrated the tissue-level effects of IL-6 signaling on cellularity (**Figure 6**), we next wanted to examine the tissue-level consequences of changed transcript signatures for ECM turnover on core biomechanics. To do so, we measured the assembloid’s linear elastic modulus as an indicator for their ability to resist longitudinal tension, which is one of the main functions of adult tendons (**Figure 9F**). In WT core // fibroblast assembloids, the linear elastic modulus decreased the most between the initial clamping of the core explant and 21 days of co-culture (light blue). The linear elastic modulus of assembloids containing a KO core (violet) or a WT core explant devitalized through multiple freeze-thaw cycles (black) decreased as well, but not as fast or as strongly. In connection with the bulk RNA-seq data, the changes in the assembloid’s ability to resist tension suggest that IL-6 signaling accelerates (catabolic) ECM turnover.

Normally, it would be hard to predict the effect of an accelerated catabolic ECM turnover *in vitro* on wound healing *in vivo* since both ECM degradation and synthesis are required to replace damaged tissue structures. However, previous studies have already reported a delayed wound healing response in IL-6 KO mice and our experiments here suggest that the decelerated catabolic ECM turnover could be responsible for this.^[57]^

## DISCUSSION

Interleukin-6 (IL-6) is an attractive translational research target. Signaling cascades related to IL-6 are upregulated in tendon tissues after exercise, acute tissue damage, and in chronic tendon disease.^[58,60,73]^ Little is currently understood of the precise role that IL-6 plays in these processes.^[74]^ The goal of the present work was to clarify the role of IL-6 signaling in the tissues of non-sheathed tendons in the context of chronic damage, with a particular focus on inter-compartmental crosstalk between the damaged tendon core and extrinsic (reparative) fibroblasts populations targeted by it.

We first reanalyzed existing microarray data of (non-sheathed) tendinopathic tendons to verify a hypothesized (dys)functional role of IL-6 that seems to run partially via the JAK/STAT pathway enriched by transcripts increased in the tendinopathic samples. The JAK/STAT pathway interweaves with ERK1/2 downstream, which fits with recent data from our lab showing that ERK inhibition alone can prevent tendon matrix deterioration while reducing the secretion of IL-11, another member of the IL-6 cytokine family that was also upregulated in the tendinopathic tendons analyzed here.^[70,71]^ More generally, overactivation of JAK/STAT/ERK has been associated with auto-immune arthritis.^[34,35]^

While analysis of human samples indicated the expression of *IL-6*, *IL6ST*, and *ADAM10* (known to transform membrane-bound IL6R into soluble IL6R) to be upregulated in tendinopathic tendons, the expression of *IL6R* itself was downregulated. We speculate that this points towards increased trans-signaling aimed at stromal cells unable to express *IL6R*, in this tendinopathic context particularly the stromal fibroblasts and fibroblast-like cell populations indicated to be enriched by the cell type signatures.^[52,75]^ This theory is supported by GO terms enriched in tendinopathic samples compared to controls relating to increased morphogenesis and wound healing, both processes supposedly powered by (reparative) fibroblasts.^[76,77]^ Most of the remaining top 20 enriched GO terms pointed towards increased cell proliferation, potentially increasing the leverage of the proliferating fibroblasts. Indeed, staining human samples with CD90, an established marker of reparative fibroblasts typically present in healing tissues,^[64,65]^ confirmed a higher percentage of CD90^+^ cells in (non-sheathed) tendinopathic compared to normal control tendons.

While beneficial in controlled dosages during normal wound healing, both excessive hypercellularity and imbalanced wound healing are hallmarks of tendinopathy.^[13,14,16,17]^ To decipher their connections to IL-6 signaling, we investigated IL-6 / IL6R concentration gradients, sources, and targets in patient-derived tissue sections. In tendinopathic patients, the presence of IL-6 over the whole tissue was smaller than expected. However, while IL-6 seemed to be largely confined to the extrinsic compartment in normal control tendons, it was distributed across compartments in tendinopathic tendons. The resulting gradient differences could play a role in the emergence of disease hallmarks.^[78]^ In contrast to the relative transcript-level decrease of *IL6R* in tendinopathic tendons indicated by the microarray, its presence was increased on the protein level. Although we can only speculate on the clearance rate of both soluble and membrane bound IL6R as well as of the cells carrying them, the mismatch in its presence on the gene transcription and the protein expression levels could be a legacy from earlier disease stages.

To identify sources and targets of IL-6 and IL6R, we co-stained the tendinopathic tissue sections with established markers for macrophages (CD68) and reparative fibroblasts (CD90). Presuming that IL-6 is more concentrated around its sources than its targets due to diffusion, it seems that although both CD68^+^ and CD90^+^ cells express IL-6, the contribution of CD90^+^ cells is significantly larger. This observation is in line with sources of IL-6 identified in growing mouse tendons.^[66]^ The number of CD90^+^ cells among IL6R^+^ cells was also significantly larger than that of CD68^+^ cells. Since it has yet to be shown that stromal fibroblasts express and translate IL6R, a better explanation might be that the IL6R on CD90^+^ cells were originally produced by i.e. the CD68^+^ cells and ended up on CD90^+^ cells as part of a trans-signaling process. Why and how CD90^+^ cells are targeted by soluble IL6R as part of IL-6 trans-signaling could be an interesting part of future research.

To dissect the specific effect of IL-6 signaling on reparative fibroblasts, we turned to tendon assembloids: hybrid explant // hydrogel models of core tendon damage and repair that were recently developed in our lab and which identified an underloaded core explant as a potentially biologically relevant source of IL-6.^[54]^ The simultaneously increased and imbalanced matrix tissue turnover perpetuated by the crosstalk between the core and an extrinsic fibroblast population seemed to put the system into a prime position to decipher the connections between IL-6, hypercellularity, and (catabolic) matrix turnover. Furthermore, their compartmentalized design adequately mimics the structure of non-sheathed tendons.

Replacing the wildtype core explant with an IL-6 KO core explant in our assembloids was sufficient to reduce the expression of genes in gene sets related to matrix turnover, proteolysis, cell proliferation, and cell migration in the extrinsic fibroblast population. We confirmed the IL-6-induced gene-level differences regarding cell migration by exploiting trackable ScxGFP^+^ fibroblasts and demonstrated effective manipulation of both recruitment and proliferation of ScxGFP^+^ cells through IL-6. We did so by using the IL-6 inhibitor Tocilizumab to desensitize resident cell populations to IL-6 and recombinant IL-6 to replace the IL-6 not secreted by an IL-6 KO core. Recombinant IL-6 rescuing proliferation but not migration of ScxGFP^+^ cells highlights the necessity of an IL-6 gradient and / or the establishment of a secondary cytokine gradient (e.g. TGF-β) by IL-6.^[19]^ Alternatively, IL-6 has also been described as an energy allocator in other musculoskeletal tissues and could in this function be accelerating a diverse range of processes (i.e. migration) by increasing the baseline cell metabolism.^[29]^ Fittingly, the extrinsic cell populations around an IL-6 KO core upregulated biological processes related to oxidative stress and hypoxia, which indicates a disrupted energy allocation. Since we found similar biological processes to be positively enriched in human tendinopathic tendons compared to controls, future studies should more closely examine the role of hypoxic signaling in the pathogenesis of tendinopathy.

To transfer these insights from our *in vitro* experiments to an *in vivo* setup, we used an Achilles tenotomy model of the *in vivo* tendon damage response. Although the number of Scx^+^ fibroblasts did increase in injured tendons, the *in vivo* experiments did not detect an effect of IL-6 on activating Scx^+^ fibroblasts^[79]^ and making them migrate into or proliferate faster in the damaged and unloaded Achilles tendon stump. In addition, previous studies with mice subjected to a patellar punch procedure reported only marginally reduced mechanical properties in the healing patellar tendon of IL-6 KO mice compared to the WT after an acute injury.^[57]^ However, we are cautious in interpreting these *in vivo* results, mainly because 14 days after an Achilles tenotomy, the tendon niche condition represents an acute rather than a chronic tendon lesion.^[49,80]^ Also, even with breeding a novel IL-6 KO x ScxGFP^+^ mouse line to alleviate widely known problems of Scx antibodies, the collagen matrix still caused considerable background noise in our images.^[57]^ Furthermore, the ablation of IL-6 might have led to the elevation of compensatory IL-6 superfamily ligands (or other attractant chemokines or cytokines) and one can currently only speculate on their *in vivo* distribution as well as the resulting patterns of fibroblast migration.^[18,73]^ To confirm this hypothesis, future *in vivo* studies could deploy IL-6 inhibitors targeting other members of the IL-6 superfamily as well.

Since the migrating Scx^+^ fibroblasts *in vitro* were apparently targeting the damaged core tissue (likely to support the limited intrinsic regenerative potential of the explanted tendon core secreting IL-6 in the first place)^[59]^, the next set of experiments we conducted in this work focused on the effects of the activated and recruited fibroblasts on the tendon core.

According to existing literature and results presented here, Scx^+^ fibroblasts are increasingly present in *in vivo* murine adult tendon lesions^[18,19]^ and depleting them alternately improves or impairs adult tendon healing depending on the timepoint of depletion.^[79,81,82]^ One underlying reason could be that adult, Scx^+^ fibroblasts hold a bi-fated potential that enables a cartilage-like differentiation when exposed to mechanical compression or tensile unloading.^[12,83]^ Our *in vitro* assembloid model captured these behaviors as well with increased Scx / Sox9 transcripts and enriched gene sets indicating a stronger presence of fibroblasts alongside cartilage development in a core surrounded by (migrating) fibroblasts compared to one embedded in a cell-free hydrogel. Besides, genes differentially expressed in a core explant surrounded by fibroblasts enriched gene sets related to hypercellularity, ERK1/2 signaling, oxygen / glucose metabolism, and ECM turnover. Most of these processes were reduced in an IL-6 KO core, speculatively because of the reduced presence of extrinsic fibroblasts resulting from the reduced migration and proliferation. On the tissue-level, we also found signs for a decreased catabolic matrix turnover in the more stable mechanical properties of IL-6 KO core explants, a process likely linked to ERK1/2 signaling.^[70]^

In summary, our data consistently point to IL-6 signaling targeting reparative fibroblasts being upregulated in chronic human (non-sheated) tendon lesions in a manner that directly leads to fibroblast recruitment and proliferation as well as aberrant morphogenesis / matrix turnover. This activity contributes to typical hallmarks of tendinopathy including hypercellularity and loss of biomechanical tissue integrity.

## EXPERIMENTAL SECTION

### Human microarray data analysis

We reanalyzed a microarray dataset (GEO: GSE26051) from 2011 with contemporary methods (principal component analysis, volcano plots, heatmaps, GSEA, and ORA), focusing on the Il-6 signaling cascade. All steps from downloading the dataset to the differential expression computation were conducted in RStudio (“Prairie Trillium”, 9f796939, 2022-02-16) running R version 4.1.2. Overall, we closely followed the steps described here: https://sbc.shef.ac.uk/geo_tutorial/tutorial.nb.html (last visited: 02.05.22). First, we log_2_-transformed the expression values and checked their distribution with boxplots. Since the original dataset was gathered from a wide variety of anatomical locations and differently aged patients (**Supplementary Table 1**), we started with a principal-component analysis to filter-out outliers. Based on the clustering, we identified differences between sheathed and non-sheathed tendons (**Supplementary Figure 2**). For the following analysis, we there excluded samples gathered from sheathed tendons:

GSM639749 (EDC), GSM639751 (Flexor-Pronator), GSM639756 (Flexor-Pronator), GSM639761 (ECRB), GSM639765 (ECRL), GSM639772 (ECRB), GSM639774 (Flexor-Pronator), GSM639779 (Flexor-Pronator), GSM639784 (ECRB), GSM639788 (ECRB).

To improve the power to detect differentially expressed genes, we filtered-out genes with very low expression. We considered 50% of genes to not be expressed and therefore used the median expression as the cut-off. In addition, we only kept genes expressed in more than 2 samples for further analysis and calculated the average of replicated probes. Afterwards, we applied the empirical Bayes’ step to receive the differential expression values and p-values. We plotted the differentially expressed genes (DEGs) as a volcano plot and annotated IL-6 signaling-related genes, other cytokines of the IL-6 family, their respective receptors, and genes involved in matrix turnover. Here, we considered genes with a p-value < 0.05 to be differentially expressed. In addition, we plotted the row scaled z-scores of a selection of the annotated genes in a heatmap.

To produce the GO annotations, we fed the list of IDs from differentially expressed genes into the enrichGO function from the clusterProfiler package (version 3.0.4, https://www.rdocumentation.org/packages/clusterProfiler/versions/3.0.4/topics/enrichGO, last visited: 31.10.22) using the org.Hs.eg.db as reference and the Benjamini-Hochberg method to calculate the false discovery rate / adjust the p-values. To visualize the data, we used the dotplot function from the enrichPlot package (https://rdrr.io/bioc/enrichplot/, last visited: 31.10.22). We also looked at the increased (logFC > 0) and decreased (logFC < 0) transcripts in isolation to estimate their contribution to the enrichment and give it directionality.

For the gene set enrichment analysis (GSEA) performed in RStudio with clusterProfiler, we used the human hallmark and the cell type signature gene set annotations from the molecular signature database (MSigDB, https://www.gsea-msigdb.org/gsea/msigdb/, last visited: 31.10.22) after ranking the genes according to their p-value. We used the following input parameters: pvalueCutoff = 1.00, minGSSize = 15, maxGSSize = 500, and eps=0. Lastly, we used the gseaplot and dotplot functions from the enrichPlot package to plot the data and the sign of the enrichment score / NES to estimate the directionality. The exact code can be found in the supplementary material (**Supplementary Material 15**).

### Human immunohistological stainings

Tendon tissues from tendinopathic and normal control tendons were gathered from human patients (Patient data and images are depicted in **Supplementary Figure 5**). We cut transversal cryosections (10 µm thickness) using a low-profile microtome blade (DB80 LX, BioSys Laboratories), collected them on a glass slide, and let them dry for 1h before storing them at −80°C until further use. Prior to staining, sections were air-dried for 30 min at RT (room temperature) and washed 3x with PBS for 5 min each. Then, sections were permeabilized and blocked with 3% BSA (bovine serum albumin) in PBS-T (PBS + 0.1% TritonX) for 1 h at RT. We washed the sections again, added the primary antibody for CD90 (Abcam, ab181469, diluted 1:100 for the co-staining with IL6R and GeneTex, GTX130072 diluted 1:200 for the co-staining with IL-6), CD68 (Abcam, ab955, diluted 1:50), IL-6 (R&D Systems, MAB2061R, diluted 1:200), and IL6R (Absolute Antibody, ab00737-23.0, diluted 1:100) in PBS-T with 1% BSA. We covered them with parafilm and left them overnight in a humid chamber at 4°C. Afterwards, we washed them again (3x 5 min with PBS) before adding the matching secondary antibodies (diluted 1:200 in PBS with 1% BSA) to the samples as well as the secondary antibody controls.

The sections were then washed again (3x 5 min with PBS + 1x 5 min with ultra-pure water) before mounting the coverslip with ROTI^®^Mount FluorCare DAPI (Roth). We used the Leica SP8 automated inverse confocal laser scanning microscope for acquiring the images, which we then processed with ImageJ 1.53q and RStudio (**Supplementary Material 16** and **17).**

### Mouse tissue harvest

We extracted tail tendon core explants and Achilles tendons from 12- to 15-week-old male and female Tgf(Scx-GFP)1Stzr and B6.129S2-Il6tm1Kopf/J Il-6^-/-^ mice (knock-out: KO, wildtype: WT) as described previously (**Figure 4B & C**).^[54,67]^ All experiments were approved by the responsible authorities (Canton Zurich license number ZH104-18 & ZH058-21).

We isolated the core explants from the tail and only kept those with a mean diameter between 100 and 150 µm in standard culture medium (DMEM/F12 GlutaMAX with 10% fetal bovine serum, 1% Penicillin / Streptomycin, 1% Amphotericin, 200 µM L-Ascorbic Acid) until clamping them. Meanwhile, we separated the Achilles tendon from the calcaneus and the calf muscle using a scalpel and washed them with PBS before starting the digestion process (Standard culture medium without L-Ascorbic Acid but 2 mg/ml collagenase for 24h at 37°C). After digestion, we cultured the cells on 2D tissue culture plastic in standard culture medium and used the resulting mixed fibroblast population between passage 2 and 4 (**Supplementary Figure 6**). All medium components were purchased from Sigma Aldrich, except for the ascorbic acid (Wako Chemicals) and the collagenase (ThermoFisher).

### Collagen isolation

We isolated collagen-1 from rat tail tendon fascicles following an established protocol.^[84]^ Briefly, tendon explants were extracted from the tail of adult (> 8 weeks) female Sprague Dawley rats with surgical clamps. Then, the collagen was dissolved by sequentially putting the core explants into acetone (5 min), 70% isopropanol (5 min), and finally 0.02 N acetic acid (48 h). The resulting viscous solution was mixed in a house-ware blender and then frozen at −20°C. Lyophilization at - 20°C turned the viscous solution into a dry collagen sponge, which was stored at −80°C and aliquots thawed when needed. Upon thawing, the collagen aliquot was mixed with 0.02 N acetic acid and then centrifuged (15,000 rpm for 45 min) at 4°C. The supernatant was then sterilized with SPECTRAPOR dialysis bags first in non-sterile acetic acid (1 h), then 1% chloroform in ddH_2_O (1 h), and finally sterile acetic acid (three times for 2 d each). The concentration of the resulting solution was determined with a hydroxyproline assay (Sigma-Aldrich, MAK008), the purity was assessed with SDS-page and Western blots, and the solution itself was stored at 4°C until usage in the experiments.

### Hydrogel preparation, core explant embedding, and assembloid culture

As described previously,^[54,67]^ core explants were fixated with clamps, placed into molds lining silicone chambers, and tensioned. These molds were then filled with cell-free or extrinsic fibroblast-laden collagen hydrogels. One hydrogel consisted of 10 µL PBS (20x), 1.28 µL of 1M NaOH (125x), 8.72 µL double-distilled water (ddH_2_O, 23x), 80 µL collagen-1 (2.5x or 1.6 mg / ml) and 100 µL culture media or cell suspension (2x). All hydrogel components were kept on ice to prevent pre-mature crosslinking. Co-culture medium (DMEM/F12 high glucose, 10% FBS, 1% non-essential amino acids, 1% Penicillin / Streptomycin, 1% Amphotericin, 200 µM L-Ascorbic Acid, 20 ng/ml macrophage-colony stimulating factor) was added to stable hydrogels after 50 min of polymerization at 37°C and tension was released. The assembloids were then cultured under tendinopathic niche conditions (37°C, 20% O_2_) with two media changes per week until the determined timepoint.^[71]^ We used a final concentration of 25 ng/ml recombinant IL-6 (PeproTech, 216-16) in those assembloids to be stimulated by it, and a final concentration of 10 µg/ml Tocilizumab (TargetMol, T9911) in those assembloids to be inhibited by it.

### RNA isolation for genome-wide RNA sequencing (bulk RNA-seq)

We pooled 20 – 24x 2 cm core explants and 2 extrinsic fibroblast-laden collagen hydrogels separate from each other and snap-froze them in liquid nitrogen. The core explant pools were generated from a single mouse and represent one biological replicate each. The collagen hydrogel pools contained a mixed population comprising migratory cells from the embedded core (same mouse) and the initially seeded mixed fibroblast population (cells pooled from 6 mice). The frozen samples were pulverized by cryogenic grinding (FreezerMill 6870, SPEX^TM^SamplePrep) and further processed with the RNeasy micro kit (Qiagen) according to the manufacturer’s instructions. We used the Nanodrop 1000 spectrophotometer 3.7.1 (ThermoFisher) to measure RNA concentration and purity, and the 4200 TapeStation System (Agilent) to measure RNA quality. For each condition (WT core // cell-free, WT core // fibroblasts, KO core // fibroblasts), all 6 of the collagen hydrogels pools but only 4 of the core explant pools passed both integrity control (RIN ≥ 2) and had a sufficiently high RNA concentration (30 – 100 ng / µl) for genome-wide RNA sequencing.

We submitted those pools to the functional genomics center Zurich (https://fgcz.ch/, last visited 06.05.22) for the Illumina (Novaseq 6000) TruSeq TotalRNA stranded sequencing protocol including library construction from total RNA using ribo-depletion, library QC, sequencing, and data delivery.

### RNA-seq data processing and bioinformatic analysis

We used the R-based SUSHI framework of the Functional Genomics Center Zurich (ETH Zurich and University of Zurich) to perform primary level bioinformatics. Specifically, we used the FastqcApp, the FastqScreenApp, and the RnaBamStatsApp for quality control, the KallistoApp (sleuth) to calculate transcript abundance after pseudoalignment, the CountQCApp to quality control after counting reads, and the DESeq2App for differential expression analysis. We then used the shiny toolset developed by the Functional Genomics Center Zurich (https://github.com/fgcz/bfabricShiny, last visited 06.05.22) based on b-fabric and R to generate the annotated volcano plots, heatmaps, and gene set functional enrichment by applying the hypergeometric overrepresentation analysis (ORA) with the following settings:

| **Comparison** | **P-value** | **P-value threshold** | **Log2 fc threshold** |
| --- | --- | --- | --- |
| Core of WT core // cell-free –<br>Core of WT core // fibroblasts | FDR-adjusted | 0.05 | 0.5 |
| Core of WT core // fibroblasts –<br>Core of KO core // fibroblasts | Raw | 0.05 | 0.5 |
| Fibroblasts from WT core // fibroblasts –<br>Fibroblasts from KO core // fibroblasts | Raw | 0.05 | 0.5 |

Heatmap:
| **Comparison** | **#Genes** | **Scale** | **Count method** |
| --- | --- | --- | --- |
| WT core // cell-free –<br>WT core // fibroblasts | Top 50<br>up & down | Diverging | Normalized + Log2 |
| Core of WT core // fibroblasts –<br>Core of KO core // fibroblasts | Top 50<br>up & down | Diverging | Normalized + Log2 |
| Fibroblasts from WT core // fibroblasts –<br>Fibroblasts from KO core // fibroblasts | Top 50<br>up & down | Diverging | Normalized + Log2 |

Overrepresentation analysis:
| **Comparison** | **Input p-value** | **Output p-value** |
| --- | --- | --- |
| WT core // cell-free –<br>WT core // fibroblasts | FDR-adjusted < 0.01 | FDR-adjusted < 0.05 |
| Core of WT core // fibroblasts –<br>Core of KO core // fibroblasts | Raw < 0.01 | FDR-adjusted < 0.05 |
| Fibroblasts from WT core // fibroblasts –<br>Fibroblasts from KO core // fibroblasts | Raw < 0.01 | FDR-adjusted < 0.05 |

We also looked at the increased (logFC > 0) and decreased (logFC < 0) transcripts in isolation to estimate their contribution to the enrichment and give it directionality. The emapplot of enriched biological processes independent of increased and decreased transcripts was generated in RStudio using the enrichPlot package.

We again used RStudio and the clusterProfiler package to perform the GSEA, taking the mouse hallmark and the cell type signature gene set annotations from the molecular signature database (MSigDB, https://www.gsea-msigdb.org/gsea/msigdb/, last visited: 31.10.22) as reference after ranking the genes according to their signed log2 ratio. We used the following input parameters: pvalueCutoff = 1.00, minGSSize = 15, maxGSSize = 500, and eps = 0. Lastly, we used the gseaplot and dotplot functions from the enrichPlot package to plot the data and the sign of the enrichment score / NES to estimate the directionality.

The RNA sequencing data gathered from assembloids as discussed in this publication have been deposited in NCBI’s Gene Expression Omnibus^[85]^ and are accessible through GEO series accession number GSE214015 (https://www.ncbi.nlm.nih.gov/geo/query/acc.cgi?acc=GSE214015).

The *in vivo* mouse RNA sequencing data comparing paratenon-derived to tendon proper-derived fibroblasts have previously been published in an open access database (PRJNA399554, https://www.ncbi.nlm.nih.gov/bioproject/?term=PRJNA399554).^[21]^ To reanalyze this data, we used the same tools and parameters as for the assembloid analysis (GO input: FDR-adjusted p-value < 0.01). The overlapping DEGs and GO terms were calculated and the resulting Venn-diagrams plotted with basic RStudio functions (i.e intersect & draw.pairwise.venn).

### Quantifying total cell proliferation, ScxGFP^+^ cell proliferation, and ScxGFP^+^ cell recruitment to WT and KO core explants

In the assembloids used here, core explants from wildtype (WT) and homozygous (KO) B6.129S2-Il6tm1Kopf/J Il-6^-/-^ mice were embedded with ScxGFP^+^ fibroblasts from homo- and heterozygous Tgf(Scx-GFP)1Stzr mice. After seven days, the assembloids were removed from the clamps and washed with PBS before staining them with Ethidium Homodimer (EthD-1, Sigma-Aldrich, 2 mM stock in DMSO) diluted to 4 µM with PBS (20 min, 37°C). They were then again washed with PBS, fixated with 4% formaldehyde (Roti-Histofix, Karlsruhe) for 1 hour at room temperature, washed again with PBS, and stored in PBS at 4°C. Immediately before the imaging, nuclei were stained with NucBlue Live Ready Probes Reagent^TM^ (R37605, ThermoFisher) for 1 hour at room temperature. We used the Nikon Eclipse T*i*2 confocal scanning microscope controlled by NIS-Elements to acquire the images (3 per sample), which we then processed with ImageJ 1.53q. Briefly, we first registered all cell locations by creating a mask from the NucBlue channel. Then, we put this mask over the ScxGFP-channel and measured the fluorescence intensity at the identified cell locations. We then transferred the signal intensity per location data to RStudio, where we first calculated the total cell numbers of all the images of one sample combined and normalized them to the wildtype median. Afterwards, we determined the fluorescence threshold for the ScxGFP-signal (using density plots and a negative control image) and applied this threshold to the dataset. We then calculated the total ScxGFP^+^ cell numbers for each sample and normalized them to the wildtype median. Finally, we combined the cell location with the fluorescence intensity data to find the distance from the core where most of the ScxGFP^+^ cells were located and to calculate the ratio between ScxGFP^+^ present at the core and those present in the surrounding extrinsic hydrogel (Code: **Supplementary Material 18** and **Supplementary Material 19**).

### Quantifying mechanical properties of assembloids

We mounted the assembloids to a custom-made uniaxial stretching device equipped with a load cell as described previously^[54]^. After five cycles of pre-conditioning to 1% L_0_, the assembloids were then stretched up to 2% L_0_ to measure the linear elastic modulus (Emod) with a pre-load of 0.03 N. This measurement was repeated after 21 days (d21) of culture. We used Matlab R2017a and RStudio to read-out the linear elastic modulus and normalize it to the measurement immediately after assembloid fabrication (d0). Media was changed every 2-3 days. For the corresponding condition, the core explants were devitalized by snap-freezing them repeatedly in liquid nitrogen.

### Achilles tenotomy

Adult wildtype and homozygous *B6.129S2-Il6tm1Kopf/J* IL-6^-/-[69]^ x Tg(Scx-GFP)1Stzr mice (between 12- and 15-weeks old) of both genders were anesthetized by isoflurane inhalation. While the mice were anesthetized, we transected the Achilles tendon of the right hindlimb by creating a small incision in the tendon midsubstance (**Figure 7A**). The contralateral hindlimb was used as the undamaged control. After the surgical intervention, we closed the skin wound with an 8/O prolene suture (Ethicon, W8703) and administered an analgesic (Buprenorphine, 0.1 mg/kg s.c., 26G needle). At one-week (**Supplementary Figure 9** and **Supplementary Figure 10**) and two-weeks (**Figure 7**) post-tenotomy, we injected 10 µl/g of 5-Ethynyl-2-deoxyuridine (EdU) into each mouse of the three-weeks group and euthanized them 24h later with CO_2_. We collected the plantaris and Achilles tendon/ neotendon from both hindlegs for histology. The isolated tissues were placed in OCT (TissueTek), cooled down on dry ice, and then stored at −80°C until further use.

### Immunofluorescence microscopy of mouse Achilles tendon sections

We cut transversal (one-week) and longitudinal (3-weeks) cryosections (10 µm thickness) using a low-profile microtome blade (DB80 LX, BioSys Laboratories), collected them on a glass slide, and let them dry for 1h before storing them at −80°C until further use. Prior to staining, sections were air-dried for 30 min at RT (room temperature) and washed 3x with PBS for 5 min each. Then, sections were permeabilized and blocked with 3% BSA (bovine serum albumin) in PBS-T (PBS + 0.1% TritonX) for 1h at RT. We then washed the sections again and incubated the sections that were previously stained with EdU with a reaction cocktail (Jena Bioscience, CLK-074, CuAAC Cell Reaction Buffer Kit (THPTA based)) prepared according to the manufacturer’s instructions (440 µl reaction buffer, 10 µl CuSo_4_, 1 µl (2 µM) Alexa Fluor azide 647, and 50 µl reducing agent) at RT for 45 min. We washed the sections again, added the primary antibody for Scx (abcam, ab58655, diluted 1:200 in PBS-T with 1% BSA), TPPP3 (Invitrogen, PA5-24925, 1:200), or CD146 (BIOSS, bs-1618R, 1:200) to the sections from the one-week timepoint, and a GFP-antibody (Abcam, ab290, 1:500 in PBS-T with 1% BSA) to those from the three-week timepoint. We covered all the sections with parafilm and left them overnight in a humid chamber at 4°C. Afterwards, we washed them again (3x 5 min with PBS) before adding the matching secondary antibodies (diluted 1:200 in PBS with 1% BSA) to the samples as well as the secondary antibody controls.

The sections were then washed again (3x 5 min with PBS + 1x 5 min with ultra-pure water) before mounting the coverslip with ROTI^®^Mount FluorCare DAPI (Roth). We used the Nikon Eclipse T*i*2 confocal scanning microscope controlled by NIS-Elements for acquiring the images, which we then processed with ImageJ 1.53q and RStudio as described previously (see “Quantifying total cell proliferation, ScxGFP^+^ cell proliferation, and ScxGFP^+^ cell recruitment to WT and KO core explants”). To quantify the migration through the location of Scx^+^ / TPPP^+^ / CD146^+^ cells in **Supplementary Figure 9** and **Supplementary Figure 10**, we defined the lesional Achilles tendon area as a circle with a 480 μm radius set in the center of the Achilles tendon.

### Secretome Analysis

Culture medium was enriched with the secretome of the different assembloids (WT core // cell-free, KO core // cell-free, WT fibroblasts in a hydrogel) for three days and until day 7 of the assembloid/hydrogel culture. IL-6 was quantified using a custom-made multiplex U-PLEX for mouse biomarkers (Meso Scale Discovery) according to the manufacturer’s instruction. Plates were read with the MESO Quickplex SQ120 (Meso Scale Discovery) and analyzed with Discovery Workbench 4.0.13 (https://www.mesoscale.com/en/products_and_services/software). The plate was read with the Epoch Microplate Spectrophotometer (Biotek), and the data were analyzed with R studio.

### Statistical analysis and graph design

Data curation, statistical analysis, and plotting was done in RStudio (“Prairie Trillium”, 9f796939, 2022-02-16) running R version 4.1.2. For normally distributed datasets, statistical information was obtained by ANOVA followed by Tukey Post-Hoc tests for pairwise comparisons. Else, the non-parametric Wilcoxon Rank Sum test was applied, directionally matching the data (less, greater, two-sided). For all tests, we tested the level of p-values. The mean and the standard error of the mean (sem) were reported for the following data: cumulative percentages of ScxGFP^+^ fibroblasts in assembloids, elastic modulus of the assembloids, and cumulative percentages of Scx^+^ / TPPP^+^ / CD146^+^ cells in the *in vivo* tenotomy model. We used bar and/or point plots to depict the mean and error bars/bands to depict the sem. We reported the median and interquartile range (IQR) in assembloids for the total cell number, the number of ScxGFP^+^ cells, and the ratio between core-resident and extrinsic ScxGFP^+^ cells, as well as for the total cell number, the number of Scx^+^ cells, and the ratio between Achilles and neotendon-resident Scx^+^ / TPPP^+^ / CD146^+^ cells in the *in vivo* tenotomy. These values were depicted as boxplots with the upper and lower hinges corresponding to the first and third quartile (25^th^ and 75^th^ percentile), the middle one to the median, the whiskers extending from the upper/lower hinge to the largest/smallest value no further than 1.5 times the interquartile range, and dots representing data beyond the whiskers. Results of the statistical analysis are indicated as follows: *p < 0.05, **p < 0.01, ***p < 0.01.

The open-source graphics software Inkscape 0.92.3 (www.inkscape.org, last visited 09.05.22) was used to finalize the graph design.

## Supporting information

SupplementaryMaterial15to19_AnalysisCode

## ACKNOWLEDGEMENTS

This work was funded by the ETH Grant 1-005733. Declarations of interest: none

We would like to thank the Functional Genomics Center Zurich, and in particular Lennart Opitz and Dr. Maria Domenica Moccia, for their support on the RNA sequencing data analysis and Dr. Roberto Fiore from the System Neuroscience Lab at ETH Zurich for providing the rat tails for the collagen-1 extraction. We further acknowledge Dr. Evi Masschelein for performing the Achilles tenotomies and the Laboratory of Nutrition and Metabolic Epigenetics for the access to their Tapestation. We also thank our own Lab Technicians Barbara Niederöst and Maja Bollhalder for their practical and emotional support. Finally, we thank Dr. Knut Husmann and Dr. Annamari Katariina Alitalo for their help and support with obtaining the license for animal experimentation and animal husbandry.

## SUPPLEMENTARY MATERIAL

**Supplementary Table 1:** Human patient microarray metadata. GEO accession number, patient sex, source tissue, patient age, donor number, and disease state of the isolated tissue ordered by GEO accession number. Samples from sheathed tendons are colored in red and were excluded from further analysis.

| GEO accession number | Sex | Source tissue | Age | Donor number | Disease state |
| --- | --- | --- | --- | --- | --- |
| GSM639748 | Male | Non-sheathed tendon;Brachialis | 45 | 5 | Normal |
| GSM639749 | Male | Sheathed tendon;EDC | 48 | 9 | Normal |
| GSM639750 | Male | Non-sheathed tendon;Quadriceps | 41 | 10 | Normal |
| GSM639751 | Male | Sheathed tendon;Flexor-Pronator | 62 | 12 | Normal |
| GSM639752 | Female | Non-sheathed tendon;Subscapularis | 62 | 13 | Normal |
| GSM639753 | Male | Non-sheathed tendon;Patellar | 32 | 15 | Normal |
| GSM639754 | Female | Non-sheathed tendon;Subscapularis | 61 | 16 | Normal |
| GSM639755 | Male | Non-sheathed tendon;Subscapularis | 59 | 17 | Normal |
| GSM639756 | Male | Sheathed tendon;Flexor-Pronator | 45 | 19 | Normal |
| GSM639757 | Female | Non-sheathed tendon;Subscapularis | 63 | 20 | Normal |
| GSM639758 | Female | Non-sheathed tendon;Biceps | 65 | 21 | Normal |
| GSM639759 | Female | Non-sheathed tendon;Subscapularis | 64 | 23 | Normal |
| GSM639760 | Male | Non-sheathed tendon;Biceps | 50 | 24 | Normal |
| GSM639761 | Male | Sheathed tendon;ECRB | 55 | 26 | Normal |
| GSM639762 | Male | Non-sheathed tendon;Subscapularis | 50 | 27 | Normal |
| GSM639763 | Male | Non-sheathed tendon;Biceps | 41 | 28 | Normal |
| GSM639764 | Male | Non-sheathed tendon;Biceps | 46 | 29 | Normal |
| GSM639765 | Male | Sheathed tendon;ECRL | 52 | 30 | Normal |
| GSM639766 | Male | Non-sheathed tendon;Subscapularis | 66 | 31 | Normal |
| GSM639767 | Female | Non-sheathed tendon;Teres minor | 46 | 32 | Normal |
| GSM639768 | Male | Non-sheathed tendon;Subscapularis | 44 | 33 | Normal |
| GSM639769 | Female | Non-sheathed tendon;Biceps | 59 | 34 | Normal |
| GSM639770 | Female | Non-sheathed tendon;Subscapularis | 49 | 35 | Normal |
| GSM639771 | Male | Non-sheathed tendon;Biceps | 45 | 5 | Tendinopathic |
| GSM639772 | Male | Sheathed tendon;ECRB | 48 | 9 | Tendinopathic |
| GSM639773 | Male | Non-sheathed tendon;Patellar | 41 | 10 | Tendinopathic |
| GSM639774 | Male | Sheathed tendon;Flexor-Pronator | 62 | 12 | Tendinopathic |
| GSM639775 | Female | Non-sheathed tendon;Suspraspinatus | 62 | 13 | Tendinopathic |
| GSM639776 | Male | Non-sheathed tendon;Patellar | 32 | 15 | Tendinopathic |
| GSM639777 | Female | Non-sheathed tendon;Suspraspinatus | 61 | 16 | Tendinopathic |
| GSM639778 | Male | Non-sheathed tendon;Suspraspinatus | 59 | 17 | Tendinopathic |
| GSM639779 | Male | Sheathed tendon;Flexor-Pronator | 45 | 19 | Tendinopathic |
| GSM639780 | Female | Non-sheathed tendon;Suspraspinatus | 63 | 20 | Tendinopathic |
| GSM639781 | Female | Non-sheathed tendon;Suspraspinatus | 65 | 21 | Tendinopathic |
| GSM639782 | Female | Non-sheathed tendon;Suspraspinatus | 64 | 23 | Tendinopathic |
| GSM639783 | Male | Non-sheathed tendon;Suspraspinatus | 50 | 24 | Tendinopathic |
| GSM639784 | Male | Sheathed tendon;ECRB | 55 | 26 | Tendinopathic |
| GSM639785 | Male | Non-sheathed tendon;Suspraspinatus | 50 | 27 | Tendinopathic |
| GSM639786 | Male | Non-sheathed tendon;Suspraspinatus | 41 | 28 | Tendinopathic |
| GSM639787 | Male | Non-sheathed tendon;Suspraspinatus | 46 | 29 | Tendinopathic |
| GSM639788 | Male | Sheathed tendon;ECRB | 52 | 30 | Tendinopathic |
| GSM639789 | Male | Non-sheathed tendon;Suspraspinatus | 66 | 31 | Tendinopathic |
| GSM639790 | Female | Non-sheathed tendon;Suspraspinatus | 46 | 32 | Tendinopathic |
| GSM639791 | Male | Non-sheathed tendon;Suspraspinatus | 44 | 33 | Tendinopathic |
| GSM639792 | Female | Non-sheathed tendon;Suspraspinatus | 59 | 34 | Tendinopathic |
| GSM639793 | Female | Non-sheathed tendon;Suspraspinatus | 49 | 35 | Tendinopathic |

**Supplementary Figure 2:**
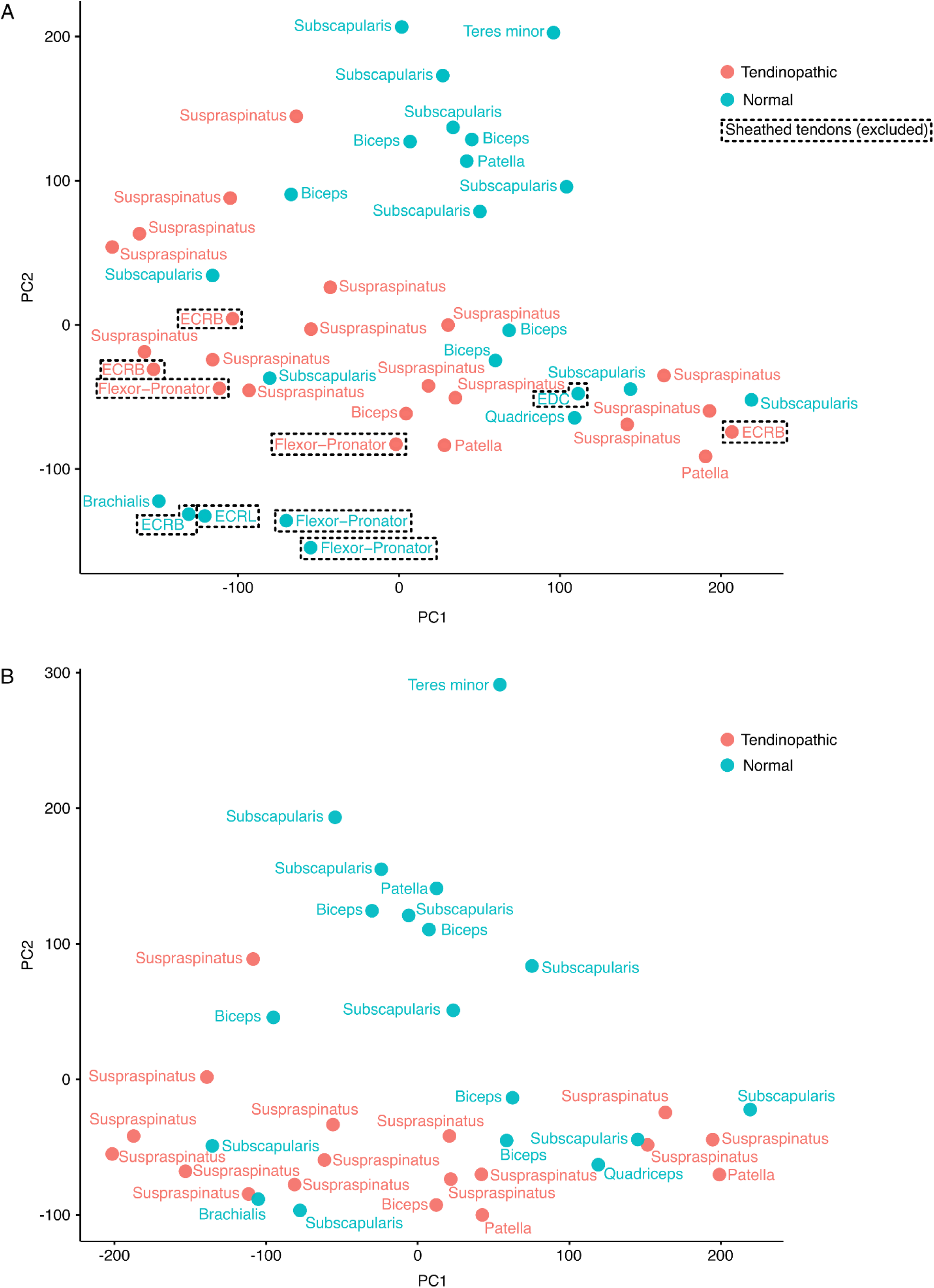
PCA plots of the human tendon microarray data. (A) Principal components 1 and 2 for the full dataset with tendinopathic (red) and normal (blue) tendons. Tendons surrounded by a sheath *in vivo* are delineated by a dashed border. (B) Principal components 1 and 2 for the same dataset after excluding sheathed tendons as delineated in A.

**Supplementary Figure 3:**
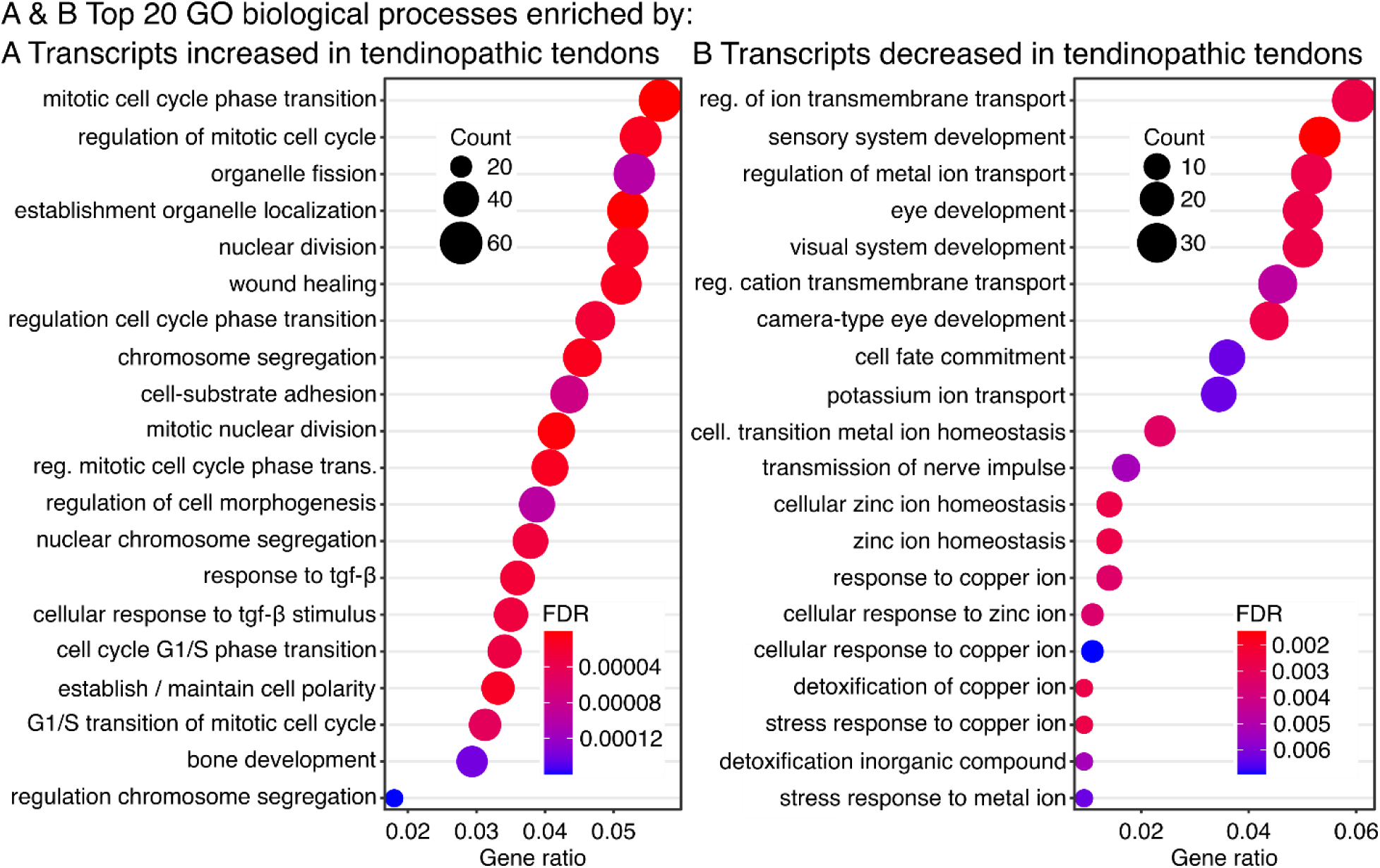
Detailed transcriptome analysis of up- and downregulated genes and pathways in normal and human tendinopathic tendons (non-sheathed) (A) Detailed annotation of the enrichment map plot clustering the top 30 biological processes significantly enriched by overlapping DEG sets from tendinopathic samples compared to normal controls. The color of the circles represents their adjusted p-values, the size represents the number of enriched genes (count), and the grey lines connect GO annotations that share the same gene subsets. (B) Dotplot showing the top 20 GO gene sets for biological processes significantly enriched by transcripts increased in human tendinopathic tendons. (C) Dotplot showing all the GO gene sets for biological processes enriched by transcripts decreased in human tendinopathic tendons. (D) Dotplot showing all the GO gene sets for molecular functions enriched by transcripts increased in human tendinopathic tendons. (E) Dotplot showing all the GO gene sets for molecular functions enriched by transcripts decreased in human tendinopathic tendons. In all the dotplots, the color of the circles represents their adjusted p-value, the size the number of enriched genes (count), and the position on the x-axis the number of enriched genes in ratio to the total number of genes annotated to the gene set (gene ratio).

**Supplementary Figure 4:**
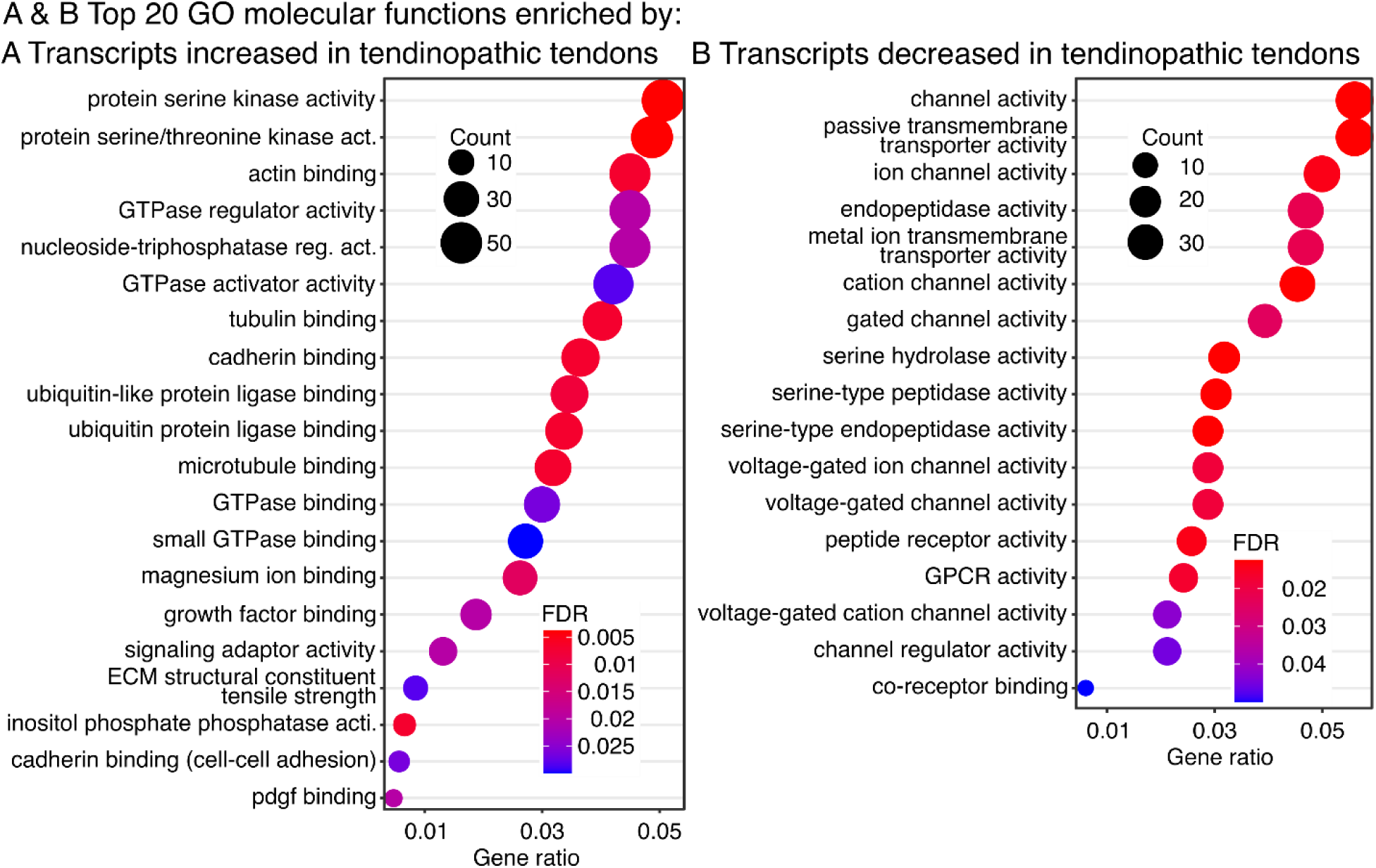
Detailed transcriptome analysis of up- and downregulated genes and pathways in normal and human tendinopathic tendons (non-sheathed) (A) Detailed annotation of the enrichment map plot clustering the top 30 biological processes significantly enriched by overlapping DEG sets from tendinopathic samples compared to normal controls. The color of the circles represents their adjusted p-values, the size represents the number of enriched genes (count), and the grey lines connect GO annotations that share the same gene subsets. (B) Dotplot showing the top 20 GO gene sets for biological processes significantly enriched by transcripts increased in human tendinopathic tendons. (C) Dotplot showing all the GO gene sets for biological processes enriched by transcripts decreased in human tendinopathic tendons. (D) Dotplot showing all the GO gene sets for molecular functions enriched by transcripts increased in human tendinopathic tendons. (E) Dotplot showing all the GO gene sets for molecular functions enriched by transcripts decreased in human tendinopathic tendons. In all the dotplots, the color of the circles represents their adjusted p-value, the size the number of enriched genes (count), and the position on the x-axis the number of enriched genes in ratio to the total number of genes annotated to the gene set (gene ratio).

**Supplementary Figure 5:**
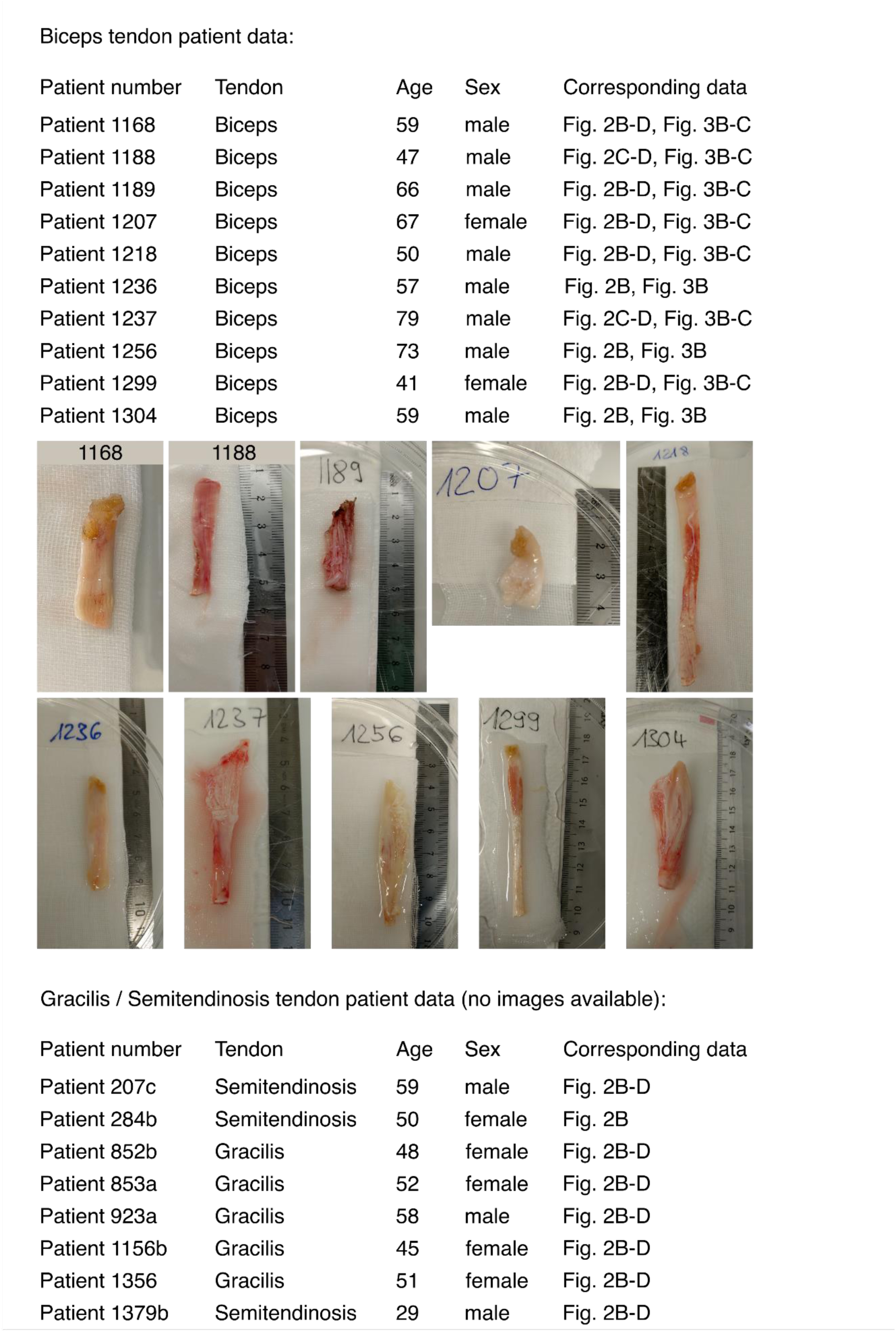
Metadata of the human patient-derived tissues analyzed with fluorescence microscopy. Tendinopathic tissues were gathered from a total of 10 patients with painful biceps tendons. The photographic images depict the tendons from which the sections were collected. Normal control tissues were gathered from the leftover semitendinosus or gracilis tendons of a total of 8 patients undergoing anterior cruciate ligament (ACL) reconstruction surgery. No images were taken from the normal control tendons.

**Supplementary Figure 6:**
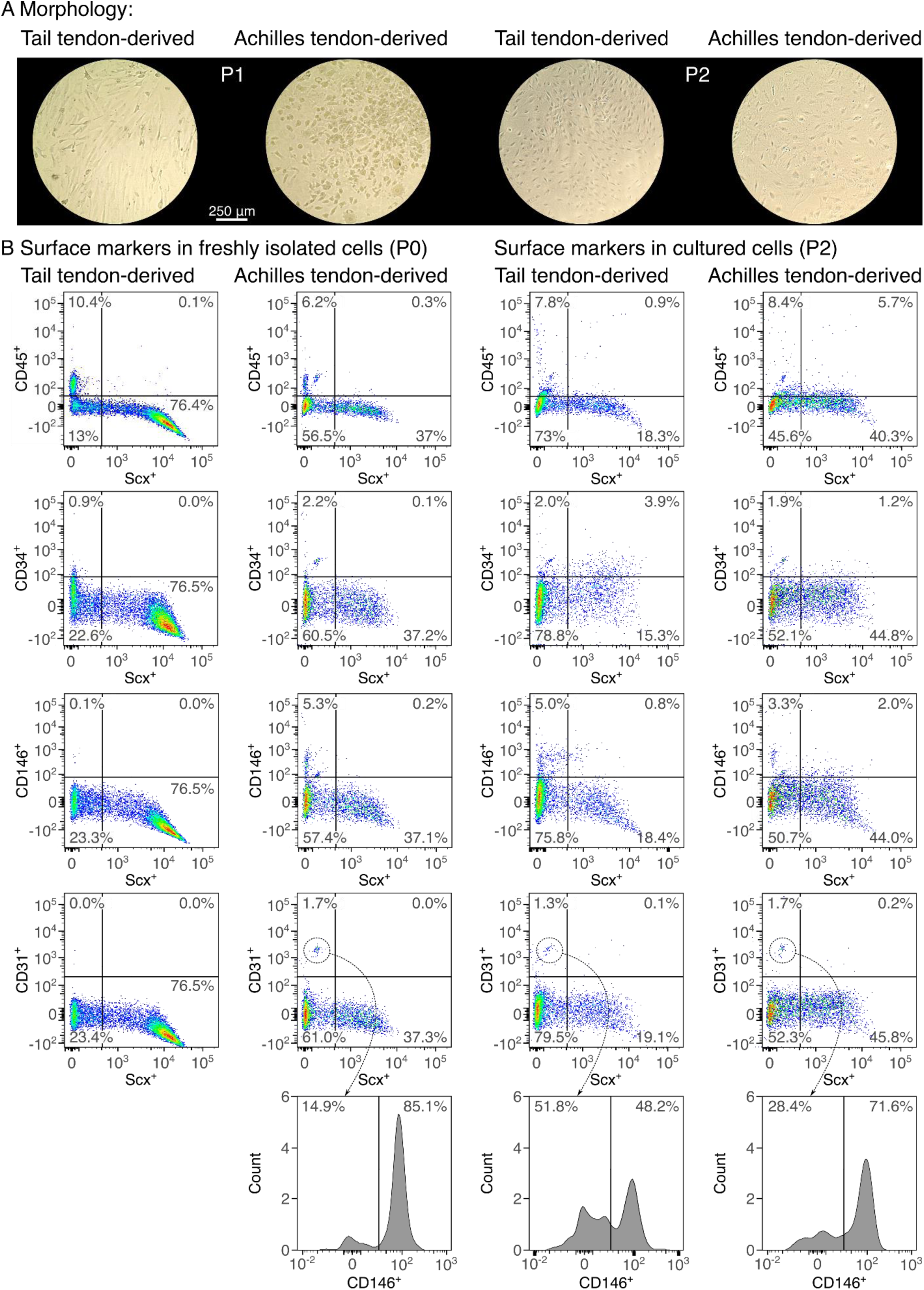
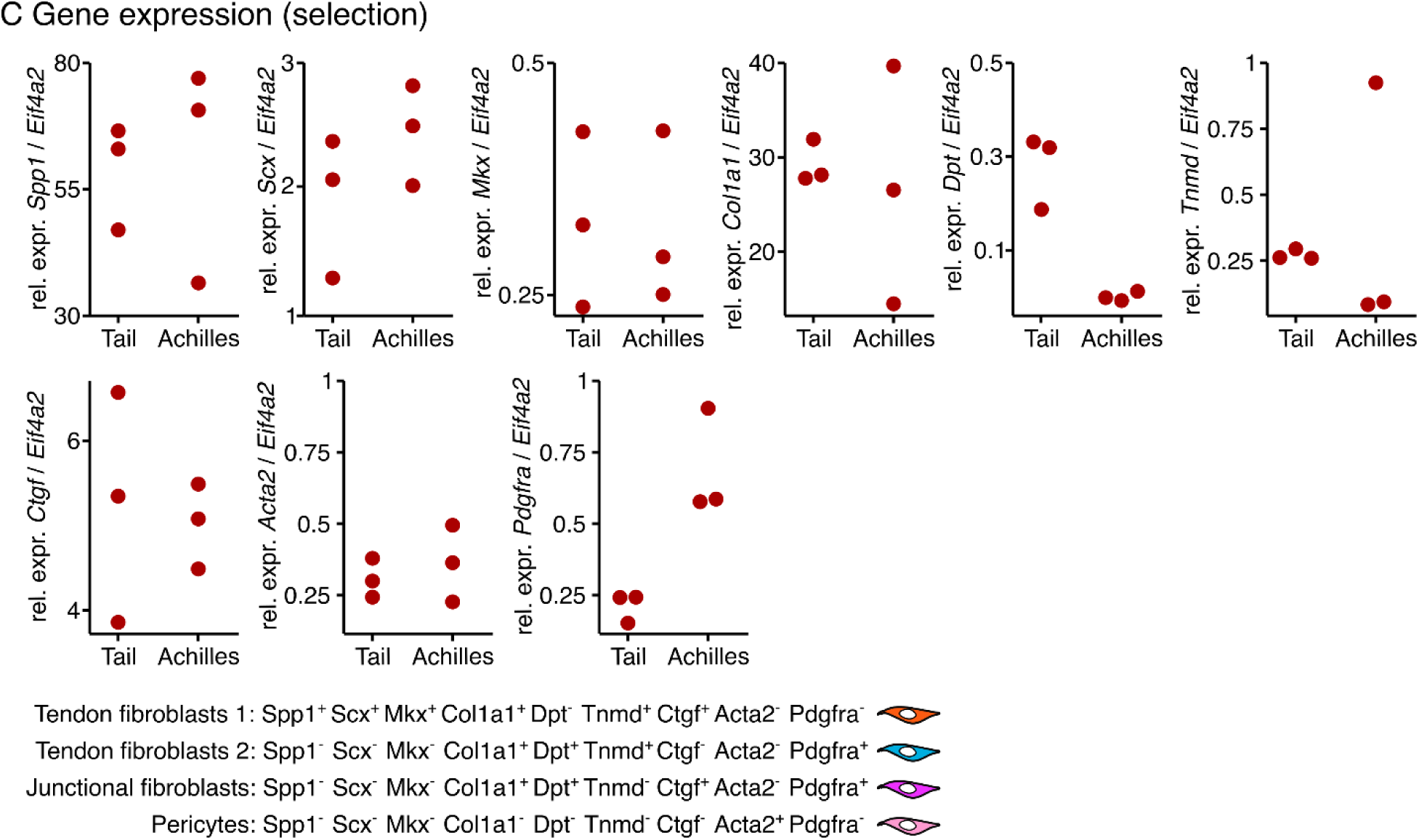
Characterization of cell populations derived from Achilles tendons and tail tendon fascicles of ScxGFP^+^ mice. (A) Representative light microscopy images depicting tail tendon fascicle- derived fibroblasts and Achilles tendon-derived fibroblasts cultured on standard tissue culture plastic before being passaged for the first time (P1) or the second time (P2). The Achilles tendon-derived fibroblasts were embedded in the collagen hydrogels at P2. (B) Presence of a selection of cell surface markers on cell populations isolated from tail tendon fascicles or Achilles tendons immediately after the digestion (left) and after passaging them twice (P2, right). Using flow cytometry, the following markers were analyzed after excluding doublets and dead cells: Scx, CD45, CD34, CD31, and CD136. (C) Presence of a selection of gene transcripts in cell populations isolated from tail tendon fascicles or Achilles tendons after passaging them twice. Using RT-qPCR, the following gene transcripts were measured: *Spp1*, *Scx*, *Mkx*, *Col1a1*, *Dpt*, *Tnmd*, *Ctgf*, *Acta2*, and *Pdgfra*. The dots indicate the relative expression values (dCt) of the transcripts normalized to that of Eif4a2, an established housekeeping gene, for a different animal each. N = 3. The depicted gene transcripts and their typical expression levels in different cell populations were previously reported in mouse Achilles tendons analyzed with single-cell RNA-seq.^[86]^

**Supplementary Figure 7:**
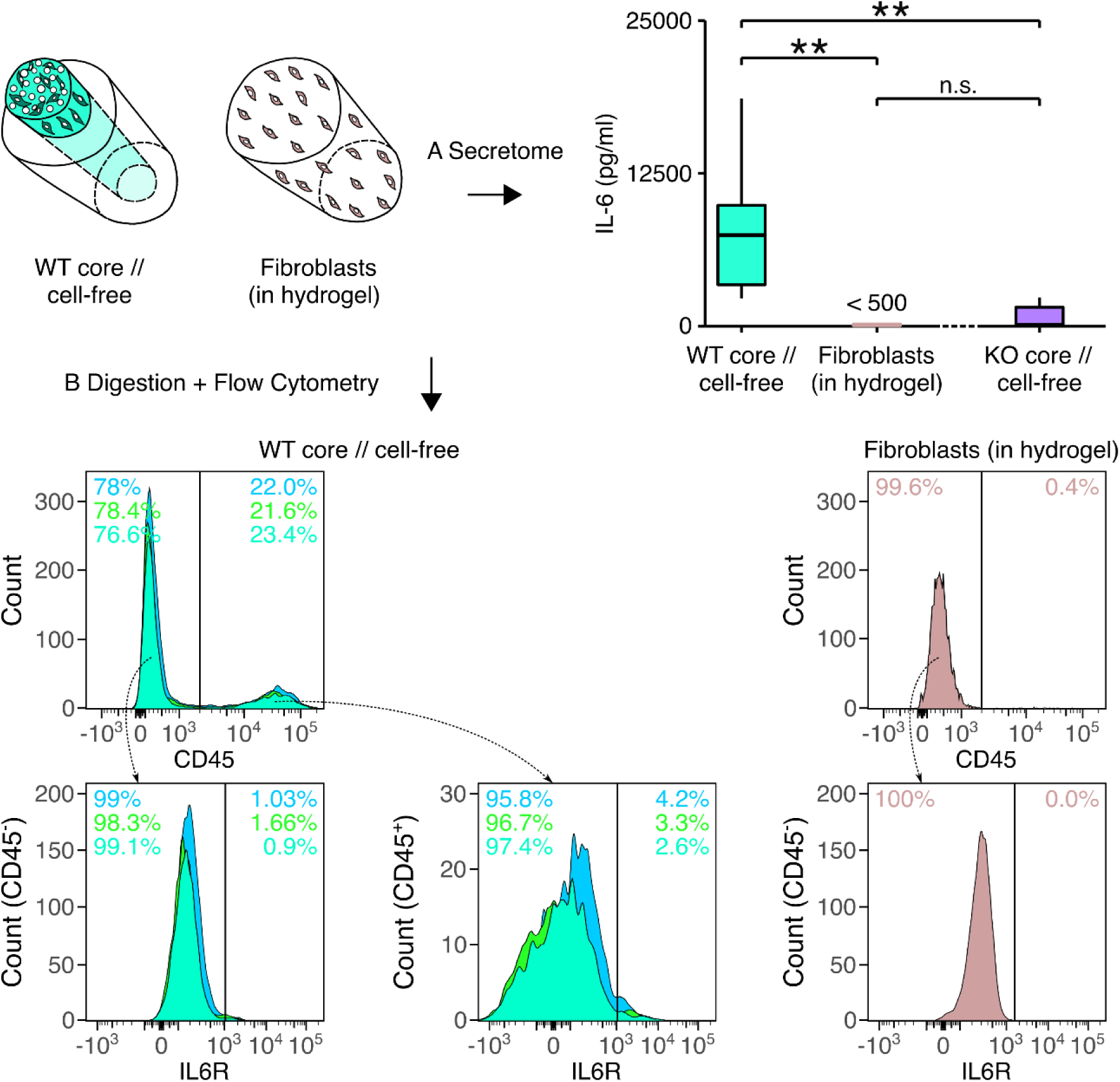
Identification of cellular IL-6 and IL6R sources in mouse tendon assembloids using flow cytometry and analyzing the supernatant. (A) Detection of IL-6 protein level differences in the supernatants obtained from IL-6 wildtype core explant surrounded by a collagen hydrogel (WT core // cell-free, light blue), IL-6 knock-out core explants surrounded by a collagen hydrogel (KO core // cell-free, violet), and extrinsic, IL-6 wildtype fibroblasts seeded into a collagen hydrogel (fibroblasts (in hydrogel), light brown) after 7 days in culture. N = 6. The upper and lower bounding boxes correspond to the first and third quartile (25th and 75th percentile) and the middle bar to the median. Whiskers extend from the upper/lower hinge to the largest/smallest value no further than 1.5 the interquartile range. Results of the statistical analysis are indicated as follows: **p < 0.01, ^n.s.^p >= 0.05. The applied statistical test was the Wilcoxon RankSum. (B) On the left side: Flow cytometric analysis of digested wildtype core explants (WT core // cell-free) surrounded by a collagen hydrogel after 7 days in culture. The different colors indicate different biological replicates (N =3). On the right side: Flow cytometric analysis of extrinsic fibroblast cultured on tissue culture plastic before being passaged for the second time (P2, N = 1). Assessed markers include the hematopoietic lineage marker CD45 and the IL6R.

**Supplementary Figure 8:**
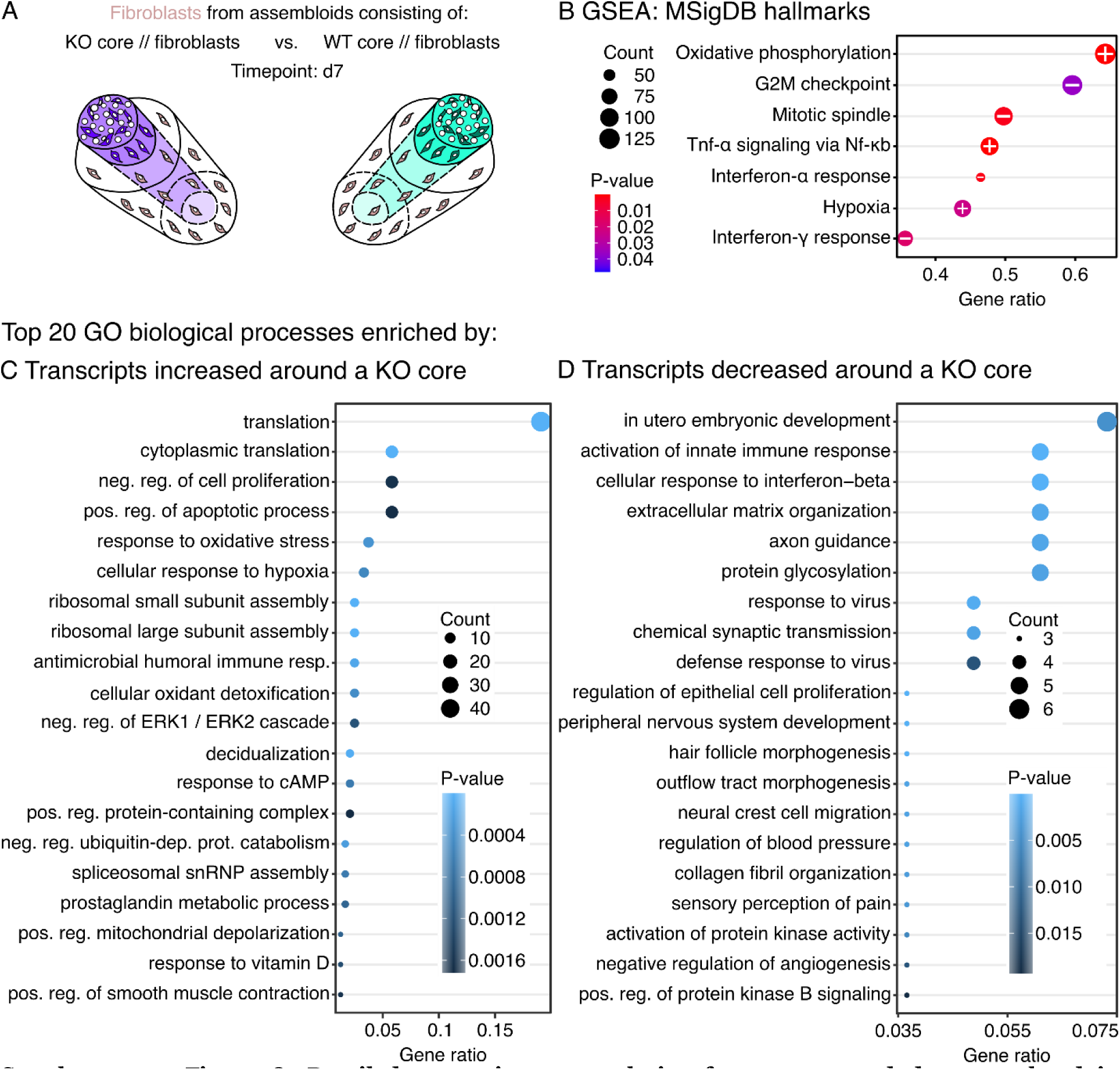
Detailed transcriptome analysis of genes up- and down-regulated in hydrogel-embedded fibroblast seeded around an IL-6 knock-out (KO) core explant compared to those seeded around a wildtype (WT) core. (A) Illustration of the assembloid combinations compared here (KO core // fibroblasts vs. WT core // fibroblasts), the assessed timepoint (d7), and the analyzed compartment (extrinsic fibroblasts only). (B) Dotplot showing significantly enriched gene sets (p-value < 0.05) as determined by GSEA based on the MSigDB mouse hallmark gene sets. The +/- signs indicate the direction of the enrichment in the extrinsic fibroblasts around a KO core compared to those around a WT core. (C) Dotplot showing the top 20 GO gene sets for biological processes significantly enriched by transcripts increased in fibroblasts seeded around a KO core. (D) Dotplot showing the top 20 GO gene sets for biological processes significantly enriched by transcripts decreased in fibroblasts seeded around a KO core. In all the dotplots, the color of the circles represents their p-value, the size the number of enriched genes (count), and the position on the x-axis the number of enriched genes in ratio to the total number of genes annotated to the gene set (gene ratio).

**Supplementary Figure 9:**
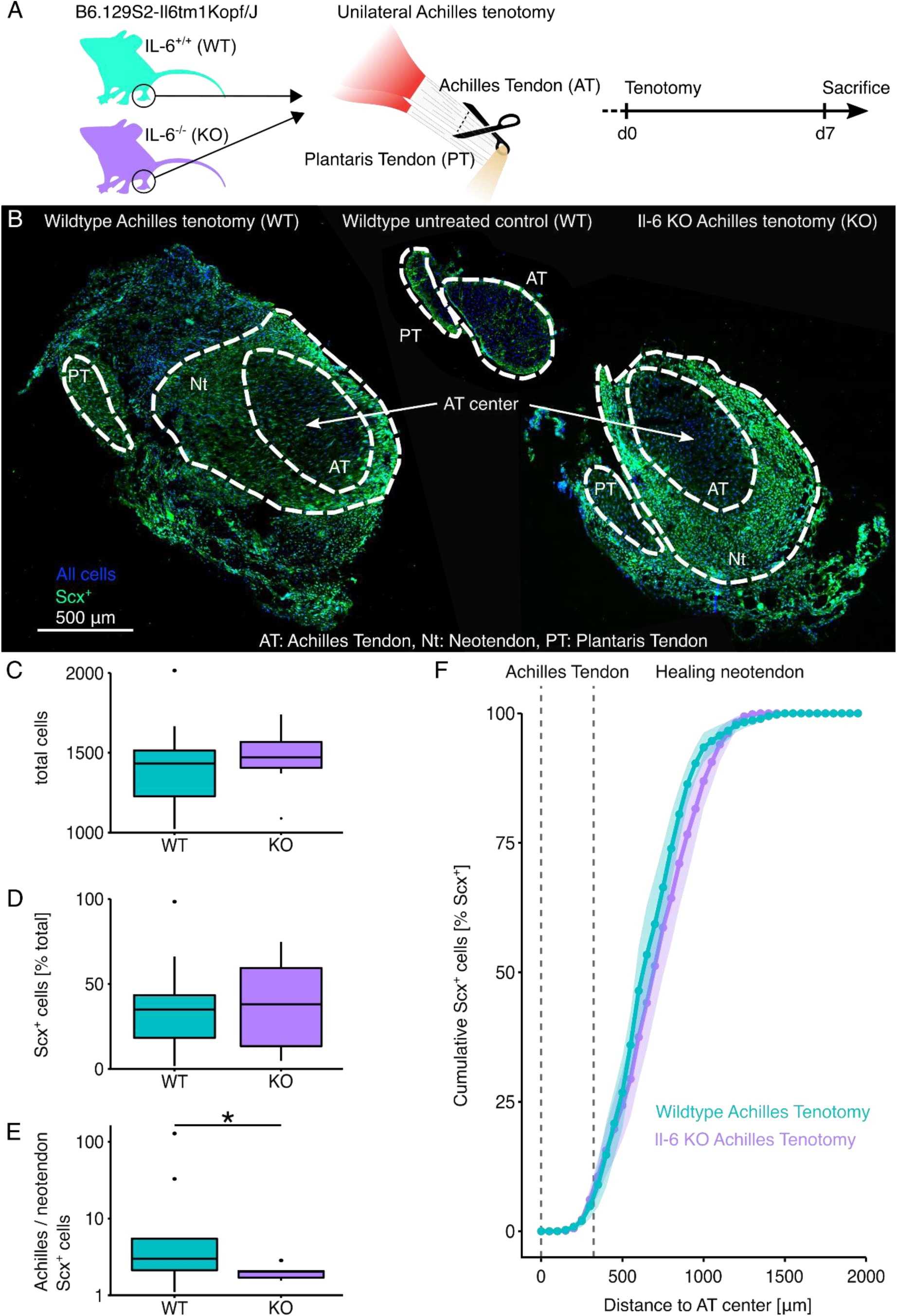
Cell proliferation around and Scx^+^ fibroblast recruitment to damaged mouse Achilles tendons. (A) Illustrative depiction of the experimental setup and the time schedule. (B) Representative fluorescence microscopy images of mouse hindleg cross-sections from wildtype (WT) Achilles tendons that underwent tenotomy (left), the contralateral untreated control (middle), as well as cross-sections from IL-6 knock-out (KO) Achilles tendons that underwent tenotomy (right). (C) Total number of cells stained with NucBlue. (D) Number of Scx^+^ cells. (E) Ratio between core-resident and extrinsic Scx^+^ cells depicted on a logarithmic y-axis. N=7. The upper and lower hinges correspond to the first and third quartile (25th and 75th percentile), the middle one to the median, the whiskers extend from the hinges no further than 1.5 times the interquartile range, and the points beyond the whiskers are treated as outliers. (F) Lineplot depicting the cumulative percentage of Scx^+^ cells depending on their distance from the Achilles tendon center. The points and the line represent the mean cumulative percentages and their error bands the standard error of the mean (sem). The dashed line indicates locations inside the Achilles tendon stump. Results of the statistical analysis are indicated as follows: *p < 0.05. The applied statistical test was the Mann-Whitney-Wilcoxon Test.

**Supplementary Figure 10:**
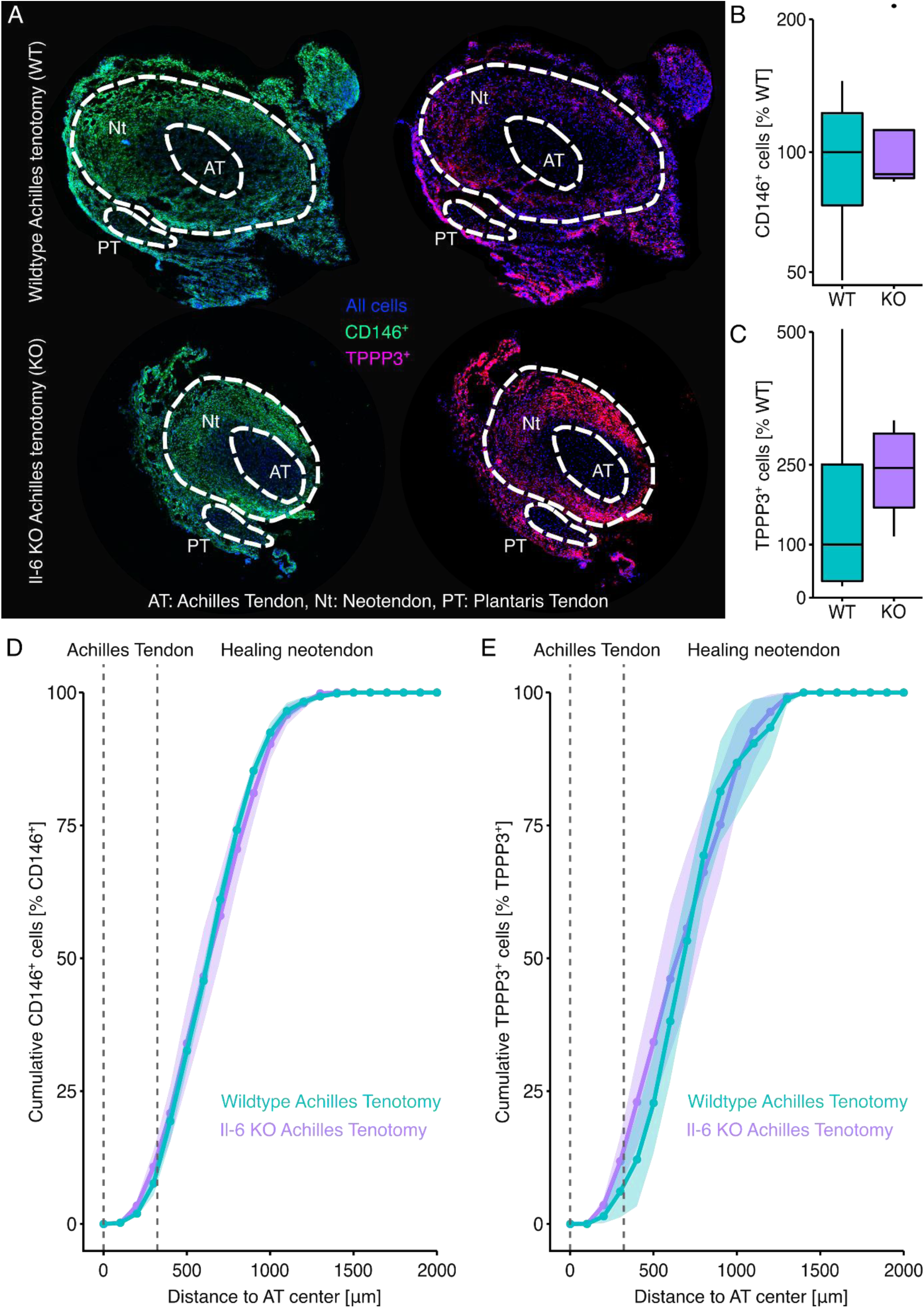
CD146^+^ and TPPP3^+^ fibroblast recruitment to damaged mouse Achilles tendons. (A) Representative fluorescence microscopy images of mouse hindleg sections from wildtype (WT) and IL-6 knock-out (KO) Achilles tendons that underwent unilateral tenotomy. (B) Boxplot reporting the number of CD146^+^ cells normalized to the WT median. (C) Boxplot reporting the number of TPPP3^+^ cells normalized to the WT median. In these boxplots, the upper and lower hinges correspond to the first and third quartile (25th and 75th percentile), the middle one to the median, the whiskers extend from the hinges no further than 1.5 times the interquartile range, and the points beyond the whiskers are treated as outliers. (D, E) Lineplots depicting the cumulative percentages of CD146^+^ and TPPP3^+^ cells depending on their distance from the Achilles tendon center. The points and the line represent the mean cumulative percentages and the error bands their standard error of the mean (sem). The dashed line indicates locations inside the Achilles tendon stump. N=4. The applied statistical test was the non-parametric Wilcoxon Rank Sum Test and no significant differences were detected.

**Supplementary Figure 11:**
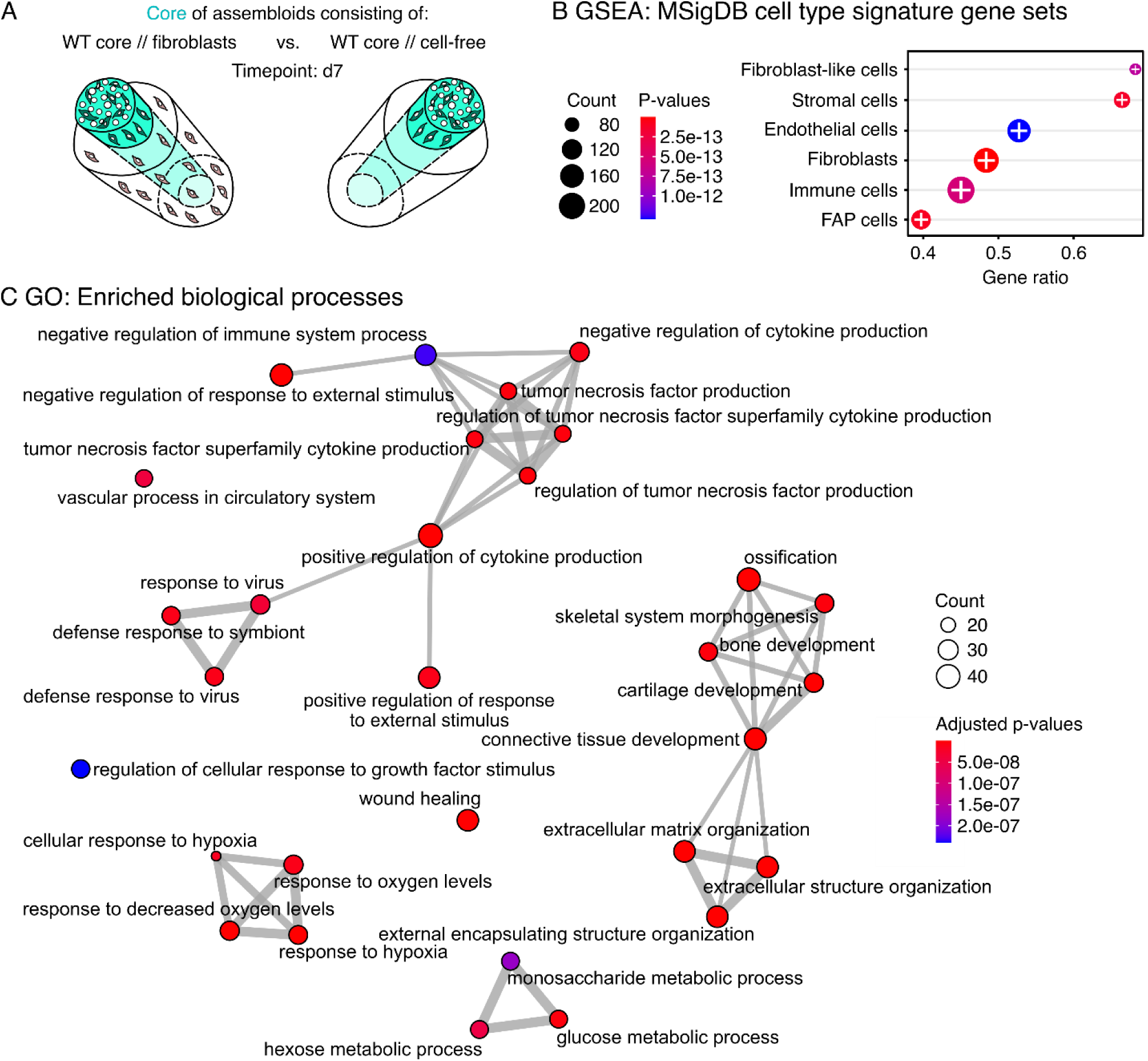
Detailed transcriptome analysis of genes up- and down-regulated in wildtype (WT) core explants surrounded by a hydrogel seeded with extrinsic fibroblasts compared to a WT core surrounded by a cell-free hydrogel. (A) Illustration of the assembloid combinations compared here (WT core // fibroblasts vs. WT core // cell-free), the assessed timepoint (d7), and the analyzed compartment (core only). (B) Dotplot depicting the top 6 significantly enriched gene sets as determined by GSEA based on the MSigDB mouse cell type signature gene sets. The color of the circles represents their p-value, the size the number of enriched genes (count), the position on the x-axis the number of enriched genes in ratio to the total number of genes annotated to the gene set (gene ratio), and the +/- signs the direction of the enrichment in the WT core surrounded by fibroblasts compared to the WT core surrounded by a cell-free hydrogel. (C) Detailed annotation of the enrichment map plot clustering the top 30 biological processes significantly enriched by DEG sets. The color of the circles represents their adjusted p-values, the size represents the number of enriched genes (count), and the grey lines connect GO annotations that share the same gene subsets.

**Supplementary Figure 12:**
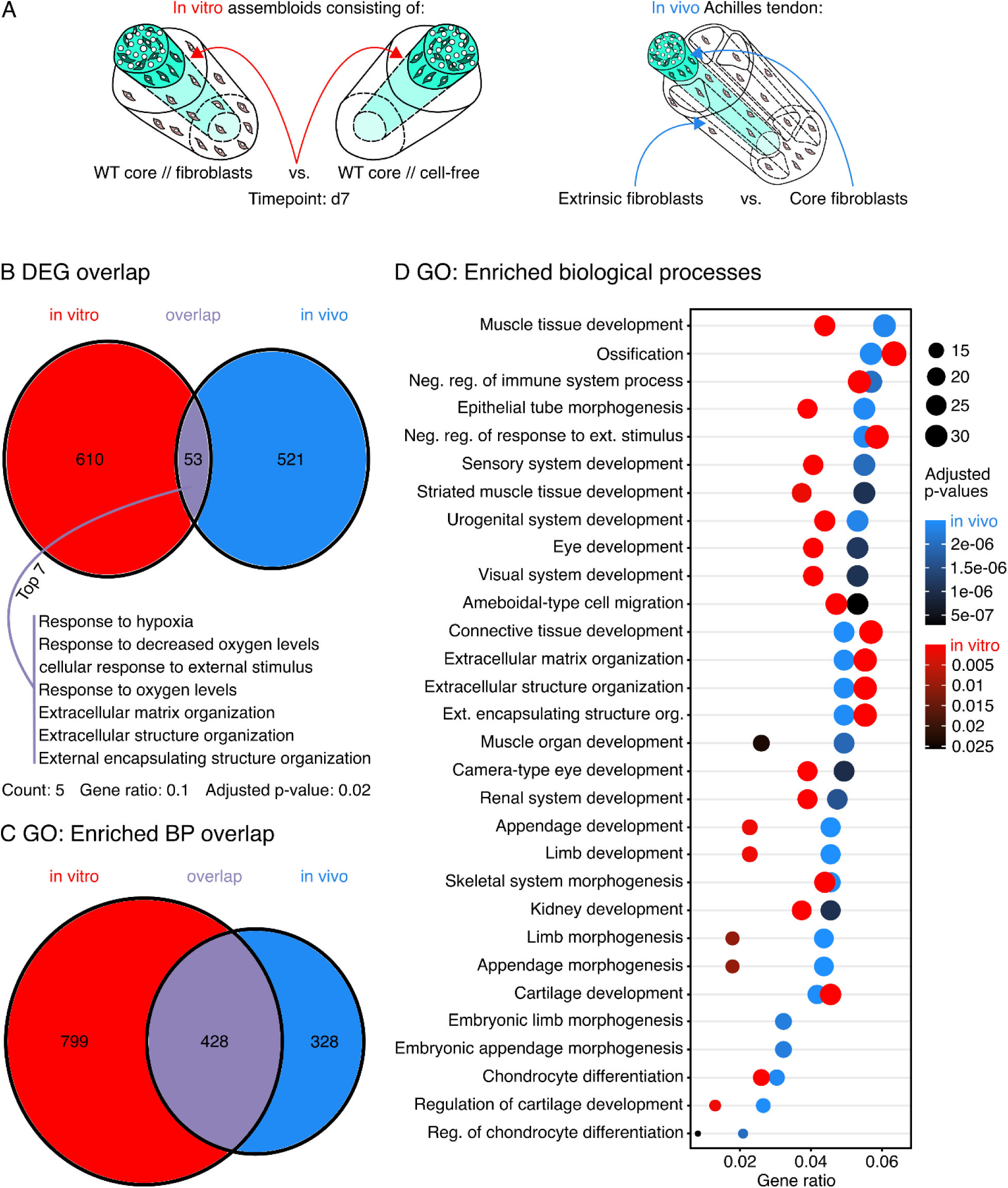
Overlap between differentially expressed transcripts in *in vitro* assembloids and differentially expressed transcripts between Achilles tendon fibroblasts from the extrinsic compartment and the tendon core *in vivo*. (A) Illustrative depiction of the comparisons whose overlap was investigated here: The core explants of WT core // fibroblast vs. the WT core // cell-free assembloids (*in vitro*, red) and the extrinsic (paratenon-derived) vs. the core (tendon proper-derived) fibroblasts (*in vivo*, blue). (B) Venn-Diagramm depicting the number and the overlap (violet) of DEGs as well as the top 7 GO gene sets for biological processes significantly enriched by the overlapping DEGs. (C) Venn-Diagramm depicting the number and the overlap (violet) of significantly enriched GO annotations for biological processes. (D) Dotplot depicting the top 30 biological processes significantly enriched by DEGs in tendon fibroblasts derived from the extrinsic compartment (paratenon-derived) compared to those derived from the core compartment (tendon proper-derived) colored in a blue to black gradient. The plot is augmented by the data of matched biological processes also significantly enriched in the core of WT core // fibroblast compared to that of the WT core // cell-free assembloids colored in a red to black gradient. The color gradient of the circles represents their adjusted p-values, the size the number of enriched genes (count), and the x-axis the number of enriched genes in ratio to the total number of genes annotated to the gene set (gene ratio).

**Supplementary Figure 14:**
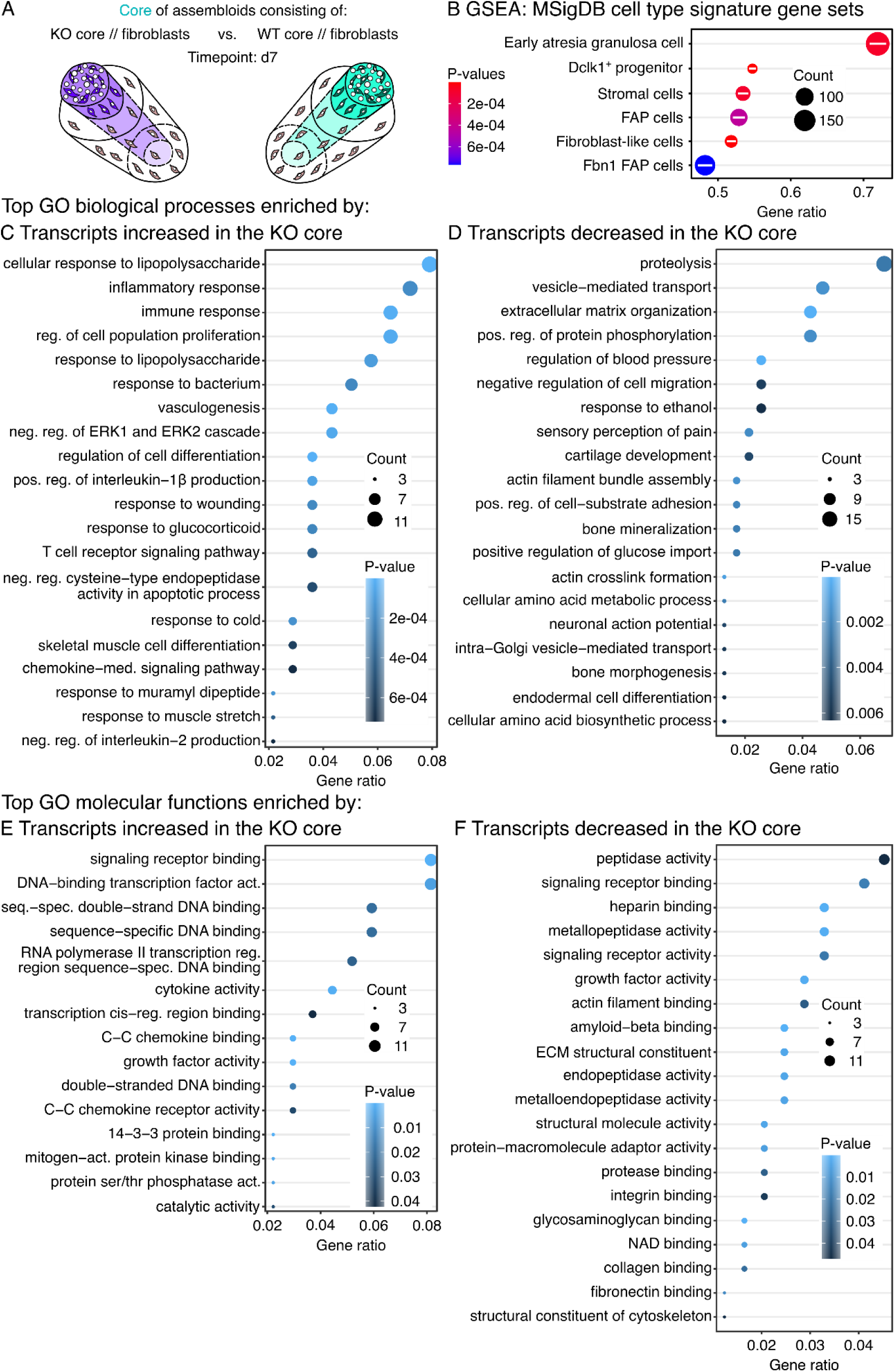
Detailed transcriptome analysis of genes up- and down-regulated in knock- out (KO) core explants surrounded by a hydrogel seeded with fibroblasts compared to a wildtype (WT) core surrounded by a hydrogel seeded with fibroblasts. (A) Illustration depicting the assembloid combinations compared here (KO core // fibroblasts vs. WT core // fibroblasts), the assessed timepoint (d7), and the analyzed compartment (core only). (B) Dotplot depicting the top 6 significantly enriched gene sets as determined by GSEA based on the MSigDB mouse cell type signature gene sets. The +/- signs indicate the direction of the enrichment in the KO core surrounded by fibroblasts compared to the WT core surrounded by fibroblasts. (C) Dotplot showing the top 20 GO gene sets for biological processes significantly enriched by transcripts increased in the core of KO core // fibroblast assembloids. (D) Dotplot showing the top 20 GO gene sets for biological processes significantly enriched by transcripts decreased in the core of KO core // fibroblast assembloids. (E) Dotplot showing the top 20 GO gene sets for molecular functions significantly enriched by transcripts increased in the core of KO core // fibroblast assembloids. (F) Dotplot showing the top 20 GO gene sets for molecular functions significantly enriched by transcripts decreased in the core of KO core // fibroblast assembloids. In all the dotplots, the color of the circles represents their p-value, the size the number of enriched genes (count), and the position on the x-axis the number of enriched genes in ratio to the total number of genes annotated to the gene set (gene ratio).

**Supplementary File 15: R code file used for the human microarray analysis.**

**Supplementary File 16: ImageJ code file used for the analysis of histological sections from humans.**

**Supplementary File 17: R code file used for the analysis of histological sections from humans.**

**Supplementary File 18: ImageJ code file used for the analysis of histological sections from assembloids.**

**Supplementary File 19: R code file used for the analysis of histological sections from assembloids.**

