## SupplementaryMaterial15to19_AnalysisCode for "IL-6 SIGNALING EXACERBATES HALLMARKS OF CHRONIC TENDON DISEASE BY STIMULATING REPARATIVE FIBROBLASTS": SupplementaryMaterial15_HumanMicroArrayAnalysis.R

```
if (!require("BiocManager", quietly = TRUE))
  install.packages("BiocManager")
BiocManager::install(version = "3.14")
BiocManager::install("GEOquery")
BiocManager::install("limma")
BiocManager::install("GOstats")
BiocManager::install("AnnotationDbi")
BiocManager::install("hgu133plus2.db")
BiocManager::install("clusterProfiler")

install.packages("pheatmap")
install.packages("msigdbr")
install.packages("ggnewscale")
install.packages("writexl")
install.packages("rlang")
install.packages("vctrs")

library(rlang)
library(Biobase)
library(GEOquery)
library(limma)
library(GOstats)
library(tidyverse)
library(hgu133plus2.db)
library(pheatmap)
library(ggrepel)
library(ggplot2)
library(msigdbr)
library(clusterProfiler)
library(org.Hs.eg.db)
library(enrichplot)
library(ggnewscale)
library(writexl)

#Load the dataset from the GEO database, select dataset from the list
gse <- 	getGEO("GSE26051")
gse <- gse[[1]] 

#assess the scale of the data. As the values are beyond 16, we need log2 transformation
exprs(gse) <- log2(exprs(gse))

#plot the distribution of each sample. If they are similar, it is reasonable to assume normalized data
boxplot(exprs(gse), outline = FALSE)

#inspect clinical variables
sampleInfo <- pData(gse)

#extract sample info------------------------------------------------------------
sampleInfo_selected <- select(sampleInfo, 
                              `tissue:ch1`,
                              `disease state:ch1`,
                              `age:ch1`,
                              geo_accession)

sampleInfo_selected <- rename(sampleInfo_selected, 
                              tendon = `tissue:ch1`,
                              disease_state = `disease state:ch1`,
                              age = `age:ch1`,
                              geo = geo_accession)

#pca and heatmap of samples-----------------------
corMatrix <- cor(exprs(gse),use="c")
pheatmap(corMatrix)

#check of the row name of the clinical samples match that of the correlation matrix (corMatrix)
rownames(sampleInfo)
colnames(corMatrix)

#check sample heatmap for clustering due to age, tendon type, and disease state
pheatmap(corMatrix,
         annotation_col=sampleInfo_selected)   

#create the principal component analysis and save it as pdf---------------------
pca <- prcomp(t(exprs(gse)))

pdf(file="PCA_Plot_HumanData_Age.pdf")
cbind(sampleInfo_selected, pca$x) %>% 
  ggplot(aes(x = PC1, y=PC2, col=disease_state,label=paste("tendon", geo))) + 
  geom_point() + 
  geom_text_repel() +
  theme(legend.position="none") +
  theme_bw() +
  theme_light() +
  theme_classic()
dev.off()

#identify and remove outliers based on having a tendon sheath
outliers <- c(2, 4, 9,14,18,25,27,32,37,41)

gse2 <- gse[,-outliers]


#pca and heatmap of samples w/o outliers----------------------------------------
pca2 <- prcomp(t(exprs(gse2)))

sampleInfo2 <- pData(gse2)

sampleInfo2_selected <- select(sampleInfo2, `tissue:ch1`,
                              `disease state:ch1`,
                              geo_accession)

sampleInfo2_selected <- rename(sampleInfo2_selected, tendon = `tissue:ch1`,
                              disease_state = `disease state:ch1`,
                              )  

corMatrix2 <- cor(exprs(gse2),use="c")

pheatmap(corMatrix2,
         annotation_col=sampleInfo2_selected,)

rownames(sampleInfo2)
colnames(corMatrix2)

pheatmap(corMatrix2,
         annotation_col=sampleInfo2_selected,
         filename = "Heatmap_HumanData_outliersremoved.pdf")

pdf(file="PCA_Plot_HumanData_unsheathedonly.pdf")
cbind(sampleInfo2_selected, pca2$x) %>% 
  ggplot(aes(x = PC1, y=PC2, col=disease_state,label=paste("tendon", geo_accession))) + 
  geom_point() + 
  geom_text_repel() +
  theme(legend.position="none")
dev.off()

#differential gene expression analysis------------------------------------------

#prepare design matrix to allocate samples along the correct parameter
design <- model.matrix(~0+sampleInfo2_selected$disease_state)

#check sample allocation to lesional / non-lesional
design
colnames(design) <- c("lesional","nonlesional")

# create annotation to allow data interpretation
anno <- fData(gse2)

anno <- select(anno, 
               `Gene Symbol`, 
               `Gene Title`,
               `Gene Ontology Biological Process`,
               `Gene Ontology Cellular Component`,
               `Gene Ontology Biological Process`,
               `Gene Ontology Molecular Function`)

# averaging replicate probes
data <- exprs(gse2)
data <- as.data.frame(data)
data$ID <- row.names(data)

anno2 <- as.data.frame(select(anno, 
                              `Gene Symbol`,))
anno2$ID <- row.names(anno2)
mydata <- merge(anno2, data, by ="ID")

A <- limma::avereps(mydata[  ,3:ncol(mydata)], mydata$`Gene Symbol`)

# calculate median expression level, which we will then use as a cutoff value 
cutoff <- median(A)

# TRUE or FALSE for whether each gene is "expressed" in each sample according 
#to our previously defined cutoff
is_expressed <- A > cutoff

# Identify genes expressed in more than 2 samples
keep <- rowSums(is_expressed) > 2

# check how many genes are removed / retained.
table(keep)

# subset to just those genes expressed in > 2 samples
gse2_kept <- A[keep,]

# fit the model to the data using lmFit after averaging replicate probes 
# and calculating weights
aw <- arrayWeights(gse2_kept, design)
fit <- lmFit(gse2_kept, design, weights = aw)
head(fit$coefficients)

# define contrast based upon which the differential analysis is performed (lesion vs non-lesion)
contrasts <- makeContrasts(lesional - nonlesional, levels=design)
fit2 <- contrasts.fit(fit, contrasts)

# apply Bayes' step to get differential expression statistics / p-values
fit2 <- eBayes(fit2)
topTable(fit2)

# see how many genes are differentially expressed overall
decideTests(fit2)
table(decideTests(fit2))

full_results2 <- topTable(fit2, number=Inf)

# Export DEG
full_results2 <- tibble::rownames_to_column(full_results2,"ID")
DEG <- as.data.frame(full_results2)

write_xlsx(DEG, "path")
getwd()

# Volcano plot
ggplot(full_results2, aes(x = logFC, y=-log10(P.Value))) + 
  geom_point()

p_cutoff <- 0.05
fc_cutoff <- 1

genesUp <- full_results2[full_results2$P.Value < p_cutoff &
                           full_results2$logFC > fc_cutoff,]
nrow(genesUp)

genesDown <- full_results2[full_results2$P.Value < p_cutoff &
                             full_results2$logFC < -fc_cutoff,]
nrow(genesDown)

full_results2 %>% 
  mutate(Significant = P.Value < p_cutoff, abs(logFC) > fc_cutoff ) %>% 
  ggplot(aes(x = logFC, y = -log10(P.Value), col=Significant)) + 
  geom_point() +
  theme_bw()

topN <- 20

full_results2 %>% 
  mutate(Significant = P.Value < p_cutoff, abs(logFC) > fc_cutoff ) %>% 
  mutate(Rank = 1:n(), Label = ifelse(Rank < topN, ID,"")) %>% 
  ggplot(aes(x = logFC, y = -log10(P.Value), col=Significant,label=Label)) + geom_point() + geom_text_repel(col="black") +
  theme_bw()


# Plot volcano plot with Il-6 annotation
full_results2$genelabels <- ""
full_results2$genelabels <- ifelse(full_results2$ID == "IL6R" 
                             | full_results2$ID == "IL6"
                             | full_results2$ID == "IL6ST"
                             | full_results2$ID == "STAT3"
                             | full_results2$ID == "STAT1"
                             | full_results2$ID == "IL11"
                             | full_results2$ID == "IL11RA"
                             | full_results2$ID == "COL1A2"
                             | full_results2$ID == "COL18A1"
                             | full_results2$ID == "ADAM17"
                             | full_results2$ID == "ADAM10"
                             | full_results2$ID == "JAK1"
                             | full_results2$ID == "GAB2"
                             | full_results2$ID == "MAPK1"
                             | full_results2$ID == "MAPK3"
                             | full_results2$ID == "COL1A1"
                             | full_results2$ID == "COL3A1"
                             | full_results2$ID == "MMP9"
                             | full_results2$ID == "MMP3"
                             | full_results2$ID == "MMP13"
                             ,
                              TRUE,FALSE)
full_results2$genelabels[full_results2$genelabels == TRUE] <- full_results2$ID[full_results2$genelabels == TRUE]
full_results2$genelabels[full_results2$genelabels == FALSE] <- NA

pdf(file="Volcanoplot_HumanData_IL6plus_averaged.pdf")
full_results2 %>% 
  mutate(Significant = P.Value < p_cutoff, abs(logFC) > fc_cutoff ) %>% 
  ggplot(aes(x = logFC, y = -log10(P.Value), col=genelabels, label=genelabels)) +
  geom_point() +
  geom_text_repel() +
  geom_vline(xintercept=c(-1, 1), color = "black") +
  geom_hline(yintercept=-log10(0.05), color = "black") +
  theme_bw()
dev.off()

# Create a heatmap of the top 20 genes
ids_of_interest <- mutate(full_results2, Rank = 1:n()) %>% 
  filter(Rank < topN) %>% 
  pull(ID)

gene_names <- mutate(full_results2, Rank = 1:n()) %>% 
  filter(Rank < topN) %>% 
  pull(Gene.Symbol) 

gene_matrix <- full_results2[ids_of_interest,]

pheatmap(gene_matrix,
         label_row = gene_names, 
         scale="row")

# Generate heatmap for Il-6 related genes
my_genes <- c("IL6", "IL6R","IL6ST", "STAT3", "STAT1", "IL11", "IL11RA", "COL1A2", "COL3A1",
              "COL18A1", "ADAM17", "ADAM10", "JAK1", "GAB2", "MAPK1", "MAPK3", "MMP3", "MMP13", 
              "MMP9", "COL1A1")

ids_of_interest2 <-  filter(full_results2,ID %in% my_genes) %>% 
  pull(ID)

gene_matrix2 <- A[ids_of_interest2,]

sampleInfo_genes2 <- select(sampleInfo, `tissue:ch1`,
                                                 `disease state:ch1`)
sampleInfo$`age:ch1`

pheatmap(gene_matrix2,
         scale="row",
         annotation_col=sampleInfo_genes2,
         cluster_cols=FALSE,
         filename = "Heatmap_HumanData_IL6plus.pdf")

#Filter dataset for GO analysis-------------------------------------------------
filtered2 <- full_results2[full_results2$P.Value < 0.01,]
genes_to_test <- filtered2$ID

GO_results <- enrichGO(gene = genes_to_test, OrgDb = org.Hs.eg.db, 
                       keyType = "SYMBOL", ont = "BP", pAdjustMethod = "BH")

goplot(GO_results)
dotplot(GO_results, showCategory=10)

GO_results_simMat <- pairwise_termsim(GO_results, method = "JC", semData = NULL, showCategory = 150)
emapplot(GO_results_simMat)

cnetplot(GO_results_simMat, categorySize="pvalue", foldChange = genes_to_test)

GO_results2 <- as.data.frame(GO_results)
write_xlsx(GO_results2, "path")


#GSEA analysis------------------------------------------------------------------ 

#create the gene ranking
rankedGenes <- full_results2 %>%
  mutate(rank = logFC*-log10(P.Value)) %>%
  arrange(-rank) %>%
  pull(rank,ID)

#GSEA analysis w/ the msigdbr human hallmark sets
msigdbr_df <- msigdbr(species = "human", category ="H")
msigdbr_t2g = msigdbr_df %>% dplyr::distinct(gs_name, gene_symbol) %>% as.data.frame()

gseaRes <- GSEA(rankedGenes,
                  TERM2GENE = msigdbr_t2g,
                  pvalueCutoff = 0.05,
                  minGSSize = 15,
                  maxGSSize = 500,
                  eps=0)
  
gseaRes2 <- as.data.frame(gseaRes)
 
dotplot(gseaRes, x = "enrichmentScore", color = "pvalue", showCategory=20)
gseaplot2(gseaRes, geneSetID = "HALLMARK_IL6_JAK_STAT3_SIGNALING", title = "Il6-JAK_Stat")

#GSEA analysis w/ the msigdbr human cell types 
msigdbr_df_ct <- msigdbr(species = "human", category ="C8")
msigdbr_t2g_ct = msigdbr_df_ct %>% dplyr::distinct(gs_name, gene_symbol) %>% as.data.frame()

gseaRes_ct <- GSEA(rankedGenes,
                TERM2GENE = msigdbr_t2g_ct,
                pvalueCutoff = 0.05,
                minGSSize = 20,
                maxGSSize = 500,
                pAdjustMethod = "BH",
                eps=0)

gseaRes_ct2 <- as.data.frame(gseaRes_ct)

#Dotplot biased for fibroblasts
dotplot(gseaRes_ct, x = "enrichmentScore", color = "pvalue", showCategory=c("HU_FETAL_RETINA_FIBROBLAST",
                                                     "CUI_DEVELOPING_HEART_C3_FIBROBLAST_LIKE_CELL",
                                                     "GAUTAM_EYE_CORNEA_FIBROBLASTS",
                                                     "GAUTAM_EYE_CHOROID_SCLERA_FIBROBLASTS",
                                                     "TRAVAGLINI_LUNG_LIPOFIBROBLAST_CELL",
                                                     "DURANTE_ADULT_OLFACTORY_NEUROEPITHELIUM_FIBROBLASTS_STROMAL_CELLS",
                                                     "TRAVAGLINI_LUNG_ADVENTITIAL_FIBROBLAST_CELL",
                                                     "TRAVAGLINI_LUNG_ALVEOLAR_FIBROBLAST_CELL",
                                                     "GAUTAM_EYE_IRIS_CILIARY_BODY_MEG3_HIGH_FIBROBLASTS",
                                                     "TRAVAGLINI_LUNG_MYOFIBROBLAST_CELL"
                                                     ))

#GSEA analysis w/ the msigdbr human biocarta 
msigdbr_df_bc <- msigdbr(species = "human", category ="C2", subcategory = "BIOCARTA")
msigdbr_t2g_bc = msigdbr_df_bc %>% dplyr::distinct(gs_name, gene_symbol) %>% as.data.frame()

gseaRes_biocarta <- GSEA(rankedGenes,
                   TERM2GENE = msigdbr_t2g_bc,
                   pvalueCutoff = 1,
                   minGSSize = 15,
                   maxGSSize = 500,
                   eps = 0)

gseaRes_biocarta2 <- as.data.frame(gseaRes_biocarta)
dotplot(gseaRes_biocarta, x = "enrichmentScore", color = "pvalue", showCategory=10)
gseaplot2(gseaRes_biocarta, geneSetID = "BIOCARTA_IL6_PATHWAY", title = "Il6_Pathway")

#GSEA analysis w/ the msigdbr human PID 
msigdbr_df_PID <- msigdbr(species = "human", category ="C2", subcategory = "PID")
msigdbr_t2g_PID = msigdbr_df_PID %>% dplyr::distinct(gs_name, gene_symbol) %>% as.data.frame()

gseaRes_PID <- GSEA(rankedGenes,
                         TERM2GENE = msigdbr_t2g_PID,
                         pvalueCutoff = 1.00, 
                         minGSSize = 15,
                         maxGSSize = 500,
                         eps = 0)

gseaRes_PID2 <- as.data.frame(gseaRes_PID)
dotplot(gseaRes_PID, color = "pvalue", showCategory=10)
gseaplot2(gseaRes_PID, geneSetID = "PID_IL6_7_PATHWAY", title = "PID_IL6_7_PATHWAY")


#GSEA analysis w/ the msigdbr human reactome
msigdbr_df_React <- msigdbr(species = "human", category ="C2", subcategory = "REACTOME")
msigdbr_t2g_React = msigdbr_df_React %>% dplyr::distinct(gs_name, gene_symbol) %>% as.data.frame()

gseaRes_React <- GSEA(rankedGenes,
                    TERM2GENE = msigdbr_t2g_React,
                    pvalueCutoff = 1.00, 
                    minGSSize = 15,
                    maxGSSize = 500,
                    eps = 0)

gseaRes_React2 <- as.data.frame(gseaRes_React)
dotplot(gseaRes_React, color = "pvalue", showCategory=10)
gseaplot2(gseaRes_React, geneSetID = "REACTOME_IL_6_TYPE_CYTOKINE_RECEPTOR_LIGAND_INTERACTIONS", 
                                title = "REACTOME_IL_6_TYPE_CYTOKINE_RECEPTOR_LIGAND_INTERACTIONS")

#GSEA analysis w/ the msigdbr human WIKIPATHWAYS
msigdbr_df_Wiki <- msigdbr(species = "human", category ="C2", subcategory = "WIKIPATHWAYS")
msigdbr_t2g_Wiki  = msigdbr_df_Wiki  %>% dplyr::distinct(gs_name, gene_symbol) %>% as.data.frame()

gseaRes_Wiki  <- GSEA(rankedGenes,
                      TERM2GENE = msigdbr_t2g_Wiki,
                      #pvalueCutoff = 0.05,
                      pvalueCutoff = 1.00, # to retrieve whole output
                      minGSSize = 15,
                      maxGSSize = 500,
                      eps = 0)

gseaRes_Wiki2 <- as.data.frame(gseaRes_Wiki)
dotplot(gseaRes_Wiki, color = "pvalue", showCategory=10)
gseaplot2(gseaRes_Wiki, geneSetID = "WP_IL6_SIGNALING_PATHWAY", title = "WP_IL6_SIGNALING_PATHWAY")

#Biased GO analysis to human data
my_pathways <- c("connective tissue development", "extracellular matrix organization", "extracellular structure organization", "external encapsulating structure organization", 
                 "response to hypoxia", "response to decreased oxygen levels", "wound healing", "glucose metabolic process",
                 "response to oxygen levels", "cellular response to hypoxia", "cellular response to decreased oxygen levels", "Wnt signaling pathway",	
                 "cellular response to oxygen levels", "negative regulation of cell migration", "epithelial cell migration", "epithelium migration",
                 "tissue migration", "endothelial cell migration", "tissue remodeling", "ameboidal-type cell migration",
                 "regulation of chemotaxis", "regulation of response to wounding", "glucose homeostasis", "collagen metabolic process",
                 "regulation of epithelial cell migration", "epithelial cell proliferation", "regulation of wound healing", "positive regulation of chemotaxis",	
                 "interleukin-6 production", "regulation of cell growth", "ERK1 and ERK2 cascade", "muscle cell proliferation",
                 "extracellular matrix assembly", "cell chemotaxis", "positive regulation of epithelial cell migration", "positive regulation of ERK1 and ERK2 cascade",
                 "smooth muscle cell migration", "regulation of ERK1 and ERK2 cascade", "collagen fibril organization", "muscle cell migration",
                 "smooth muscle cell proliferation", "positive chemotaxis", "sprouting angiogenesis", "regulation of peptidase activity", "regulation of endopeptidase activity"
)

datplot <- filter(GO_results2, Description %in% my_pathways)

datplot <- datplot %>% 
  separate(GeneRatio, into = c("pathwayGenes", "totalGenes"), sep = "/") %>% 
  mutate(pathwayGenes = as.numeric(pathwayGenes),
         totalGenes = as.numeric(totalGenes)) %>% 
  mutate(geneRatio = pathwayGenes / totalGenes) %>%
  select(-pathwayGenes, -totalGenes)

datplot <- datplot %>%
  mutate(Description = fct_reorder(Description, geneRatio)) %>%
  filter(pvalue <= 0.05)

plot_dot <- ggplot(datplot, aes(x = geneRatio,
                                y = Description,
                                size = Count,
                                color = -p.adjust))
plot_dot +
  geom_point() +
  labs(x = 'GeneRatio', 
       y = '') +
  scale_color_gradient(low = "black", high = "red") +
  theme_bw()
```
