## SupplementaryMaterial15to19_AnalysisCode for "IL-6 SIGNALING EXACERBATES HALLMARKS OF CHRONIC TENDON DISEASE BY STIMULATING REPARATIVE FIBROBLASTS": SupplementaryMaterial16_HistologicalAnalysisHuman_ImageJ.ijm

```
// Author: Patrick Jager;
// Fiji / ImageJ version: 2.1.0 (2021-02-09)
// Requires BioVoxxel Toolbox (https://imagej.net/BioVoxxel_Toolbox)
// How to install BioVoxxel Toolbox: 
//   Help > Update... (> (restart Fiji > Update...) > Manage update sites >
//    > Check box in front of "BioVoxxel" > Cancel > Apply changes

#@ File (label = "Input directory", style = "directory") input
#@ File (label = "Output directory", style = "directory") output

///////////////
//   MAIN    //
///////////////

import_image(input);
full_img_name = File.name;
img_name = File.nameWithoutExtension;

set_center();
close(full_img_name);

split_channels();

find_centroids("DAPI");

create_blob_mask(20);

measure_channel("CD90");
measure_channel("IL6R");

close("CD90");
close("IL6R");

save_table("res_CD90");
save_table("res_IL6R");

close("*");


///////////////
// FUNCTIONS //
///////////////

function import_image(img_path) { 
// Opens an image using the Bio-Formats extension.
	run("Bio-Formats Macro Extensions");
  run("Bio-Formats Importer", "open=[" + img_path +"] " + 
  "color_mode=Default rois_import=[ROI manager] view=Hyperstack stack_order=XYCZT");
}


function set_center() { 
  // Creates a duplicate of the image, merges channels, adjusts contrasts,
  // asks for user input to select a point and returns an array with 
  // the x- and y-coordinates of the selected point.
  run("Duplicate...", "title=center_image duplicate");
  
  run("Z Project...", "projection=[Max Intensity]");
	
  run("Make Composite");
  Stack.setChannel(1);
  run("Enhance Contrast", "saturated=0.35");
  Stack.setChannel(2);
  run("Enhance Contrast", "saturated=0.35");
  Stack.setChannel(3);
  run("Enhance Contrast", "saturated=0.35");

  //setTool("point");
  //waitForUser("Do it!", "Please select the center.");
  //Roi.getCoordinates(xpoints, ypoints);
  //_center = newArray(2);
  //run("Coordinates...", "x=0 y=0");
  //run("Coordinates...", "x=" + _center[0] + " y=" + _center[1]);
  close("center_image");
  //close("MAX_center_image");
}

//split_channels();
function split_channels() { 
// Split the channels and rename the images.

  run("Split Channels");

  // Rename.
  for (i=1; i<=nImages; i++) {
    selectImage(i);
    //run("8-bit");
    image_title = getTitle();
    if (matches(image_title, "(.*C1.*)")) {
      rename("DAPI");
    } else if (matches(image_title, "(.*C2.*)")) {
      rename("IL6R");
    } else if (matches(image_title, "(.*C3.*)")) {
      rename("CD90");
    }
  }
}

//find_centroids("DAPI");

function find_centroids(img_title) { 
// Enhances the image and finds the coordinates of the 
// nuclei centroids. The "Results" that this function generates
// is will be used by "create_blob_mask".

  selectWindow(img_title);
  run("Duplicate...", " ");

  //waitForUser("ROI", "Select the correct region of interest using the polygon selection tool, and copy it to the other channels");
  //run("Clear Outside");

  //run("Pseudo flat field correction", "blurring=100 hide"); //Different tissue density cause distortions too. 
  run("Bandpass Filter...", "filter_large=20 filter_small=10 " +
      "suppress=None tolerance=5 autoscale saturate");
  waitForUser("ROI", "Check if all the ROI fit the image and contain real cell nuclei");    
  setAutoThreshold("Otsu dark");
  
  setOption("BlackBackground", true);
  
  run("Convert to Mask");

  //run("Open");
  run("Watershed");

  if (nResults > 0) run("Clear Results");
  run("Find Maxima...", "prominence=1 strict output=List");

  close("DAPI-1");
}

//create_blob_mask(10);

function create_blob_mask(diameter) { 
// Creates a circle around the supplied x and y coordinates
// supplied by the "Results" of the "find_centroids" function
// and creates a mask with blobs of the diameter specified
// with "diameter".
  
  radius = round(diameter/2);

	// Create mask template.
	getDimensions(width, height, channels, slices, frames);
	newImage("blob_mask", "8-bit black", width, height, 1);
	setColor(255);

	// Draw blobs.
	for (i = 0; i < nResults; i++) {
		x_temp = round(getResult("X", i)) - radius;
		y_temp = round(getResult("Y", i)) - radius;
		fillOval(x_temp, y_temp, diameter, diameter);
	}

	// Watershed blobs and add them to the ROI-manager.
	run("Watershed");
	if (roiManager("count") > 0) roiManager("Delete");
	run("Analyze Particles...", "add");
	close("blob_mask");
	waitForUser("ROI", "Check if all the ROI fit the image and contain real cell nuclei");

}


function measure_channel(img_title) { 
// Measure the contents of all the blobs that were identified
// with create blob mask in the MTT channel.
// Requires output from get_center().

  selectWindow(img_title);
  waitForUser("ROI", "Check if the channel measured ("+ img_title +") has the correct ROI overlay");

  //run("8-bit");
  //run("Duplicate...", " ");

//run("Pseudo flat field correction", "blurring=50 hide");
//run("Gaussian Blur...", "sigma=4");
//setThreshold(0, 155);
//setOption("BlackBackground", true);
//run("Convert to Mask");
  //waitForUser;

//run("Set Measurements...", "mean modal integrated median display redirect=None decimal=1");
run("Set Measurements...", "mean modal centroid integrated median display redirect=None decimal=1");
roiManager("multi-measure measure_all");

//close(img_title + "-1");
Table.rename("Results", "res_" + img_title);
}

function save_table(table) {
  // Saves a table as .csv-file. 
  selectWindow(table);
  Table.save(output + File.separator + img_name + "_" + table + "_" + ".csv");
  close(table);
}
```
