## SupplementaryMaterial15to19_AnalysisCode for "IL-6 SIGNALING EXACERBATES HALLMARKS OF CHRONIC TENDON DISEASE BY STIMULATING REPARATIVE FIBROBLASTS": SupplementaryMaterial17_HistologicalAnalysisHuman_R.R

```
# Author: Patrick Jager;
# R version 4.0.5 (2020-11-23)
library(tidyverse)  # Version 1.4.0
library(ggrepel)
# Import data -----------------------------------------------------------------

setwd("path")
getwd()

dat <- tibble(paths = list.files(full.names = T),
              data = map(paths, read_csv)) %>% 
  mutate(paths = basename(paths)) %>% 
  separate(col = paths,
           into = c('sample', 'condition',  'trash1', 'trash2', 'trash3', 'channel'),
           sep = '_', extra = 'drop') %>% 
  select(-trash1, -trash2, -trash3) %>% 
  rowid_to_column('img_id')

dat <- unnest(dat, cols = 'data') %>% 
  select(img_id, sample, channel, condition, Mean, Mode, Median, X, Y) %>% 
  rename(mean = Mean, mode = Mode, median = Median, x = X, y = Y)

dat$sample <- paste(dat$sample, dat$condition, sep="_")

# Plot Histogram for the SecABCtrl and the samples to set thresholds------------
ggplot(dat, aes(mean, color = channel)) +
  geom_freqpoly()

dat_ABCtrl <- dat[dat$condition == "SecABCtrl",]

ggplot(dat_ABCtrl, aes(mean, color = channel)) +
  geom_freqpoly(binwidth = 10)
ggplot(dat_ABCtrl, aes(mean, color = channel)) +
  geom_density() 

mean(dat_ABCtrl[dat_ABCtrl$channel == "IL6R",]$mean)
mean(dat_ABCtrl[dat_ABCtrl$channel == "CD90",]$mean)

max(dat_ABCtrl[dat_ABCtrl$channel == "IL6R",]$mean)
max(dat_ABCtrl[dat_ABCtrl$channel == "CD90",]$mean)

ggplot(dat, aes(mean, color = channel)) +
  geom_density() +
  scale_x_continuous(expand = c(0,0), limits = c(0,240), breaks=seq(0,240,20))

mean(dat[dat$channel == "IL6R",]$mean)
mean(dat[dat$channel == "CD90",]$mean)

max(dat[dat$channel == "IL6R",]$mean)
max(dat[dat$channel == "CD90",]$mean)

dat_bizeps <- dat[dat$condition == "Bizeps",]
ggplot(dat_bizeps, aes(mean, color = channel)) +
  geom_density() +
  scale_x_continuous(expand = c(0,0), limits = c(0,100), breaks=seq(0,100,20))

dat_knee <- dat[dat$condition == "Knee",]
ggplot(dat_knee, aes(mean, color = channel)) +
  geom_density() +
  scale_x_continuous(expand = c(0,0), limits = c(0,100), breaks=seq(0,100,20))

# Apply threshold based on plots to find the positive cells for each channel----
dat_CD90 <- dat %>% 
  filter(channel == 'CD90',
         mean > 20)

dat_IL6R <- dat %>% 
  filter(channel == 'IL6R',
         mean > 18)

# Find the double positive cells------------------------------------------------
dat_CD90_double <- dat_CD90 %>%
  select(-mean, -mode, -median, -img_id)

dat_IL6R_double <- dat_IL6R %>%
  select(-mean, -mode, -median, -img_id)

double <- inner_join(dat_CD90_double, dat_IL6R_double,
                     by = c("sample", "condition","x", "y"))

# Count single and double positive cells----------------------------------------
n_total <- dat %>% 
  group_by(sample, condition) %>% 
  summarise(n = n()/2)

n_CD90 <- dat_CD90 %>% 
  group_by(sample, condition) %>% 
  summarise(CD90 = n())

n_IL6R <- dat_IL6R %>% 
  group_by(sample, condition) %>% 
  summarise(IL6R = n())

n_all <- full_join(full_join(n_total, n_CD90), n_IL6R)

n_double <- double %>% 
  group_by(sample, condition) %>% 
  summarise(n = n())

colnames(n_double) <- c("sample", "condition", "double_pos")

n_all <- full_join(n_all, n_double)

# replace NA with 0 and calculate percentages-----------------------------------
n_all[is.na(n_all)] <- 0

print <- n_all %>%
  mutate(perc_CD90_pos = CD90 / n * 100,
         perc_IL6R_pos = IL6R / n * 100,
         perc_double_pos = double_pos / n * 100,
         perc_double_of_IL6R = double_pos / IL6R * 100,
         perc_double_of_CD90 = double_pos / CD90 * 100
  )

print[is.na(print)] <- 0

# Plot percentages
ggplot(print, aes(x=condition, y=perc_CD90_pos)) +
  geom_boxplot() +
  geom_point() +
  geom_text_repel(aes(label = sample))

ggplot(print, aes(x=condition, y=perc_IL6R_pos)) +
  geom_boxplot() +  
  geom_point() +
  geom_text_repel(aes(label = sample))

ggplot(print, aes(x=condition, y=perc_double_pos)) +
  geom_boxplot() +  
  geom_point() +
  geom_text_repel(aes(label = sample))

ggplot(print, aes(x=condition, y=perc_double_of_IL6R)) +
  geom_boxplot() +  
  geom_point() +
  geom_text_repel(aes(label = sample))

ggplot(print, aes(x=condition, y=perc_double_of_CD90)) +
  geom_boxplot() +  
  geom_point() +
  geom_text_repel(aes(label = sample))

#Collect and export statistical data
median_IQR <- matrix(ncol = 5, nrow = 4)
colnames(median_IQR) <- c("perc_CD90_pos", 
                          "perc_IL6R_pos", 
                          "perc_double_pos", 
                          "perc_double_of_IL6R", 
                          "perc_double_of_CD90")
rownames(median_IQR) <- c("bizeps_median",
                          "bizeps_IQR",
                          "knee_median",
                          "knee_IQR")

median_IQR[1,1] <- median(print[print$condition == "Bizeps",]$perc_CD90_pos)
median_IQR[2,1] <- IQR(print[print$condition == "Bizeps",]$perc_CD90_pos)
median_IQR[3,1] <- median(print[print$condition == "Knee",]$perc_CD90_pos)
median_IQR[4,1] <- IQR(print[print$condition == "Knee",]$perc_CD90_pos)

median_IQR[1,2] <- median(print[print$condition == "Bizeps",]$perc_IL6R_pos)
median_IQR[2,2] <- IQR(print[print$condition == "Bizeps",]$perc_IL6R_pos)
median_IQR[3,2] <- median(print[print$condition == "Knee",]$perc_IL6R_pos)
median_IQR[4,2] <- IQR(print[print$condition == "Knee",]$perc_IL6R_pos)

median_IQR[1,3] <- median(print[print$condition == "Bizeps",]$perc_double_pos)
median_IQR[2,3] <- IQR(print[print$condition == "Bizeps",]$perc_double_pos)
median_IQR[3,3] <- median(print[print$condition == "Knee",]$perc_double_pos)
median_IQR[4,3] <- IQR(print[print$condition == "Knee",]$perc_double_pos)

median_IQR[1,4] <- median(print[print$condition == "Bizeps",]$perc_double_of_IL6R)
median_IQR[2,4] <- IQR(print[print$condition == "Bizeps",]$perc_double_of_IL6R)
median_IQR[3,4] <- median(print[print$condition == "Knee",]$perc_double_of_IL6R)
median_IQR[4,4] <- IQR(print[print$condition == "Knee",]$perc_double_of_IL6R)

median_IQR[1,5] <- median(print[print$condition == "Bizeps",]$perc_double_of_CD90)
median_IQR[2,5] <- IQR(print[print$condition == "Bizeps",]$perc_double_of_CD90)
median_IQR[3,5] <- median(print[print$condition == "Knee",]$perc_double_of_CD90)
median_IQR[4,5] <- IQR(print[print$condition == "Knee",]$perc_double_of_CD90)

write.table(median_IQR, file = "IL6R_CD90.txt", sep = ";", row.names = TRUE, col.names = TRUE)

#Statistical analysis knee vs bizeps
bizeps_knee <- print[print$condition != "SecABCtrl",]

CD90_bizeps_knee <- t.test(perc_CD90_pos ~ condition, data=bizeps_knee, alternative = "greater")
print(CD90_bizeps_knee)

CD90_bizeps_knee <- wilcox.test(perc_CD90_pos ~ condition, data=bizeps_knee, alternative = "greater")
print(CD90_bizeps_knee)

IL6R_bizeps_knee <- t.test(perc_IL6R_pos ~ condition, data=bizeps_knee, alternative = "greater")
print(IL6R_bizeps_knee)

IL6R_bizeps_knee <- wilcox.test(perc_IL6R_pos ~ condition, data=bizeps_knee, alternative = "greater")
print(IL6R_bizeps_knee)

double_CD90_IL6R_bizeps_knee <- t.test(perc_double_pos ~ condition, data=bizeps_knee, alternative = "greater")
print(double_CD90_IL6R_bizeps_knee)

perc_double_of_IL6R_bizeps_knee <- t.test(perc_double_of_IL6R ~ condition, data=bizeps_knee, alternative = "greater")
print(perc_double_of_IL6R_bizeps_knee)

perc_double_of_CD90_bizeps_knee <- t.test(perc_double_of_CD90 ~ condition, data=bizeps_knee, alternative = "greater")
print(perc_double_of_CD90_bizeps_knee)

capture.output(CD90_bizeps_knee, 
               IL6R_bizeps_knee, 
               double_CD90_IL6R_bizeps_knee, 
               perc_double_of_IL6R_bizeps_knee,
               perc_double_of_CD90_bizeps_knee,
               file="StatAnalysis_CD90_IL6R_InVivo_ttest.doc")


write.table(print, file = "IL6R_CD90_allSamples.txt", sep = ";", row.names = TRUE, col.names = TRUE)
```
