## SupplementaryMaterial15to19_AnalysisCode for "IL-6 SIGNALING EXACERBATES HALLMARKS OF CHRONIC TENDON DISEASE BY STIMULATING REPARATIVE FIBROBLASTS": SupplementaryMaterial19_HistologicalAnalysisAssembloid_R.R

```
# Author: Patrick Jager;
# R version 4.0.5 (2020-11-23)
library(tidyverse)  # Version 1.4.0

# Import data -----------------------------------------------------------------
setwd("path")
getwd()
dat <- tibble(paths = list.files(full.names = T),
              data = map(paths, read_csv)) %>% 
  mutate(paths = basename(paths)) %>% 
  separate(col = paths,
           into = c('condition', 'donor', 'sample', 'trash', 'channel'),
           sep = '_', extra = 'drop') %>% 
  select(-trash) %>% 
  # mutate(across(c('donor', 'slide', 'section'), ~as.integer(.))) %>% 
  rowid_to_column('img_id')

dat <- unnest(dat, cols = 'data') %>% 
  select(img_id, sample, donor, channel, condition, Mean, Mode, Median, X, Y) %>% 
  rename(mean = Mean, mode = Mode, median = Median, x = X, y = Y) %>% 
  mutate(r = abs(x)) %>% 
  select(-x, -y)


# Apply threshold --------------------------------------------------------------
dat_Scx <- dat %>% 
  filter(channel == 'Scx',
         mean > 650)

# Plot histograms --------------------------------------------------------------
ggplot(dat, aes(mean, color = channel)) +
  geom_freqpoly()

ggplot(dat, aes(mean, color = channel)) +
  geom_density() +
  scale_x_continuous(expand = c(0,0), limits = c(0,2000), breaks=seq(0,2000,50))


# Count ------------------------------------------------------------------------
n_total <- dat %>% 
  group_by(sample, donor, condition) %>% 
  summarise(n = n())

n_Scx <- dat_Scx %>% 
  group_by(sample, donor, condition) %>% 
  summarise(Scx = n())

n_all <- full_join(n_total, n_Scx)
n_all %>% print(n = nrow(.))


# Calculate percentages --------------------------------------------------------
print <- n_all %>% 
  mutate(perc_Scx_pos = Scx / n * 100)

ggplot(print, aes(x=condition, y=perc_Scx_pos)) +
  geom_boxplot() +
geom_violin(draw_quantiles = c(0.25, 0.5, 0.75))
  geom_point()
  
#StatisticalAnalysis of ScxGFP percentage
  fit_ScxPercentages <- aov(print$perc_Scx_pos ~ print$condition, 
                                      data = print)
  summary(fit_ScxPercentages)
  
  capture.output(summary(fit_ScxPercentages),
                 file="StatAnalysis_ScxPerc_Constructs_ANOVA_Th650.doc")
  
  tukey_ScxPercentages <- TukeyHSD(fit_ScxPercentages)
  
  tukey_ScxPercentages
  
  capture.output(tukey_ScxPercentages,
                 file="StatAnalysis_ScxPerc_Constructs_Tukey_Th650.txt")

# Staining gradient ----------------------------------------------------------
## Classify cells as positive or negative.
Scx_threshold <- 650
dat_Scx_classified <- dat %>% 
  filter(channel == 'Scx') %>% 
  mutate(staining = if_else(mean <= Scx_threshold,
                                  'negative', 'positive'))

## Count stuff.
get_counts_rel_3 <- function(x, .breaks) {
  # Basically a relative histogram that gets the percentage of live cells
  # in a given interval.
  # x: a dataframe
  # .breaks: number of breaks for the histogram
  
  # Get total histogram.
  hist_total <- x %>% 
    pluck('r') %>% 
    hist(plot = F, breaks = .breaks)
  
  # Copy breaks from total histogram so breaks of the grouped histograms
  # match.
  break_values <- hist_total$breaks
  
  # Get histogram of negative cells.
  hist_negative <- x %>% 
    filter(staining == 'negative') %>% 
    pluck('r') %>% 
    hist(plot = F, breaks = break_values)
  
  # Get histogram of positive cells.
  hist_positive <- x %>% 
    filter(staining == 'positive') %>% 
    pluck('r') %>% 
    hist(plot = F, breaks = break_values)
  
  # Calculate relative histogram.
  counts_i <- 
    tibble(breaks = hist_total$breaks[2:length(hist_total$breaks)-1],
           counts_total = hist_total$counts[1:length(hist_total$counts)],
           counts_negative  = hist_negative$counts[1:length(hist_negative$counts)],
           counts_positive = hist_positive$counts[1:length(hist_positive$counts)],
           counts_negative_rel = counts_negative /counts_total*100,
           counts_positive_rel = counts_positive /counts_total*100) %>%
    mutate(counts_negative_rel = if_else(is.na(counts_negative_rel), 
                                      0, counts_negative_rel),
           counts_positive_rel = if_else(is.na(counts_positive_rel), 
                                      0, counts_positive_rel)) %>%
    mutate(counts_positive_rel_cum = cumsum(counts_positive_rel)/sum(counts_positive_rel))
  return(counts_i)
}

dats <- full_join(dat_Scx_classified, dat_TPPP3_classified)

dat_counts_rel_2 <- 
  dats %>% 
  group_nest(img_id, channel, condition, donor) %>% 
  mutate(counts = map(data, get_counts_rel_3, .breaks = seq(0,1000,50))) %>% 
  select(-data) %>% 
  unnest(cols = c(counts))

dat_counts_rel_2_long <- dat_counts_rel_2 %>% 
  #select(-counts_total, -counts_negative, -counts_positive) %>% 
  # RELATIVE PLOT
  # rename(positive = 'counts_positive_rel', negative = 'counts_negative_rel') %>% 
  # CUMULATIVE PLOT:
  rename(positive = 'counts_positive_rel_cum', negative = 'counts_negative_rel') %>% 
  gather(key = 'staining', value = 'count', -img_id, -channel, -breaks, -condition, -donor)
  
dat_plot <- dat_counts_rel_2_long %>% 
  filter(staining == 'positive')

## Plot stuff.
ggplot(dat_plot, aes(breaks/1000, count, 
                     fill = condition, 
                     color = condition)) +
  #facet_grid(condition~.) +
  stat_summary(fun.data = mean_se, geom = 'ribbon', 
               alpha = 0.3, color = F, show.legend = F) +
  stat_summary(fun.data = mean_se, geom = 'point', cex = 2) +
  stat_summary(fun.data = mean_se, geom = 'line', cex = 1.0, show.legend = T) +
  labs(x = 'Distance to center [mm]', 
       y = 'Positive cells [%]') +
  theme_bw() 
```
